# Therapy-induced normal tissue damage promotes breast cancer metastasis

**DOI:** 10.1101/2020.10.17.343590

**Authors:** Douglas W. Perkins, Ivana Steiner, Syed Haider, David Robertson, Richard Buus, Lynda O’Leary, Clare M. Isacke

## Abstract

Disseminated tumour cells frequently exhibit a period of dormancy that renders them insensitive to targeting by chemotherapeutic agents, conversely the systemic delivery of chemotherapies can result in normal tissue damage. Using multiple mouse and human breast cancer models, we demonstrate that prior chemotherapy administration enhances metastatic colonisation and outgrowth. *In vitro*, chemotherapy treatment induces fibroblast senescence associated with a senescence associated secretory phenotype (SASP) that accelerates 3D tumour spheroid growth. These chemotherapy-treated fibroblasts, and their pro-tumourigenic function, can be effectively eliminated by targeting the anti-apoptotic protein BCL-xL. *In vivo*, chemotherapy treatment induces SASP expression in normal tissues, however the accumulation of senescent cells is limited and BCL-xL inhibitors are unable to reduce chemotherapy-enhanced metastasis. This likely reflects that chemotherapy-exposed normal tissues support metastatic colonisation via the secretion of pro-tumourigenic factors and remodelling of the extracellular matrix, but that damaged stromal cells do not enter a full BCL-xL-dependent senescence or switch their dependency to other anti-apoptotic BCL-2 family members. In summary, this study highlights the role of the metastatic microenvironment in controlling outgrowth of disseminated tumour cells and the need to identify novel therapeutic approaches to effectively limit the pro-tumourigenic effects of chemotherapy-induced normal tissue damage.

## Introduction

In breast cancer patients, prevention of local recurrence is achieved by surgery and localised radiotherapy, whilst adjuvant systemic chemotherapy or targeted agents are aimed at limiting the outgrowth of disseminated tumour cells (DTCs) at secondary sites. Oestrogen receptor-negative (ER^-^) and ER^+^ breast cancers have a different risk of recurrence profiles with the majority of ER^-^ tumours displaying an early (<5 years after primary diagnosis) relapse and ER^+^ tumours showing a broader profile with a higher incidence of late (>5 years) relapse reflecting the propensity of DTCs to reside in a prolonged period of dormancy prior to re-entering proliferation. Cytotoxic chemotherapy is most effective in targeting proliferating metastatic lesions while dormant DTCs are relatively chemoresistant. In both scenarios the ratio of tumour cells compared to surrounding secondary site tissue is low; hence the stromal milieu has a profound impact on the fate of disseminated tumour cells and disease outcome. For example, an ageing or fibrotic microenvironment has been shown to promote outgrowth of ER^+^ DTCs (Chandler et al., 2019; Turrell et al., 2023) whist normal tissue radiation exposure enhances metastatic outgrowth in ER^-^ models (Nolan et al., 2022). One feature of ageing tissues is the accumulation of senescent cells (Baker et al., 2016; Baker et al., 2008), which underpin a range of age-related pathological conditions including idiopathic pulmonary fibrosis, atherosclerosis, osteoarthritis and renal failure (Baker et al., 2016; Baker et al., 2011; Schafer et al., 2017). Further, multiple studies have revealed that increased normal tissue cell senescence can promote tumour progression (Alspach et al., 2014; Gonzalez-Meljem et al., 2018), creating a link between ageing and cancer (Krtolica et al., 2001).

Here we have investigated the effect of systemic chemotherapy on the induction of normal tissue damage, including the induction of cellular senescence, and the role of therapy-exposed stromal cells in shaping disease progression.

## Results

### Prior chemotherapy treatment promotes metastatic tumour growth

To test the hypothesis that systemic chemotherapy treatment can prime the host tissue microenvironment, making it a more receptive niche for metastatic tumour development, we employed experimental metastasis assays that discriminate between the effects of the chemotherapy treatment on the stroma from any direct effects on the tumour cells. Naïve BALB/c mice were treated with a course of chemotherapy; consisting of three or four doses over two weeks of doxorubicin combined with cyclophosphamide. This schedule reflects the cyclic delivery of chemotherapy in breast cancer patients in the clinic (Fisher et al., 2001; Fisher et al., 1990; Jones et al., 2006; Mainetti et al., 2013) and anti-tumour efficacy in syngeneic models (**Supplementary Figure 1**). Mice were then given a 7-10 day recovery period chosen to allow for clearance of the chemotherapeutic agents (Johansen, 1981; Park et al., 2012) and chemotherapy-induced apoptotic cells (Tilsed et al., 2022), and for the repopulation of circulating lymphocytes (Saida et al., 2015).

After the recovery period mice were inoculated intravenously with tumour cells. We chose mouse mammary tumour or human breast cancer lines which show limited metastatic outgrowth to model the scenario of non-proliferative disseminated cells in the secondary sites (Barkan et al., 2008; Naumov et al., 2002; Prunier et al., 2021). Initial experiments were performed with the D2.OR cell line which exhibits a dormant-like phenotype *in vivo* (Barkan et al., 2008; Montagner et al., 2020; Morris et al., 1994). 11 days after tumour cell intravenous inoculation into syngeneic BALB/c mice, single tumour cells were found scattered throughout the lung tissue of vehicle-treated mice, whereas discrete tumour lesions were readily detectable in the mice receiving prior chemotherapy (**Figure 1A-1B**). By 98 days after tumour cell inoculation, this translated into a significantly increased lung tumour burden in chemotherapy-treated mice (**Figure 1C**). To validate these findings, this experiment was repeated using the related D2A1 cell line (Montagner et al., 2020; Morris et al., 1994). Metastatic lesions were detectable in the vehicle-alone group 15 days after intravenous inoculation, however, the size of the lesions significantly increased in the mice receiving prior chemotherapy treatment indicating more efficient outgrowth (**Figure 1D**).

**Figure 1.**
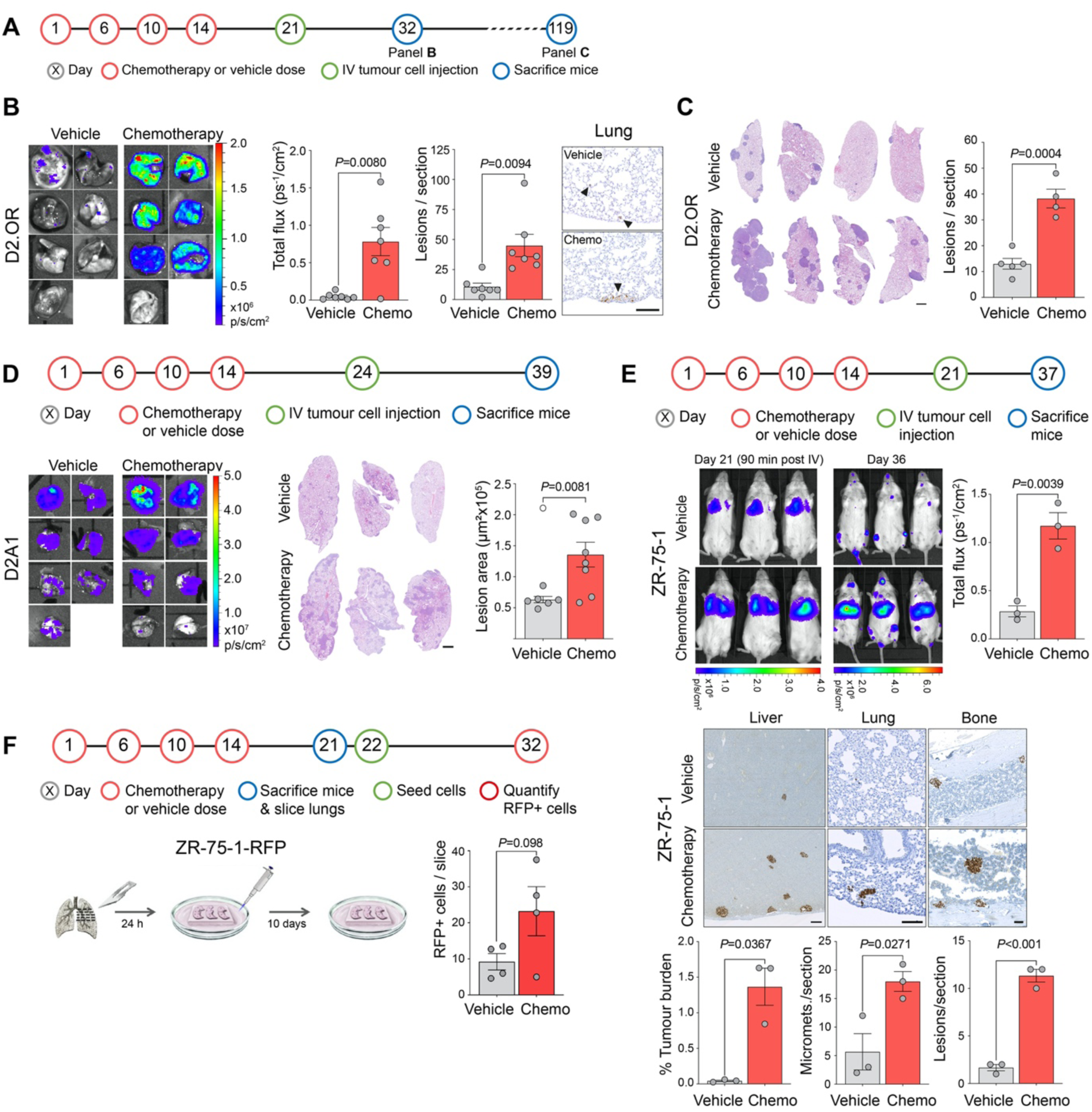
Prior chemotherapy treatment increases metastatic colonisation. **A** Schedule of chemotherapy (doxorubicin 2.7 mg kg-1 and cyclophosphamide 43 mg kg-1) or vehicle treatment and subsequent intravenous (IV) tumour cell inoculation. **B** BALB/c mice (n=7 per group) were treated with 4 doses of combination chemotherapy or vehicle as indicated. On Day 21 mice were inoculated via the tail vein with 1 x 10^6^ D2.OR-Luc tumour cells. Mice were sacrificed 11 days after inoculation (Day 32). Shown are *ex vivo* IVIS images of the lungs, quantification of IVIS signal in the lungs (mean ±SEM, Welch’s *t*-test), quantification of the mean number of metastatic lesions from 3 lung sections per mouse (mean ±SEM, Welch’s *t*-test), and representative immunohistochemical images of luciferase staining (scale bar, 125 µm). Black arrowheads indicate single disseminated tumour cells (vehicle), or macrometastatic deposits (chemotherapy). **C** BALB/c mice (n=4 or 5 per group) were treated with a schedule of chemotherapy or vehicle as outlined in panel A. On Day 21 mice were inoculated via the tail vein with 1 x 10^6^ D2.OR-Luc tumour cells. Mice were sacrificed 98 days after inoculation (Day 119). Shown are representative H&E stained sections of the lungs (scale bar, 1 mm) and quantification of the mean number of metastatic lesions from 3 lung sections per mouse (mean ±SEM, Student’s *t*-test). **D** BALB/c mice (n=7 or 8 per group) were treated as indicated. On Day 24 mice were inoculated via the tail vein with 4 x 10^5^ D2A1-Luc tumour cells. Mice were sacrificed 15 days after inoculation (Day 39). Shown are *ex vivo* IVIS images of the lungs, representative H&E stained lung sections (scale bar, 1 mm) and quantification of the mean size of the lung metastatic lesions in 2 lung sections per mouse (mean values per mouse ±SEM, Welch’s *t*-test). One mouse in the vehicle group was determined to be a statistical outlier, indicated by a hollow data point. **E** NOD Rag gamma (NRG) mice (n=3 per group) were treated with a schedule of chemotherapy or vehicle as illustrated in the timeline. Mice were implanted subcutaneously with a 0.36 mg oestrogen pellet on Day 17. On Day 21, 2 x 10^6^ ZR-75-1-Luc human breast tumour cells were inoculated via the tail vein and IVIS imaged after 90 minutes. 15 days after tumour cell inoculation (Day 36), mice were IVIS imaged *in vivo* (mean ±SEM, Student’s *t*-test), sacrificed 24 hours later (Day 37), and percentage of metastatic burden based on lamin A/C staining was quantified in 3 liver sections (mean ±SEM, Welch’s *t*-test; scale bar, 250 µm), 3 lung sections, and 2 bone sections (mean ±SEM, Student’s *t*-tests; scale bar, 100 µm) per mouse. **F** BALB/c mice (n=2 per group) were treated with chemotherapy or vehicle as indicated. On Day 21 mice were sacrificed, agarose-perfused lungs sliced and cultured on an inert matrix in M199 medium for 24 hours before 1,000 ZR-75-1-RFP human breast cancer cells were seeded onto the slices and incubated for a further 10 days. Two lung slices per mouse were imaged by confocal microscopy and the number of RFP^+^ tumour cells per slice quantified (±SEM, Student’s *t*-test).

Finally, the experimental approach was repeated using immunocompromised NOD RAG gamma (NRG) mice inoculated with human ER^+^ ZR-75-1 breast cancer cells, a cell line exhibiting a dormant phenotype in *in vivo* models (Gawrzak et al., 2018; Turrell et al., 2023). Again there was a striking enhancement of metastatic colonisation in the mice that had received prior chemotherapy. Whilst intravenously injected cells initially disseminated to the lungs, the majority of subsequent metastatic burden manifested in the liver (**Figure 1E**), with lung and bone metastasis also significantly increased. These effects were recapitulated in *ex vivo* assays with a significant increase in the number of viable ZR-75-1 cells surviving 10 days after seeding onto *ex vivo* organotypic lung slices derived from chemotherapy-treated, compared to vehicle-treated, mice (**Figure 1F**).

### Chemotherapy treatment causes normal tissue damage

Using freshly extracted RNA from lungs of mice sacrificed 7 days after the last chemotherapy treatment (Day 21) (**Figure 2A**), gene expression profiling was performed using the NanoString mouse PanCancer Immune and PanCancer Pathways panels (**Figure 2B, Supplementary Tables 1, 2**). By principal component analysis (PCA) the vehicle-treated mice clustered separately from the chemotherapy-treated mice, implying global differences across the 750 target gene panels. The significantly upregulated genes in the chemotherapy-treated samples clustered in pathways associated with pro-inflammation, apoptosis, cytokines and chemokines, and pro-cancer progression, indicating that chemotherapy treatment creates a favourable, tumour-promoting microenvironment. These include: the Il6 family cytokine *Lif*, which promotes cell proliferation via STAT3 signalling and has been reported to promote breast cancer metastasis (Li et al., 2014; Niwa et al., 2009); *Cxcl10*, reported to enhance breast cancer lung metastasis (Pein et al., 2020; Perrott et al., 2017); the pro-tumourigenic factor *Wnt5a* (Azazmeh et al., 2020; Kobayashi et al., 2018); *Ccl2* (MCP-1), a potent macrophage chemoattractant, reported to play a role in cancer progression (Acosta et al., 2013; Park et al., 2012; Sanoff et al., 2014), and the related factors *Ccl7* (MCP-3), *Ccl8* (MCP-2), *Ccl12* (MCP-5), *Cd14* and *Cx3cr1* (Ingersoll et al., 2010; Marshall et al., 2020; Proudfoot, 2002; Rajaram et al., 2013). In contrast, downregulated genes clustered primarily in humoural immunity pathways such as the B-cell surface antigens *Cd79b*, *Ms4a1*, and *Cd19* and the B-cell lineage specific *Pax5*. Analysis of the immune cell abundance using the NanoString Immune data confirmed a significantly reduced abundance in B cells and an increased abundance of dendritic cells in chemotherapy-treated lungs (**Supplementary Figure 2A**). This analysis also identified a small but significant decrease in T cell abundance but no differences in any T cell subsets. Similarly, immunohistochemical staining of lungs taken between 1 and 27 days after the last dose of chemotherapy revealed no notable changes in the number of CD8- or CD4-positve cells (**Supplementary Figure 2B**).

**Figure 2.**
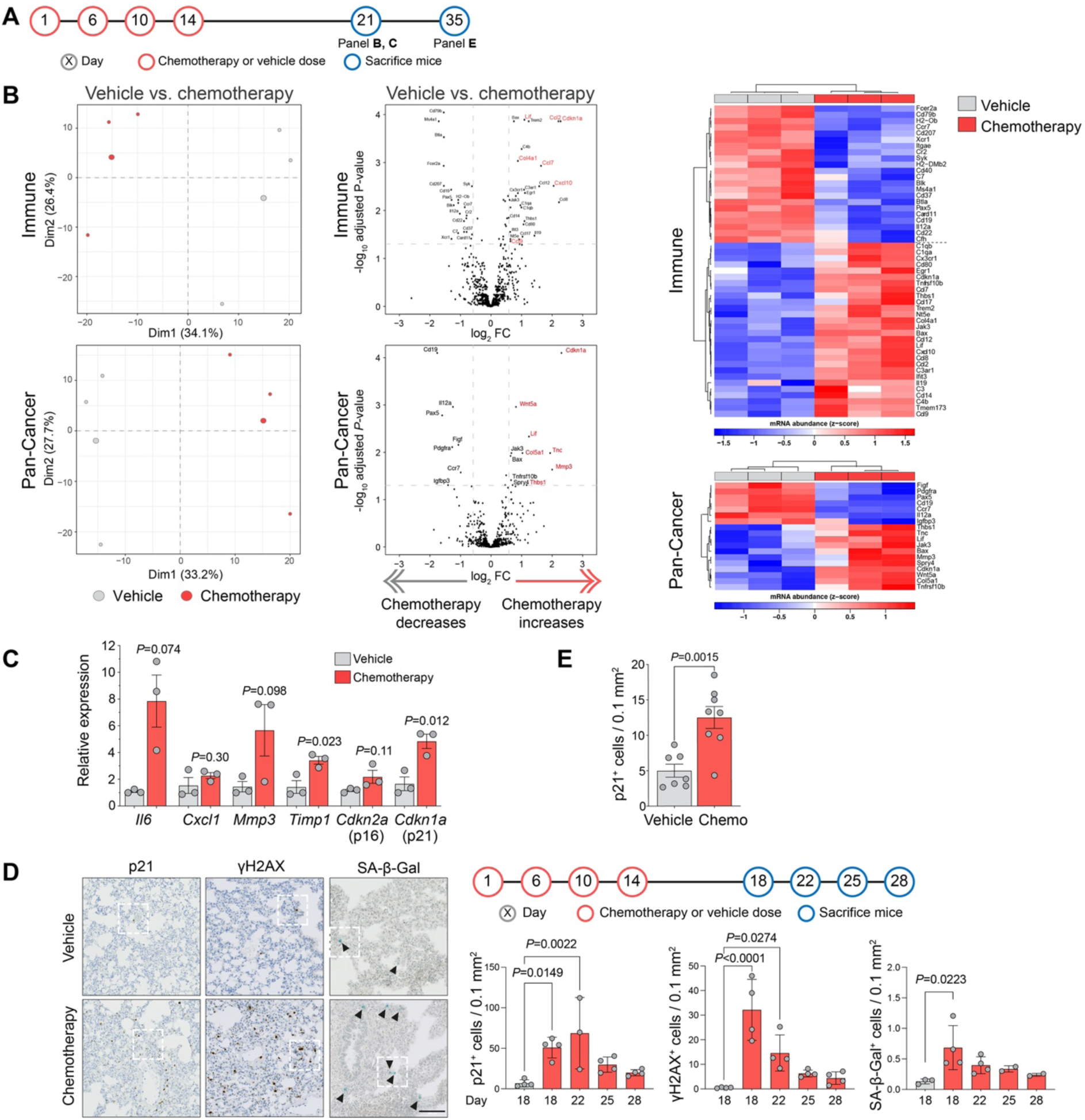
Characterisation of the chemotherapy-treated mouse lung. **A** Schedule of the chemotherapy or vehicle treatment and subsequent tissue collection timepoints. **B** BALB/c mice (n=3 per group) were treated with 4 doses of chemotherapy or vehicle as illustrated in panel A. On Day 21 mice were sacrificed and the lungs snap-frozen prior to RNA extraction. Gene expression profiling using the NanoString mouse PanCancer Immune (upper) or PanCancer Pathways (lower) panels. Left panels, principal component analysis; middle panels, volcano plot showing differentially expressed (DE) genes in chemotherapy *vs*. vehicle treated groups. Genes with absolute log_2_ fold change ≥ 0.585 and FDR adjusted *p-*value < 0.05 were considered significant and shown in the heat map (right panels). Genes highlighted in red are discussed in the text. **C**. Expression of the selected DE genes indicated in panel B measured by RTqPCR (mean ±SEM, Student’s *t*-test except *Il6* Welch’s *t*-test). **D** BALB/c mice (n=4 per group) were treated with 4 doses chemotherapy or vehicle as illustrated in the timeline, and sacrificed on Days 18, 22, 25 or 28 (n=4 per group). Lungs were removed and right lobes were fixed in formalin for immunohistochemistry, while left lobe was embedded in OCT and snap frozen for SA-β-Gal staining. Left panel, representative images of p21 (Abcam) and γH2AX staining, and SA-β-Gal activity (Day 18, vehicle; Day 22, chemo; scale bar, 100 µm). Arrowheads indicate blue SA-β-Gal^+^ cells. Right panels, quantification of staining (mean ±SD, one-way ANOVA). Higher power images of indicated areas are shown in Supplementary Figure 2C. **E** BALB/c mice (n=7 or 8 per group) were treated as illustrated in panel A and sacrificed on Day 35. p21 staining (Dako) in lung tissue was quantified (mean values ±SEM, Student’s *t*-test).

It has been reported that normal tissue senescence is elevated following systemic chemotherapy treatment (Demaria et al., 2017). Despite being long-lived and non-proliferative, senescent cells are far from static and inert. Rather, they are highly metabolically active and have elevated expression of the senescence-associated secretory phenotype (SASP), a diverse cocktail of factors responsible for many of their non-cell-autonomous functions (Coppé et al., 2008; Coppé Jean-Philippe and Judith, 2010; Schmitt et al., 2022). In both NanoString panels, a top hit within the significantly upregulated genes in chemotherapy-treated mice was *Cdkn1a* (p21) a marker of DNA damage-induced cell cycle arrest and cellular senescence (de Carné Trécesson et al., 2011; Jurk et al., 2012; Lawless et al., 2010) along with elevated expression of SASP genes such as *Mmp3, Cdkn2a, Il6* and *Cxcl1* and *Cxcl10*, and upregulation of *Ifit3* gene implicated in inducing p21 expression (Xiao et al., 2006). RTqPCR analysis confirmed a significantly increased expression of canonical SASP factors in the lung tissue of the chemotherapy-treated mice (**Figure 2C**) while staining of lung sections revealed elevated numbers of p21^+^ cells in chemotherapy treated lungs taken from mice sacrificed between Day 18 and 28 (**Figure 2D; Supplementary Figure 2C**). Similarly, chemotherapy induced activation of the DNA damage response (DDR), manifested by the phosphorylation of DNA damage mediator proteins such as γH2AX, was evident in the mouse lungs after 18 and 22 days (**Figure 2D**). In an independent experiment where mice were sacrificed 21 days following chemotherapy treatment (Day 35) an elevated number of p21^+^ cells were detected persisting in the lungs (**Figure 2E)**. Finally, assessing fresh-frozen sections for senescence-associated β-galactosidase (SA-β-Gal) activity as alternative marker of cellular senescence, revealed an elevated number of SA-β-Gal^+^ cells in chemotherapy-treated lungs (**Figure 2D; Supplementary Figure 2C**). However, it is notable that a much smaller number of SA-β-Gal^+^ cells were detected compared to p21^+^ cells, indicating either that SA-β-Gal^+^ cells are cleared within the 7 days after chemotherapy treatment and/or that the tolerable chemotherapy regime used *in vivo* is insufficient to drive stromal cells from a SASP-positive state into full senescence.

### Chemotherapy-treated fibroblasts promote tumour cell growth

Amongst the increased levels of SASP and pro-tumourigenic factors in the gene expression analysis were extracellular matrix remodelling and fibrosis genes (**Figure 2B**), including collagens type Va1 (*Col5a1*) and type IVa1 (*Col4a1*), *Mmp3, Thbs1* (thrombospondin1), and *Tnc* (tenascin C), profiles associated with activated fibroblasts (Walraven and Hinz, 2018). Tissue fibrosis is associated with increased tumourigenicity in multiple cancer types (Barkan et al., 2010; Cox and Erler, 2014; Turrell et al., 2023).

To elucidate whether damaged/activated fibroblasts are present in non-tumour bearing chemotherapy-treated mice, lung sections from vehicle or chemotherapy-treated mice were co-stained for p21 alongside cell-type markers for myofibroblasts (alpha-smooth muscle actin, αSMA), epithelial cells (EpCAM) and endothelial cells (endomucin) (**Figure 3A**). Consistent with the data presented in Figure 2, an increased number of p21^+^ cells were observed in lung tissue of the chemotherapy-treated mice (**Figure 3A**, middle panel). Since p21^+^ cells in the lungs of vehicle-treated mice are scarce, very few double-positive cells of any type were identified. In contrast, in the chemotherapy-treated lung tissue, p21^+^ cells were readily identified with the greatest increase being in double-positive p21 αSMA stained cells (**Figure 3A**, right panel). The identification of p21^+^ αSMA^+^ cells in chemotherapy-treated lung tissue was validated in immunohistochemical staining of lung sections from an independent experiment. By immunofluorescence and immunohistochemistry, the majority of αSMA staining in vehicle-treated lungs is localised to the vasculature, consistent with αSMA expression by pericytes and perivascular-localised fibroblasts. By contrast, in chemotherapy-treated lungs, p21 αSMA double positive cells are detected both associated with, and distant from, the vasculature, indicating that systemic chemotherapy drives the accumulation of damaged stromal cells in the lung tissue (**Figure 3A,B**).

**Figure 3.**
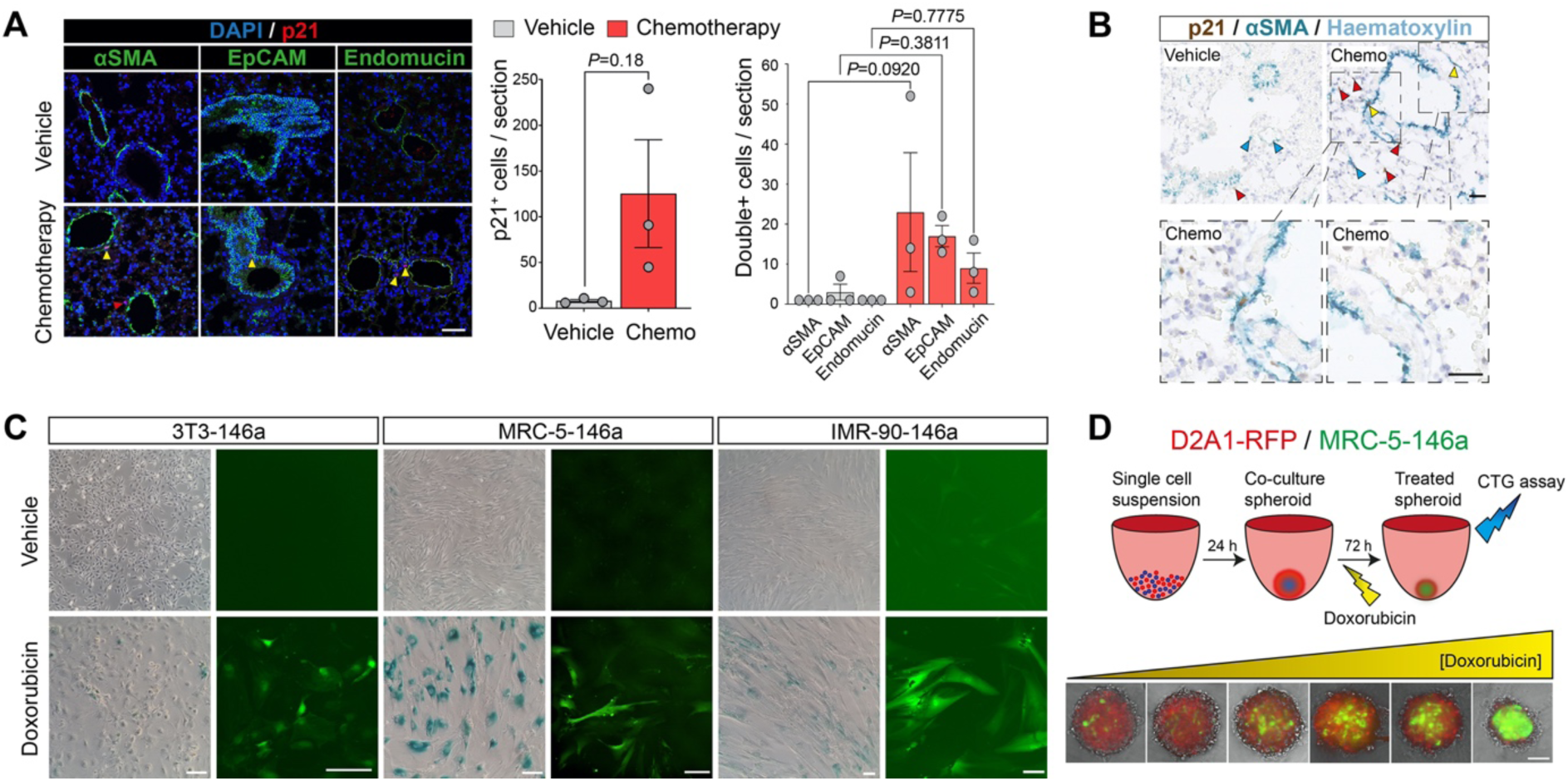
Fibroblast responses to chemotherapy treatment. **A** BALB/c mice (n=3 per group) were treated with 3 doses of chemotherapy or vehicle on Day 1, 5 and 9 and sacrificed on Day 16. FFPE lung sections were stained for p21 (red; Dako)) and DAPI (blue), and either α smooth muscle actin (αSMA), EpCAM or endomucin (green), scale bar, 50 µm. Red arrowheads indicate p21 single-positive cell, yellow arrowheads indicate double-positive cells. Staining was quantified by counting all p21^+^ cells per section (middle panel, mean ±SEM, Welch’s *t*-test), then counting double-positive cells (right panel, mean ±SEM, one way ANOVA). **B** Lung FFPE sections from BALB/c mice sacrificed 7 days following the final chemotherapy treatment or 1 day following the final vehicle treatment were stained for p21 (Dako, brown) and αSMA (cyan) and counterstained with haematoxylin. Blue arrowheads indicate αSMA single-positive cells, red arrowheads indicate p21 single-positive cells, yellow arrowheads indicate double positive cells. Left panels; scale bar, 50 µm. Lower panels; higher magnification views of indicated regions; scale bar, 25 µm. **C** Human lung MRC-5-146a fibroblasts, human lung IMR-90-146a fibroblasts and mouse 3T3-146a fibroblasts were treated with 170 nM doxorubicin for 24 hours, then drugs were removed. 20 (3T3-146a) or 21 (MRC-5-146a and IMR-90-146a) days later parallel wells were fixed and stained for SA-β-Gal, or expression of the GFP senescence reporter was visualised under a fluorescence microscope (scale bars, 200 µm). Vehicle-treated fibroblasts were seeded 3 days prior to staining or imaging alongside chemotherapy-treated fibroblasts. **D** 1,000 D2A1-RFP mouse tumour cells were co-seeded with 1,000 MRC-5-146a human fibroblasts into ultra-low attachment U-bottom 96-well plates to form co-culture spheroids. After 24 hours spheroids were treated with doxorubicin (0 - 850 nM). 72 hours later, spheroids were imaged (scale bar, 100 µm.)

To investigate the response of fibroblasts to chemotherapy treatment *in vitro*, primary human or established mouse fibroblasts were treated for 24 hours with doxorubicin, followed by culture in the absence of doxorubicin, resulting in a robust cell cycle arrest and accumulation of SA-β-Gal^+^ cells (**Figure 3C**). In an independent approach, fibroblasts were transfected with a senescence reporter construct pmiR146a-GFP (Kang et al., 2015b), in which GFP expression is driven by the senescence-associated microRNA146a promoter. Doxorubicin treatment resulted in increased expression of the senescence reporter, which is maintained for up to 20 days following drug withdrawal (**Figure 3C**). To model this in more complex *in vitro* models, RFP-tagged D2A1 mouse mammary tumour cells were admixed with human miR146a-transfected MRC-5 fibroblasts and co-seeded into ultra-low attachment U-bottom 96-well plates to form 3D co-culture spheroids (**Figure 3D**). Treatment with increasing concentrations of doxorubicin resulted in effective elimination of the tumour cells, whilst the admixed fibroblasts were induced to express markers of senescence and persist until the end of the assay.

Chemotherapy-treated primary human lung fibroblasts and mouse fibroblast lines show a robust upregulated expression of canonical SASP transcripts (**Figure 4A**), accompanied by increased secretion of SASP factors into the culture medium (**Figure 4B**). Incubation of mouse or human tumour spheroids with conditioned medium harvested from doxorubicin-treated human and mouse fibroblasts resulted in a striking increase in spheroid size compared to incubation with conditioned medium from control fibroblasts (**Figure 4C**). To confirm that the increased growth of the spheroids resulted from increased tumour cell proliferation as opposed to decreased cell death, two approaches were taken. First, tumour spheroids were fixed and embedded, and sections were stained for the proliferation marker Ki67 (**Figure 4D**). As the experiments were conducted in low (2%) serum conditions, few if any Ki67^+^ cells were detected in spheroids cultured with control conditioned medium. By contrast, spheroids incubated with chemotherapy-treated fibroblast conditioned medium were larger in size and had a ring of Ki67^+^ cells around their periphery. Second, tumour cell colony formation assays showed no significant difference in colony number in the presence of chemotherapy-treated fibroblast conditioned medium, but a notable increase in colony size (**Figure 4E**). To investigate the mechanism underlying this increased 3D tumour spheroid proliferation, D2A1 mouse or ZR-75-1 human tumour spheroids were treated with conditioned medium for 30 minutes, lysed and then assayed by phosphoprotein array. Increased phosphorylation of several key mitogenic factors was observed in the spheroids incubated with doxorubicin-treated fibroblast conditioned medium, in particular increased phosphorylation of Tyr701 STAT1, Tyr705 STAT3 and pan-SRC in the D2A1 spheroids and increased phosphorylation of Ser473 AKT, Thr202/Tyr204 ERK p42/p44 and Tyr705 STAT3 in the ZR-75-1 spheroids (**Figure 4F**). These findings are consistent with the activation of STAT3 signalling downstream of SASP cytokines such as IL6 and LIF (Hodge et al., 2005; Li et al., 2014; Niwa et al., 2009), whose expression is upregulated in chemotherapy-treated fibroblasts, promoting proliferation of mouse and human tumour cells in both *in vitro* and *in vivo* models.

**Figure 4.**
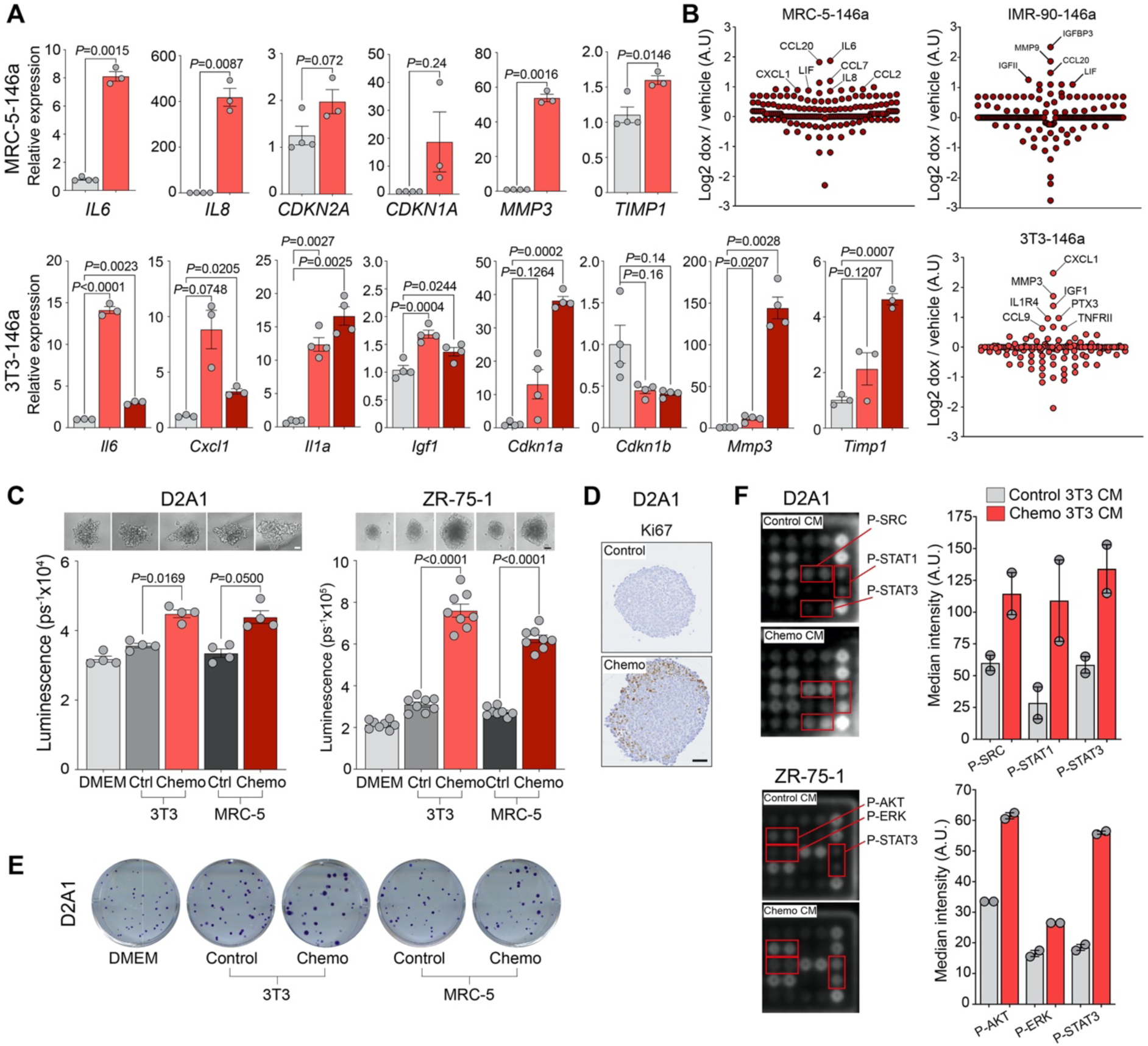
*In vitro* characterisation of chemotherapy-treated fibroblasts. **A** Human MRC-5-146a or mouse 3T3-146a fibroblasts were treated with 170 nM doxorubicin or vehicle for 24 hours. At the indicated time point after drug withdrawal RNA was extracted for RTqPCR analysis (mean ±SEM, Student’s *t*-test (MRC-5-146a) or one-way ANOVA (3T3-146a) for individual genes). **B** Human MRC-5-146a, human IMR-90-146a or mouse 3T3-146a fibroblasts were treated with doxorubicin or vehicle as described in panel A. Serum-free CM collected 20 days after doxorubicin withdrawal or from vehicle-treated fibroblasts was analysed using G2000 cytokine arrays. **C** 2,000 ZR-75-1 or D2A1 tumour cells per well were seeded into ultra-low attachment U-bottom 96-well plates in DMEM plus 2% FBS. After 24 hours, spheroids were incubated with DMEM supplemented with 2% FBS or CM from control or chemotherapy-treated 3T3-146a mouse fibroblasts or MRC-5-146a human fibroblasts supplemented with 2% FBS. After 4 days (ZR-75-1; n=4 spheroids per condition) or 6 days (D2A1; n=8 spheroids per condition), spheroid viability was measured using the CellTiter-Glo assay (mean ±SEM, one-way ANOVA; one-way ANOVA with Dunnett’s correction for multiple comparisons - ZR-75-1). Equivalent results were obtained in 2 (ZR-75-1) or >3 (D2A1) independent experiments. Representative phase contrast images of spheroids are shown above (scale bar, 100 µm). **D** D2A1 spheroids were treated with control or chemotherapy- treated 3T3-146a CM as described in panel C and collected after 4 days incubation. FFPE sections from embedded spheroids were stained for Ki67 (scale bar, 50 μm). **E** 100 D2A1 tumour cells per well were seeded into 6-well plates in DMEM plus 2% FBS or in CM from control or chemotherapy-treated 3T3-146a mouse fibroblasts or MRC-5-146a human fibroblasts supplemented with 2% FBS. 14 days later, plates were stained with crystal violet. **F** 12,500 ZR-75-1 or D2A1 tumours cells per well were seeded into U-bottom 96-well plates in DMEM plus 2% FBS and incubated for 48 hours. Spheroids were serum-starved for 1 hour, and then incubated for 30 minutes at 37°C with serum-free control or chemotherapy-treated 3T3-146a fibroblast CM for 30 minutes. Protein was extracted and analysed on phosphoprotein antibody arrays slides; ZR-75-1, 112.5 µg total protein per sample; D2A1, 30 µg total protein per sample. Median intensity of each target antibody spot was quantified. Shown are cropped images of the relevant portion of the array. Bar graphs show mean of the two replicate target antibody spots (mean ±SEM).

### Chemotherapy-treated fibroblasts can be selectively eliminated *in vitro*

Prolonged survival of senescent cells requires the avoidance of apoptotic-mediated cell death, commonly by upregulating expression of anti-apoptotic BCL-2 family members (de Carné Trécesson et al., 2011; Malaquin et al., 2020). BCL-2 family proteins bind to and inhibit BH3- initiator proteins such as BAD, which would otherwise activate the apoptotic effector protein BAX to trigger caspase activation and mitochondrial outer membrane permeabilisation (Willis et al., 2003). Consequently, agents which target the BCL-2 family members, such as BCL-2, BCL-xL, BCL-w and MCL-1 (**Figure 5A**), have been assessed for their efficacy in triggering apoptosis. In particular, navitoclax, which targets BCL-2, BCL-xL and BCL-w, has been shown to selectively kill senescent cells *in vivo* (Chang et al., 2016; Pan et al., 2017b; Saleh et al., 2020; Shahbandi et al., 2020; Troiani et al., 2022) and has been evaluated in human clinical trials for ageing/senescence related indications. However, there has been little focus on their use in targeting therapy-induced senescent fibroblasts.

**Figure 5.**
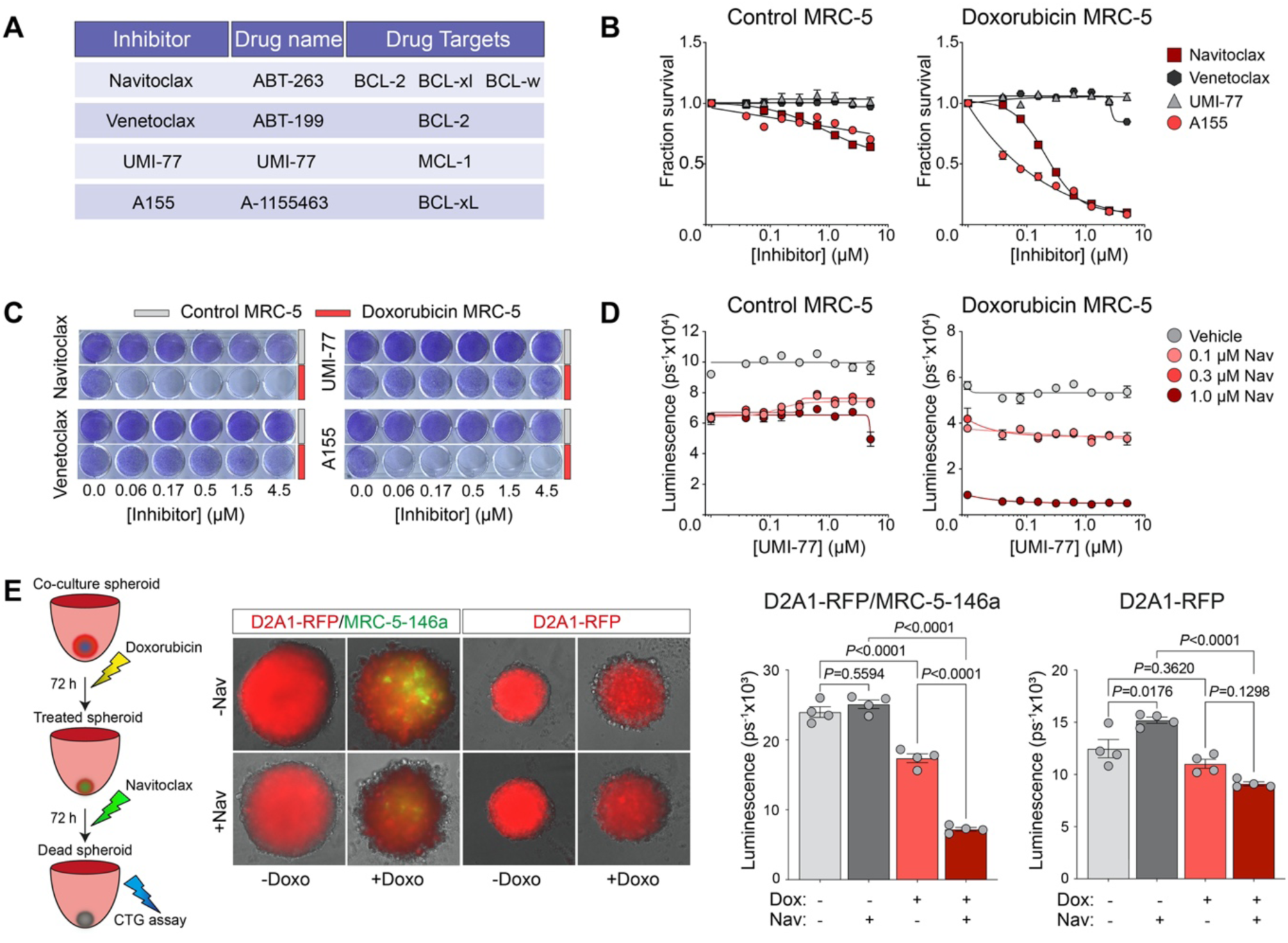
Navitoclax eliminates chemotherapy-induced senescent fibroblasts *in vitro*. **A** Details of the BCL-2 family inhibitors used in the study. **B** MRC-5-146a fibroblasts were treated with 1.7 µM doxorubicin for 24 hours. 9 days after treatment withdrawal, control or doxorubicin-treated fibroblasts were plated into 96-well plates (see methods; n=3 wells per condition). After 24 hours fibroblasts were treated with inhibitors as indicated and incubated for 72 hours before cell viability was measured by CellTiter-Glo. **C** Doxorubicin-treated MRC-5 fibroblasts, 10 days following treatment withdrawal, or control fibroblasts were seeded into 24-well plates (see methods; n=1 well per condition), then 24 hours later treated inhibitors as indicated. After 72 hours, plates were fixed and stained with crystal violet. **D** Doxorubicin-treated MRC-5 fibroblasts, 9 days following treatment withdrawal, or control fibroblasts were plated into 96- well plates (n=3 wells per condition). After 24 hours fibroblasts were treated with a range of concentrations of UMI-77, alone or in combination with 3 concentrations of navitoclax, and incubated for 72 hours before cell viability was measured by CellTiter-Glo assay. **E** 5,000 D2A1-RFP mouse tumour cells were seeded alone or co-seeded into ultra-low attachment U-bottom 96-well plates with 5,000 MRC-5-146a human fibroblasts (n=4 wells per condition). After 24 hours spheroids were treated with 170 nM doxorubicin or vehicle for 72 hours, followed by treatment with 4.5 µM navitoclax or vehicle for a further 72 hours. At the end of the assays spheroids were imaged (scale bar, 100 µm) and cell viability was quantified by CellTiter-Glo (one-way ANOVA). **B-E** Equivalent results were obtained in 2 - 4 independent experiments.

Fibroblasts treated with doxorubicin for 24 hours and then incubated for 10 days in the absence of drug were effectively killed by treatment with either navitoclax or the BCL-xL specific inhibitor A-1155463 (A115) whereas these inhibitors had limited effect on control vehicle-treated fibroblasts (**Figure 5B**). By contrast, treatment with either the BCL-2-only inhibitor venetoclax or the MCL-1 inhibitor UMI-77 had no impact on chemotherapy-treated or control fibroblast viability. Comparable results were obtained using human IMR-90 fibroblasts, mouse 3T3 fibroblasts and fibroblasts treated with docetaxel (**Supplementary Figure 3A-D**). A concern in these assays is that CellTiter-Glo monitors ATP levels in the cell and it is likely that non-proliferating cells have different basal ATP metabolism. Equivalent results were obtained using non-enzymatic crystal violet staining (**Figure 5C; Supplementary Figure 3A**). As MCL-1 exhibits some functional redundancy with other BLC2 family members (Eichhorn et al., 2014), the effect of combining the MCL-1 inhibitor UMI-77 with either navitoclax (**Figure 5D; Supplementary Figure. 3A**) or A115 (**Supplementary Figure 3D**) was examined. In combination, the addition of an MCL-1 inhibitor did not enhance killing of chemotherapy-treated fibroblasts *in vitro*. Finally, equivalent results were obtained treating fibroblasts with navitoclax formulated in corn oil (not shown), phosal or culture medium (**Supplementary Figure 3B and E)**.

To assess the ability of navitoclax to target chemotherapy-treated fibroblasts in a more complex environment, RFP-labelled D2A1 tumour cells alone or together with MRC-5- miR146a-GFP fibroblast were cultured as 3D spheroids and treated for 72 hours with vehicle or doxorubicin and then for a further 72 hours in the presence or absence of navitoclax (**Figure 5E**). As previously observed (**Figure 3D**), doxorubicin reduced viability of D2A1 cells *in vitro*, illustrated by the reduction in RFP expression within the spheroids. This effect was more evident in the co-culture spheroids and accompanied by a concomitant increase in expression of the fibroblast GFP senescence reporter. Subsequent navitoclax treatment had no effect on tumour cell only spheroid viability and similarly no impact on vehicle-treated co-culture 3D spheroids. In contrast, in chemotherapy-treated co-culture spheroids navitoclax treatment resulted in a further significant reduction in cell viability and a notable loss of GFP-expressing fibroblasts (**Figure 5E**).

### Combination chemotherapy and navitoclax treatment *in vivo*

Encouraged by the efficacy of navitoclax *in vitro*, we investigated whether these agents could reduce the impact of prior chemotherapy treatment on metastatic tumour outgrowth. BALB/c mice were treated with a course of chemotherapy or vehicle followed by 5 daily doses of navitoclax or vehicle and 3 days later inoculated intravenously with D2.OR tumour cells (**Figure 6A**). Consistent with previous data **(Figure 1A, B**), on Day 77 there was significantly increased tumour burden in the chemotherapy-treated mice compared to vehicle-treated mice, monitored by IVIS imaging, however, no significant difference between groups which had, or had not, received subsequent navitoclax treatment. On Day 91, when the first mice began to show signs of ill health, the experiment was terminated. In line with the Day 77 IVIS data, chemotherapy-treated mice had a significantly increased metastatic burden in the lungs, quantified by the numbers of metastatic lesions and percentage tumour area. Again, no significant effect of navitoclax treatment on vehicle-treated mice was observed. Equivalent results were obtained in NRG mice inoculated with human ZR-75-1 breast cancer cells (**Figure 6B**). As reported here (**Figure 1E**), mice pre-treated with a course of chemotherapy develop significantly more ZR-75-1 liver metastases than vehicle-treated mice (**Figure 6B**) but subsequent treatment with navitoclax did not reduce metastatic outgrowth.

**Figure 6.**
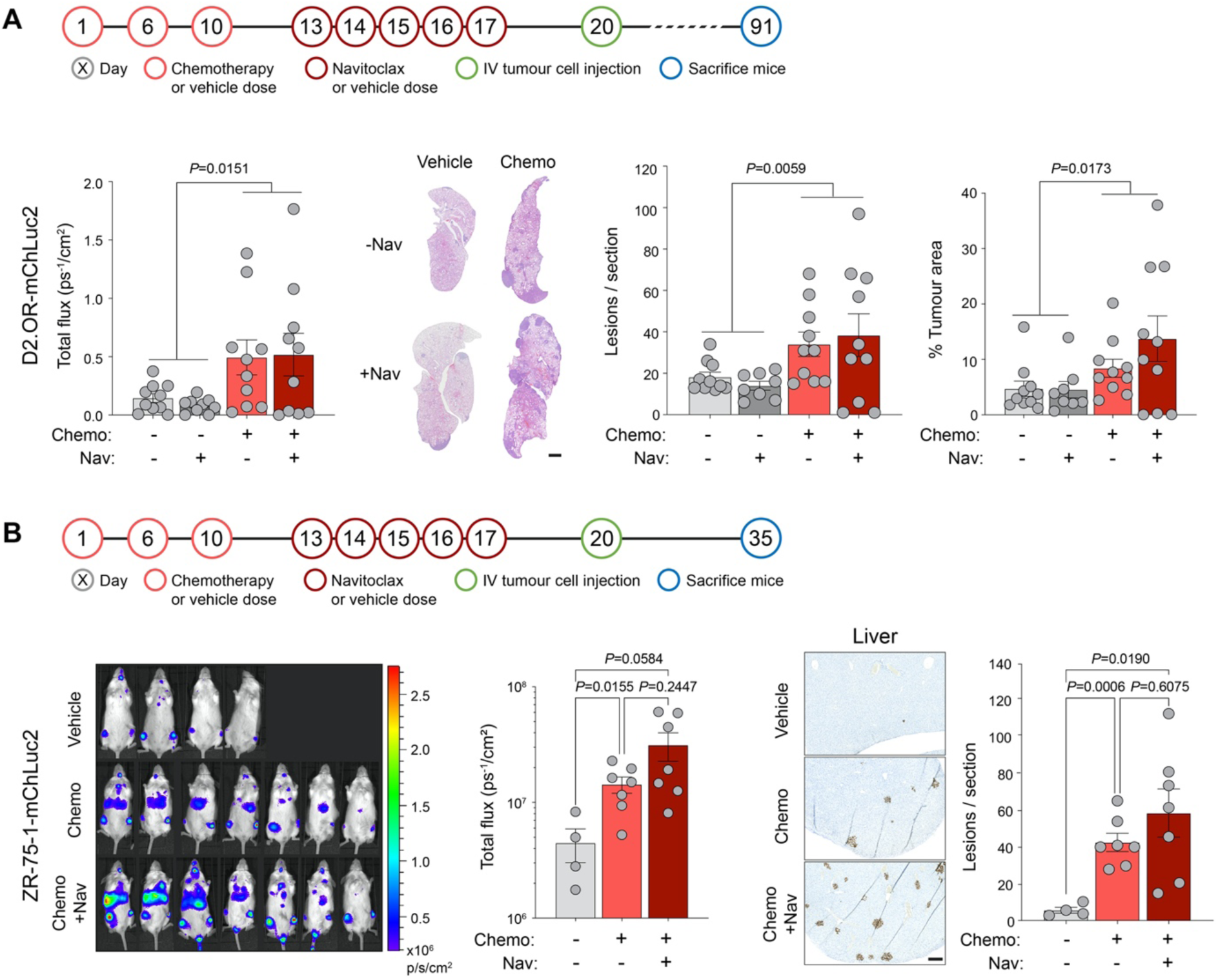
Effect of combined chemotherapy and navitoclax treatment on metastasis and survival. **A** Schedule for treatment of BALB/c mice with chemotherapy (Chemo) or vehicle followed by navitoclax (Nav) or vehicle, and subsequent intravenous (IV) inoculation of 1 x 10^6^ D2.OR-mChLuc2 tumour cells (n=9-10 mice per group). Quantification of *in vivo* IVIS imaging of the thoracic region on Day 77 (mean ±SEM, Mann-Whitney *U*-test between vehicle and chemotherapy groups combined). Quantification of lung metastatic burden at termination of the experiment (Day 91). Representative H&E lung sections (scale bar, 1 mm). Number of metastatic lesions per lung section. Percentage tumour burden in the lungs (mean ±SEM, Mann-Whitney *U*-tests between vehicle and chemotherapy groups combined). **B** Schedule for treatment of NOD Rag gamma mice and subsequent IV inoculation of 2 x 10^6^ ZR-75- 1-mChLuc2 tumour cells. *In vivo* IVIS quantification of mice (n=4 mice vehicle group, n=7 mice Chemo and Chemo + Nav groups) on Day 34 (±SEM, Welch’s ANOVA). Representative lamin A/C staining in the liver (scale bar, 250 µm) and quantification of metastatic lesions in the liver (mean ±SEM, Welch’s ANOVA).

To investigate this lack of efficacy in limiting metastatic outgrowth, naive BALB/c mice were treated with a course of vehicle or chemotherapy followed by 4 daily doses of vehicle or navitoclax, and 24 hours after the last dose (Day 24) lungs were profiled using the NanoString PanCancer Immune and Pathways panels (**Figure 7A)**. Lungs isolated from mice receiving chemotherapy alone showed, as previously reported (**Figure 2B**), a significantly upregulated SASP expression, including increased expression of *Il6*, *Lif*, *Cdkn1a* (p21), *Mmp3*, *Ccl2*, *Ccl7*, *Cxcl1* and *Cxcl10* (**Figure 7B; Supplementary Tables 1,2**), while PCA analysis revealed discrete clustering of the chemotherapy and vehicle groups (**Supplementary Figure 4A**). Of note, when analysing these data together with the data shown in Figure 2B, the vehicle and chemotherapy-treated samples from the independent experiments clustered together (**Supplementary Figure 4B**), indicating a reproducible effect of chemotherapy treatment on the lung tissue despite having different experimental endpoints (**Supplementary Tables 1,2**). Similarly, assessment of immune cell abundance scores again showed a decrease in B cells and significant increase in dendritic cells (**Supplementary Figure 5A**).

**Figure 7.**
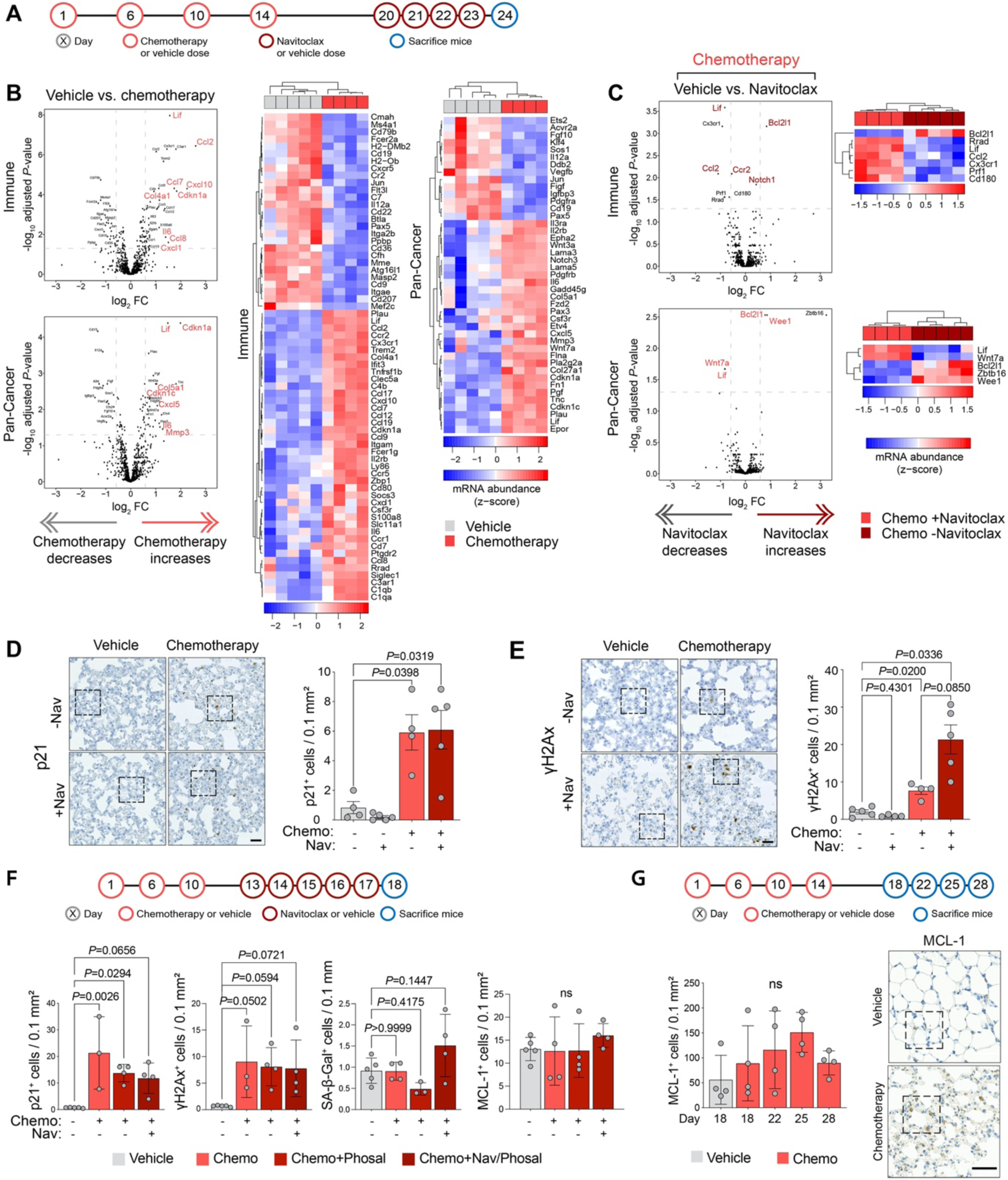
Navitoclax does not reverse chemotherapy-induced tissue damage *in vivo*. **A** Schedule for the treatment of BALB/c mice with chemotherapy (Chemo) or vehicle followed by navitoclax (Nav) or vehicle. **B-C** Mice were sacrificed on Day 24, RNA was extracted from snap-frozen lung tissue and gene expression profiling performed using the NanoString mouse PanCancer Immune and PanCancer Pathways panels. Normalisation was carried out as described in Figure 2B. Volcano plots showing differentially expressed genes between chemotherapy and vehicle treated mice (panel B), or between chemotherapy-treated mice subsequently treated with navitoclax or vehicle (panel C). Genes with absolute log_2_ fold change ≥ 0.585 and FDR adjusted *p-*value < 0.05 were considered significant and shown in the associated heat maps. Genes highlighted in red are discussed in the text. **D-E** Samples from panels B and C were stained for γH2Ax and p21 (Dako) (scale bars, 25 µm) and quantified (Welch’s ANOVA). Higher power images are shown in Supplementary Figure 5B and C. **F** Schedule for the treatment of BALB/c mice with chemotherapy or vehicle, followed by navitoclax or vehicle (Phosal). Mice were sacrificed one day after the final dose of navitoclax. Lung tissue was divided, prepared either for FFPE or snap-frozen embedded in OCT, and sections were stained for p21 (Abcam), γH2Ax, SA-β-Gal or MCL-1 and quantified (mean ±SD, one-way ANOVA). **G** Schedule for the treatment of BALB/c mice with chemotherapy or vehicle. Mice were sacrificed 4, 8, 11 and 14 days after the final dose of chemotherapy. Lung sections were stained for MCL-1. Quantification of MCL-1 staining on indicated days (mean ±SD, one-way ANOVA). Representative images (Day 18, vehicle; Day 22, chemotherapy; scale bar, 50 µm). Higher power images of indicated areas are shown in Supplementary Figure 5D.

Lung tissue from mice treated with navitoclax as a monotherapy showed no significant differentially expressed genes in either of the NanoString panels. In addition, consistent with the lack of efficacy in metastasis assays, there was little indication that navitoclax reverses the effects of chemotherapy treatment, with only 10 genes within the two panels showing a significant change in expression in mice receiving a subsequent course of navitoclax (**Figure 7C**) and no impact on the abundance of any immune cell populations (**Supplementary Figure 5A**). Importantly, within these chemotherapy-treated groups, while subsequent navitoclax administration significantly reduces the expression of *Lif* and *Ccl2* as well as the CCL2 receptor *Ccr2*, there was no significant reduction in expression of p21 (*Cdkn1a*). To confirm these findings, lung sections from the same mice were stained for p21. As previously reported (**Figure 2D**), few p21^+^ cells are detected in the lung tissue of vehicle-treated mice but their number significantly increased in lungs of chemotherapy-treated mice (**Figure 7D; Supplementary Figure 5B**). In chemotherapy-treated mice that received a subsequent course of navitoclax there was no change in the number of p21^+^ cells (**Figure 7D**) confirming that, and in contrast to the *in vitro* findings (**Figure 5**), navitoclax is unable to eliminate p21^+^ cells *in vivo*. Moreover, γH2Ax^+^ cells are readily detected in chemotherapy-treated mice (**Figure 7E; Supplementary Figure 5C**). Similar results were obtained in an independent experiment using different formulation of navitoclax delivery system (see Methods) and an alternative p21 antibody (**Figure 7F**). Finally, navitoclax treatment failed to deplete SA-β-Gal^+^ cells regardless of the formulation strategy. Although chemotherapy-treated fibroblasts showed no sensitivity to MCL-1 inhibition *in vitro* (**Figure 5B; Supplementary Figure 3A-D**) it remained a possibility, given its functional redundancy with other BLC-2 family members (Eichhorn et al., 2014), that upregulation of MCL-1 *in vivo* might promote escape from navitoclax-mediated killing. However, there was no significant change in MCL-1^+^ cell number following chemotherapy treatment, with or without a subsequent course of navitoclax (**Figure 7F,G; Supplementary Figure 5D**). By contrast, expression of *Bcl2l1*, which encodes BCL-xL was significantly upregulated in the chemotherapy plus navitoclax group, raising the prospect that upregulation of BCL-xL could provide an *in vivo* resistance mechanism allowing DNA-damaged senescent stromal cells to survive navitoclax treatment.

## Discussion

Here we show that in multiple independent models of both immune-competent and immune-compromised mice, pre-treatment with a clinically-relevant chemotherapy regimen of combined doxorubicin and cyclophosphamide followed by subsequent inoculation of tumour cells results in a significantly increased burden of metastatic disease. The use of the mouse D2.OR and human ER^+^ ZR-75-1 tumour cell lines which, when inoculated into naive mice, given rise to dormant DTCs at secondary sites has clinical relevance in the context of ER^+^ breast cancers, which frequently exhibit a late relapse phenotype developing metastatic disease years or decades after original diagnosis and treatment (Dittmer, 2017; Pan et al., 2017a; Pedersen et al., 2022; Ribelles et al., 2013) In both the mouse models and patients, DTCs have a prolonged exposure to the stromal microenvironment, which would be rendered a more pro-metastatic environment in patients receiving systemic chemotherapy as part of their treatment. In addition, although a late relapse phenotype is much less common in hormone receptor-negative breast cancers and these cancers frequently respond well to chemotherapy, there remains a significant proportion of patients with chemoresistant residual disease. Consequently, in both ER^+^ and ER^-^ scenarios, the data presented here highlight the potential negative consequences of chemotherapy treatment in which chemotherapy-mediated damage in normal tissue, paradoxically, has the potential to increase the risk of recurrence.

Here we demonstrate that systemic delivery of chemotherapy causes DNA damage in normal mouse tissues and increased accumulation of cells expressing SASP factors and staining positive for the senescence markers p21 and SA-β-Gal. The clinical importance of targeting both naturally-occurring and therapy-induced senescent cells has become increasingly clear (Childs et al., 2017; Kirkland and Tchkonia, 2020). Seminal proof of principle experiments, involving the genetic elimination of cells expressing the senescence marker p16^Ink4a^, have demonstrated an alleviation of age and therapy-induced pathologies (Baker et al., 2016; Baker et al., 2011), including progression and metastasis in a PyMT-MMTV spontaneous mammary carcinoma model (Demaria et al., 2017). Experimental studies into the therapeutic targeting of senescent cells has focused on administration of the BLC-2/BCL-xL/BCL-w inhibitor navitoclax (ABT-263), the BCL-2/BCL-xL inhibitor ABT-737 or the non-related combination of dasatinib and quercetin (D&Q) which have been shown to eliminate or reduce the number of senescent cells in ageing mouse models and alleviative the ageing phenotype (Chang et al., 2016; Xu et al., 2018; Yosef et al., 2016).

In our studies, D&Q was not selective for senescent fibroblasts *in vitro* and had no effect *in vivo* (data not shown). By contrast, navitoclax or the BCL-xL specific inhibitor A115 selectively eliminated chemotherapy-induced senescent fibroblasts *in vitro* and impaired their ability to promote tumour cell proliferation. However, using dosing regimens shown to be effective in ageing mouse models (Chang et al., 2016), navitoclax treatment failed to eliminate chemotherapy-induced p21^+^ cells *in vivo* and thus did not reduce metastatic outgrowth in different chemotherapy-treated mouse models. Whilst gene expression profiling provided some evidence of on-target effects of navitoclax in chemotherapy-treated mice, the number and magnitude of the changes were small. As the number of SA-β-Gal^+^ cells was substantially lower than the number of p21^+^ cells (**Figure 2D**; **Figure 7F**), this likely reflects that a course of systemic chemotherapy treatment results in the accumulation of damaged SASP-producing fibroblasts and other stromal cells sufficient to create a pro-tumourigenic metastatic niche but insufficient to induce a full senescent phenotype sensitive to BCL-xL inhibition. In addition, mice receiving the combination treatment showed an increased expression of *Bcl2l1* (encoding the navitoclax target BC-LxL) raising the possibility that, as has been observed in models of non-Hodgkin’s lymphoma (Merino et al., 2012), increased BCL-xL levels may contribute to BCL-2/BCL-xL inhibitor resistance. Of note, MCL-1 exhibits some functional redundancy with other BLC-2 family members (Eichhorn et al., 2014) however chemotherapy treatment did not significantly increase the number of MCL-1^+^ cells in the lung.

Whilst our 3D *in vitro* models support the hypothesis that increased SASP signalling by stromal fibroblasts directly promotes tumour growth by increasing mitogenic signalling and tumour cell proliferation; there are other consequences that chemotherapy-exposed cells could inflict on the microenvironment, contributing indirectly to metastatic colonisation. For example, the SASP can exhibit immune suppressive properties via the promotion of type-2 immune polarisation (Ruhland et al., 2016), resulting in reduced immune surveillance. Indeed, our cytokine array data reveal that amongst the most abundant chemokines secreted by chemotherapy-treated fibroblasts are macrophage inflammatory protein 1γ (MIP-1γ, CCL-9) and macrophage chemoattractant protein 1 (MCP-1, CCL-2), which is reflected in increased expression of *Ccl9* and *Ccl2* in the chemotherapy-treated lung tissue. Interestingly, NanoString profiling of chemotherapy-treated lungs revealed a decrease in B cells and increase in dendritic cells. However, a profound pro-tumourigenic effect of chemotherapy treatment is observed in NRG mice which lack a functional immune system, including having defective dendritic cells. By contrast, notable in our studies was the increased expression of extracellular matrix remodelling and fibrosis genes in chemotherapy-treated lungs, features associated with ageing tissues and increased metastatic outgrowth in multiple cancer models (Barkan et al., 2010; Cox and Erler, 2014; Turrell et al., 2023).

In summary, this study highlights the role of the metastatic microenvironment in controlling outgrowth of disseminated tumour cells and the need to identify novel therapeutic approaches to effectively limit chemotherapy-induced normal tissue damage and its pro-tumourigenic effects.

## Methods

### Reagents and cells

RTqPCR probes are listed in **Supplementary Table 3**. Antibodies, and the dilutions used, are detailed in **Supplementary Table 4**. MRC-5 and IMR-90 fibroblasts were purchased from ATCC in 2009 and 2016, respectively. NIH-3T3 and ZR-75-1 cells were from Isacke laboratory stocks. D2A1 and D2.OR cells were from Ann Chambers’ laboratory stocks (Morris et al., 1994). The generation of the metastatic D2A1-m2 subline has been described previously (Jungwirth et al., 2018). All cells were used within ten passages after resuscitation and were routinely subjected to mycoplasma testing. ZR-75-1 cells were short tandem repeat tested every 4 months (Stem Elite ID System, Promega). Doxorubicin (S1208), docetaxel (S1148), navitoclax (S1001), A-1155463 (S7800), venetoclax (S8048) and UMI-77(S7531) were purchased from Selleckchem. Cyclophosphamide monohydrate was purchased from Sigma-Aldrich (C0768).

#### Lentiviral transfection of tumour cells with RFP, luciferase and mCherry

As indicated, tumour cells were transduced with lentiviral expression particles containing either: (1) a firefly luciferase gene with a blasticidin-resistance gene (cells denoted -Luc) (Amsbio, LVP326); (2) a firefly luciferase gene, with an mCherry encoding gene (cells denoted -mChLuc2) (Genescript, Sall-IRES-Luc2-XbalXhol_PGK-H2BmCherry); or (3) an RFP encoding gene (cells denoted -RFP) (lentiviral vector pDEST/pHIV-H2BmRFP-rfa_verB, a gift from Matthew Smalley, University of Cardiff). mCherry^+^/RFP^+^ cells were selected by fluorescence-activated cell sorting (FACS). Luciferase only transduced cells were enriched by culturing the cells in DMEM with 10% FBS containing blasticidin for 2-3 passages.

#### Lentiviral transfection of fibroblasts with PmiR146a-GFP plasmid

The miR146a-GFP plasmid (35 μg) (Kang et al., 2015a) was combined with packaging vectors pMD2.G (11 μg) and psPAX2 (25.6 μg) in 18 mL Optimem medium containing 216 μL Lipofectamine-2000, and incubated at room temperature for 20 minutes. Lentivirus containing medium from HEK293T cells was harvested at 24 and 48 hours, centrifuged at 300*g* and filtered through a 0.2 μm syringe filter before use for cell transfections. Fibroblasts were seeded in 25 cm^2^ flasks in medium containing miR146a-GFP lentiviral particles and 4 μg mL^-1^ polybrene and for incubated 24 hours. Virus containing medium was replenished after 24 hours. Transfected fibroblasts were then maintained in DMEM with 10% FBS containing puromycin for 2-3 passages.

### *In vivo* procedures

All animal work was carried out under UK Home Office Project Licenses 70/7413 and P6AB1448A granted under the Animals (Scientific Procedures) Act 1986 (Establishment Licence, X702B0E74 70/2902) and was approved by the ’Animal Welfare and Ethical Review Body’ at The Institute of Cancer Research (ICR). All mice used were female and aged between 6-9 weeks and 18-25 g in weight at the beginning of an experiment. Syngeneic studies were carried out in BALB/c mice. Human cell line studies were carried out in NOD RAG gamma (NRG) mice (NOD-Rag1^-/-^ IL2rg^-/-^), implanted with 90 day, sustained release 0.36 mg 17β-oestradiol pellets (Innovative Research of America, NE-121) 4-5 days prior to tumour cell implantation. All mice were housed in individually ventilated cages, monitored daily by ICR Biological Services Unit staff and had food and water *ad libitum*. In all cases, experiments were terminated if the primary tumour reached a maximum allowable diameter of 17 mm, if thoracic IVIS signal exceeded 1 x 10^9^ photons per second or if a mouse showed signs of ill health. For non-tumour bearing mice, mice were randomised into groups based on body weight. For experimental and spontaneous metastasis experiments, mice were randomised prior to drug administration based on IVIS signal or tumour volume, respectively. Dosing schedules for individual experiments are presented in the figures.

#### Chemotherapy treatment of mice

Mice were dosed via 100 μL intraperitoneal injection with a combination of doxorubicin (2.3-2.7 mg kg^-1^, Selleckchem S1208) and cyclophosphamide (40-44 mg kg^-1^, Sigma 50850) in 0.9% NaCl. Doxorubicin and cyclophosphamide were stored separately at a 2X concentration (1.7 mM and 57 mM, respectively) and were mixed at a 1:1 ratio immediately prior to injection. Vehicle-treated mice were injected intraperitoneally with 100 μL 0.9% NaCl. *Navitoclax treatment of mice*: Navitoclax was stored as aliquots of 100 mM solution in DMSO at −80°C and was administered as a dose of 50 mg/kg daily dose. For each dose one aliquot was thawed and diluted into either corn oil to a final concentration of 5 mM (5% DMSO final concentration) or, where indicated, in 10% ethanol:30% PEG 400 (Sigma 202398):60% Phosal 50 PG (LIPOID, LLC 368315). Vehicle-treated mice were treated with a 5% solution of DMSO with corn oil or the Phosal mixture. Mice were dosed via oral gavage daily for 5 days using a 200 µL volume.

#### Spontaneous metastasis assays

2 x 10^5^ D2A1-m2 cells were injected orthotopically into the 4^th^ mammary fat pad under general anaesthesia. Tumour growth was measured twice a week using callipers up to a maximum diameter of 17 mm. Tumour volume was calculated as 0.5236 x [(width + length)/2]^3. At endpoint tumours were excised and weighed. *Intravenous inoculation*: 5 x 10^5^ D2A1, 1 x 10^6^ D2.OR or 2 x 10^6^ ZR-75-1 cells (transfected with either -Luc or -mChLuc2 vectors) were injected into the lateral tail vein of mice. Metastatic growth was monitored by repeated IVIS imaging starting at ∼90 minutes following inoculation. At endpoint lungs were excised and, where indicated, IVIS imaged *ex vivo*.

#### IVIS imaging

Mice were injected intraperitoneally with 150 mg kg^-1^ D-luciferin (Caliper Life Sciences) in 100 μL and mice imaged *in vivo* using an IVIS imaging chamber (IVIS Illumina II). Luminescence measurements (photons per second per cm^2^) were acquired over 1 minute and analysed using the Living Image software (PerkinElmer) using a constant size region of interest over the tissues. Alternatively, >15 minutes after D-luciferin injection, dissected lungs and/or livers were imaged *ex vivo*.

#### Quantification of metastatic burden and immunostaining

Tissues were formalin-fixed and paraffin-embedded (FFPE). 3-4 µm lung, liver or bone FFPE sections were cut, dewaxed in xylene, re-hydrated through ethanol washes and stained with haematoxylin and eosin (H&E). Slides were scanned using Hamamatsu microscope with a NanoZoomer XR camera using the 20x objective and file names blinded. For quantifying tumour burden, total number of individual nodules was counted manually in 1-3 sections approximately 150 μm apart, per tissue. Lung metastatic area was quantified as the mean size of the nodules or percentage of the area of the metastatic nodules normalised to the total lung area. Alternatively, sections were subjected to high-temperature antigen retrieval, blocked using avidin/biotin, before incubation with primary antibodies. Immunohistochemical detection was achieved with the VectaStain ABC system. Stained sections were scanned as described above and file names blinded. p21, γH2Ax or MCL-1 positive cells were quantified either manually from ≥6 randomly selected 0.1 mm^2^ fields of view per lung section, or by automated cell detection of whole tissue sections using QuPath <u>(Bankhead et al., 2017)</u>, avoiding regions containing bronchi and bronchioles. Alternatively, FFPE sections of mouse lungs were stained with p21 alongside αSMA, endomucin or EpCAM and DAPI, then incubated with the appropriate secondary antibody conjugated to fluorophores. Fluorescent stained sections were scanned as described above. p21 positive cells were counted across the whole section. p21 positive cells were then scored for positivity of αSMA, endomucin or EpCAM. Representative higher power images were collected on a Leica SP8 microscope using a 40X oil immersion objective.

#### Senescence-associated β-galactosidase staining of lung tissue sections (SA-β-Gal)

Lungs were excised, lobes dissected, and left lobe was frozen in OCT embedding matrix (CellPath KMA-0100-00A) on dry ice. Frozen blocks were sectioned in cryostat to 10 µm thick sections. Sections were air-dried before fixing for 10 minutes in 2% formaldehyde (Sigma F8775) : 0.2% glutaraldehyde (Sigma G6257) solution in PBS. Slides were washed three times with PBS and incubated with X-gal staining mixture (Sigma CS0030) at 37°C without CO_2_ overnight. Next day, the slides were washed in PBS and counterstained with haematoxylin (Sigma MHS1) before mounting.

#### Ex vivo lung culture

Following sacrifice, mouse lungs were perfused with agarose, removed and allowed to solidify in cold PBS for ∼1 hour. Lungs were cut into 150 µm slices and cultured on an inert matrix in M199 medium for 24 hours before seeding of tumour cells. *RNA extraction from tissue*: Mice were sacrificed 7 or 10 days after the final chemotherapy or vehicle treatment. The lungs were excised and the left lobe was fixed and processed for immunohistochemical staining as described above. The right lobes were immediately snap-frozen in liquid nitrogen in a cryovial and then stored at −80°C. To extract RNA, a ∼20 mg chunk of frozen lung tissue was collected using a scalpel and placed in 1 mL RLT buffer (Qiagen) with 1:100 β-mercaptoethanol in a Precellys silicone bead dissociation tube (Bertin-Corp). Tissue were dissociated by rigorous vortexing for 1 minute using the Precellys 24 Tissue Homogenizer and lysates immediately frozen at −80°C. RNA extraction was carried out using the Qiagen RNeasy kit according to the manufacturer’s protocol.

### In vitro studies

#### RTqPCR

RNA was isolated using the Qiagen RNeasy kit and cDNA was generated by using the QuantiTect reverse transcription kit (Qiagen) according to the manufacturer’s instructions. RTqPCR was performed with human or mouse Taqman Gene Expression Assay probes on an Applied Biosciences QuantStudio6 Flex Real-time PCR machine and relative quantification was performed using QuantStudio Real-time PCR software. Each reaction was performed in triplicate. Relative expression levels were normalised to *B2m/B2M*, *Gapdh/GAPDH,* or *Tmem199/TMEM199* endogenous controls, and ≥2 controls were used in each experiment. In control 3T3 fibroblast RNA samples, the *Mmp3* and *Il1a* probes did not amplify during the run. Consequently, RQ values were calculated using an assumed CT value of 40 for the control samples, based on the maximum number of cycles used. Confidence intervals were set at 95% for all assays.

#### NanoString gene expression analysis

RNA was extracted from frozen lung tissue as described above. 80 ng RNA was hybridised with the Mouse PanCancer Immune panel or Mouse PanCancer Pathways panel and processed using the nCounter SPRINT Profiler (NanoString) following the manufacturer’s instructions. Raw NanoString data was pre-processed using R package NanoStringNorm (v1.2.1) (Waggott et al., 2012) and further normalised using voom (TMM normalisation) followed by differential gene expression analysis with R package limma (v3.40.6) (Ritchie et al., 2015) Genes with absolute log_2_ fold change ≥ 0.585 and FDR adjusted *p* value < 0.05 were considered significantly different. Principle component analysis (PCA) was performed using R package FactoMineR(v2.3). All analyses were performed in R statistical programming language (v3.6.14). Visualisations were generated using in-house plotting libraries. Immune cell population abundance was performed as described previously (Jenkins et al., 2022).

#### Chemotherapy induction of senescence

Fibroblasts were cultured in DMEM with 10% FBS in 75 cm^2^ flasks to ∼75% confluence, then medium was replaced with DMEM with 10% FBS containing chemotherapeutic agents. Fibroblasts were incubated in medium containing the drug for 24 hours, washed twice with PBS and then cultured in fresh DMEM with 10% FBS. Senescent fibroblasts were maintained in the same flasks for up to 6 months without further passage, growth medium was replaced weekly with fresh DMEM plus 10% FBS.

#### Senescent cell imaging

Fibroblasts induced into senescence as described above were seeded in 6-well or 96-well plates at 80-90% confluency 7-21 days following drug withdrawal and incubated for 24 hours before staining. Control fibroblasts were seeded into a 6-well plate and cultured for 2-3 days until they matched the confluency of the senescent fibroblasts. Fibroblasts were washed with PBS, fixed for 10 minutes with 2% paraformaldehyde and 0.2% glutaraldehyde, washed twice with PBS and 1.5 mL staining solution (SA-β-Gal histochemical staining kit, Sigma CS0030) containing X-Gal was added. Plates were sealed with Parafilm and incubated at 37°C in a non-humidified incubator at atmospheric CO_2_ concentration. After 24 hours the cells were washed with PBS and imaged at 100x magnification. For live cell imaging of GFP signal, cells were imaged using either the EVOS fluorescence microscope or the Incucyte live cell imaging fluorescence microscope using Excitation 470/Emission 525 wavelengths.

#### Conditioned media (CM)

CM was collected from chemotherapy-treated fibroblasts 7-30 days following chemotherapy withdrawal. Chemotherapy-treated or control fibroblasts were washed twice and then cultured in serum-free DMEM. After 24 hours, CM was collected, centrifuged at 300*g,* filtered through a 0.2 μm pore filter and used fresh or stored at −80°C for future use. CM was used within 6 months of freezing and was filtered post-thawing.

#### Cytokine arrays

CM was freshly collected as described above from chemotherapy-treated human and mouse fibroblasts or from control fibroblasts. CM was collected independently from two flasks per condition, and 100 μL added to duplicate RayBiotech G2000 array slides. Wash and incubation steps were performed according to the manufacturer’s protocol. Fluorescence signal detection at 532 nm was performed on a GenePix 4000B array scanner. Median fluorescence intensity (MFI) was quantified using ImageJ software. Fold changes were calculated by dividing the mean MFI for the two chemotherapy-treated CM replicates by the mean MFI for the two control replicates for each fibroblast type.

#### Phosphoprotein arrays

Tumour cells were seeded into ultra-low attachment U-bottom 96-well plates in DMEM with 2% FBS at density of 12,500 cells per well and incubated for 48 hours. 192 spheroids were divided into 2 groups of 96 and placed into 15 mL Falcon tubes, washed once with PBS and then incubated in 12 mL serum-free DMEM for 2 hours at 37°C on a roller. Spheroids were centrifuged at 260*g*, medium removed and replaced with 10 mL of control or chemotherapy-treated fibroblast CM and incubated for 30 minutes at 37°C on a roller. Spheroids were then washed 2X in 10 mL ice-cold PBS then lysed in 100 μL Cell Signalling Technologies lysis buffer containing 1 mM PMSF. Cell lysates were sonicated in an ice-cold water bath for 10 minutes, frozen to −80°C for 30 minutes, thawed, centrifuged at 16,000*g* for 10 minutes at 4°C, the supernatant was transferred to a fresh tube on ice and the pellet discarded. Protein concentration was measured using the Millipore Direct Detect Spectrometer according to the manufacturer’s protocol. Protein samples were diluted in array blocking buffer and 150 μL was added to the Pathscan antibody-array slides (Cell Signalling Technologies #7982). Wash and incubation steps, and sample detection using a chemiluminescent HRP substrate were performed according to the manufacturer’s protocol. Luminescence was imaged using the Odyssey Fc Imaging System. Median fluorescence intensity (MFI) was quantified using ImageJ software. Fold changes were calculated by dividing the mean MFI for the two technical replicate spots for each target in the senescent CM treated spheroids by the mean MFI for the two technical replicate spots for each target in the control CM treated spheroids.

#### Co-culture spheroid assays

1,000 tumour cells and 1,000 fibroblasts were co-seeded into ultra-low attachment U-bottom 96-well plates and incubated for 24 hours. Spheroids were treated with doxorubicin (0 – 0.85 µM) and incubated for a further 72 hours before imaging on an EVOS microscope. Alternatively, 5,000 tumour cells and 5,000 fibroblasts were co-seeded, incubated for 24 hours before addition of 0.17 µM doxorubicin or vehicle. 72 hours later, spheroids were treated with 4.5 µM navitoclax or vehicle and incubated for a further 72 hours before being imaged and assayed by CellTiter-Glo.

#### BCL-2 family inhibitors dose response assays

Chemotherapy-treated fibroblasts were seeded into 96-well or 24-well plates for CellTiter-Glo or crystal violet analysis, respectively. Chemotherapy-treated fibroblasts were seeded at ∼95% confluence in parallel with control fibroblasts at ∼70% confluence and incubated for 24 hours prior to treatment with BCL-2-family inhibitors, alone or in combination, as indicated for 72 hours before assaying cell viability by CellTiter-Glo or staining with 0.5% crystal violet in methanol.

### Statistics

Statistics were performed using GraphPad Prism 8 or 9. All data is represented as the mean value ± standard error of the mean. Comparisons between two groups were made using two-tailed, unpaired Student’s *t*-tests. If there was a significant difference between the standard deviations of the two groups, then Welch’s correction was applied. If the analysis did not pass the Shapiro-Wilk normality test, then groups were analysed by Mann-Whitney *U*-test. If more than 2 groups were compared One-way ANOVA analysis was performed with Dunnett’s correction for multiple comparisons. If a significant difference between the standard deviations of the means was measured then the Brown-Forsythe and Welch correction was applied. If data did not pass the Shapiro-Wilk normality test then a Kruskal-Wallis ANOVA was performed with Dunn’s correction for multiple comparisons.

## Supplemental Information

Supplemental Information includes 5 Supplementary Figures and 4 Supplementary Tables

## Acknowledgements

This work was funded as part of Programme Funding to the Breast Cancer Now Toby Robins Research Centre (CMI) and a Chairman’s ICR PhD studentship (DP). We acknowledge NHS funding to the NIHR Biomedical Research Centre at the Royal Marsden and the ICR. We would like to thank Steve Elledge for the PmiR146a-GFP plasmid, Marjan Iravani, David Vicente and Sarah Ash for experimental contribution, the ICR Biological Services Unit and Dr Naomi Guppy and her team in the Breast Cancer Now Toby Robins Research Centre Nina Barough Pathology Core Faculty for histopathology support.

## Author Contributions

Conceptualization (DWP, IS, LO’L, CMI)

Investigation (DWP, IS, SH, DR, RB, LO’L, CMI)

Writing - Review & Editing (DWP, IS, CMI with input from all other authors)

Funding Acquisition (SH, CMI)

## Declaration of interests

The authors have no financial or non-financial competing interests to declare

## Notes

### Competing Interest Statement

The authors have declared no competing interest.

### Summary of Updates

New data provided

