## Supplementary Figs 1-5, Supplementary Tables 1-4 for "Therapy-induced normal tissue damage promotes breast cancer metastasis"

**SUPPLEMENTARY MATERIAL**

**Supplementary Figures 1 - 4**

Supplementary Figure 1 | Chemotherapy limits tumour growth *in vivo*.

Supplementary Figure 2 | Effects of chemotherapy treatment on immune cell populations in the lung.

Supplementary Figure 3 | Response of chemotherapy-treated fibroblasts to BCL-2 family inhibitors *in vitro*.

Supplementary Figure 4 | Principal component analysis (PCA) of NanoString PanCancer Immune and PanCancer Pathways panel data.

Supplementary Figure 5. | Effects of chemotherapy and navitoclax treatment on immune cell populations in the lung and higher power images from Figure 7.

**Supplementary Tables 1 - 4**

Supplementary Table 1 | NanoString (PanCancer Immune panel) differentially expressed genes

Supplementary Table 2 | NanoString (PanCancer Pathways panel) differentially expressed genes

Supplementary Table 3 | RTqPCR probes

Supplementary Table 4 | Antibodies

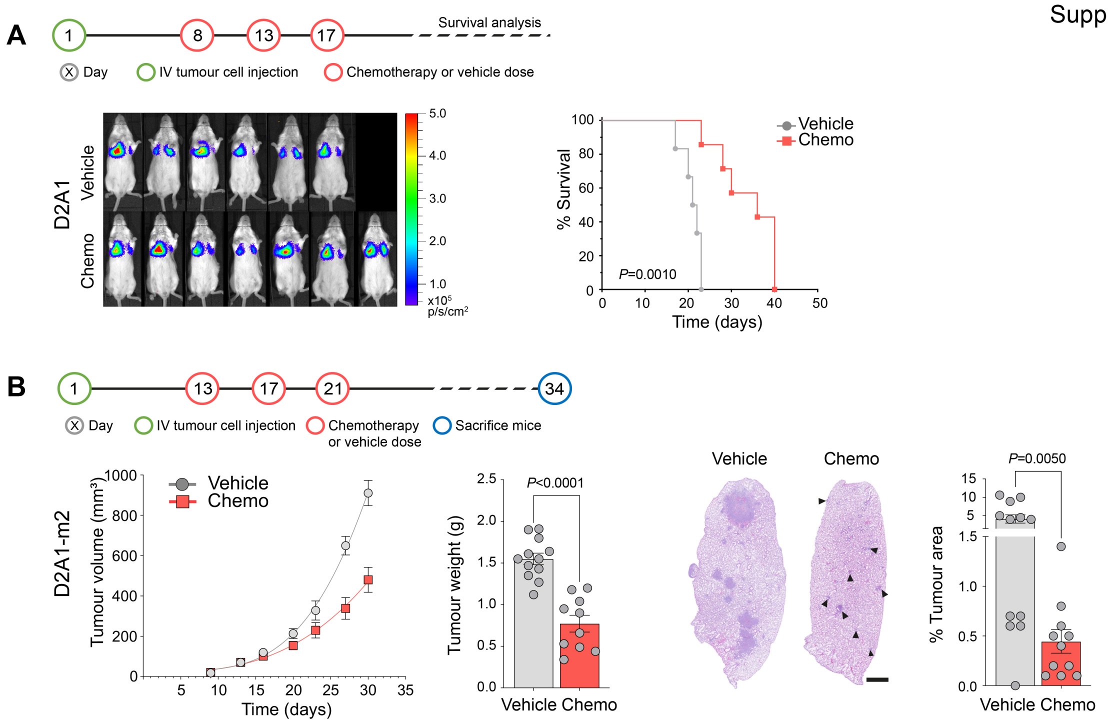


**Supplementary Figure 1.** Chemotherapy limits tumour growth *in vivo.* **A** Experimental timeline of BALB/c mice inoculated intravenously on Day 1 with 5 x $10^5$ D2A1-mChLuc2 tumour cells (n=6-7 mice per group) followed by combination chemotherapy or vehicle treatments starting on Day 8. Shown are IVIS images taken ~90 minutes after tumour cell injection and Kaplan-Meier survival analysis (Log-rank (Mantel-Cox) test). Mice were culled individually when thoracic IVIS signal exceeded 1 x $10^9$ photons per second or if a mouse showed signs of ill health. **B** Experimental timeline of BALB/c mice inoculated orthotopically (4th mammary fat pad) on Day 1 with 2 x $10^5$ D2A1-m2 tumour cells (n=10-12 mice per group). Shown are: primary tumour growth measured twice weekly; tumour weight at necropsy on Day 34 (±SEM, unpaired t-test); representative images of lung H&E stained sections, arrowheads indicate micrometastatic deposits in chemotherapy-treated lungs (scale bar, 1 mm); quantification of spontaneous metastasis to the lungs in 3 lung sections per mouse (mean % tumour burden per lung section ±SEM, unpaired t-test).

Supp. Figure 2

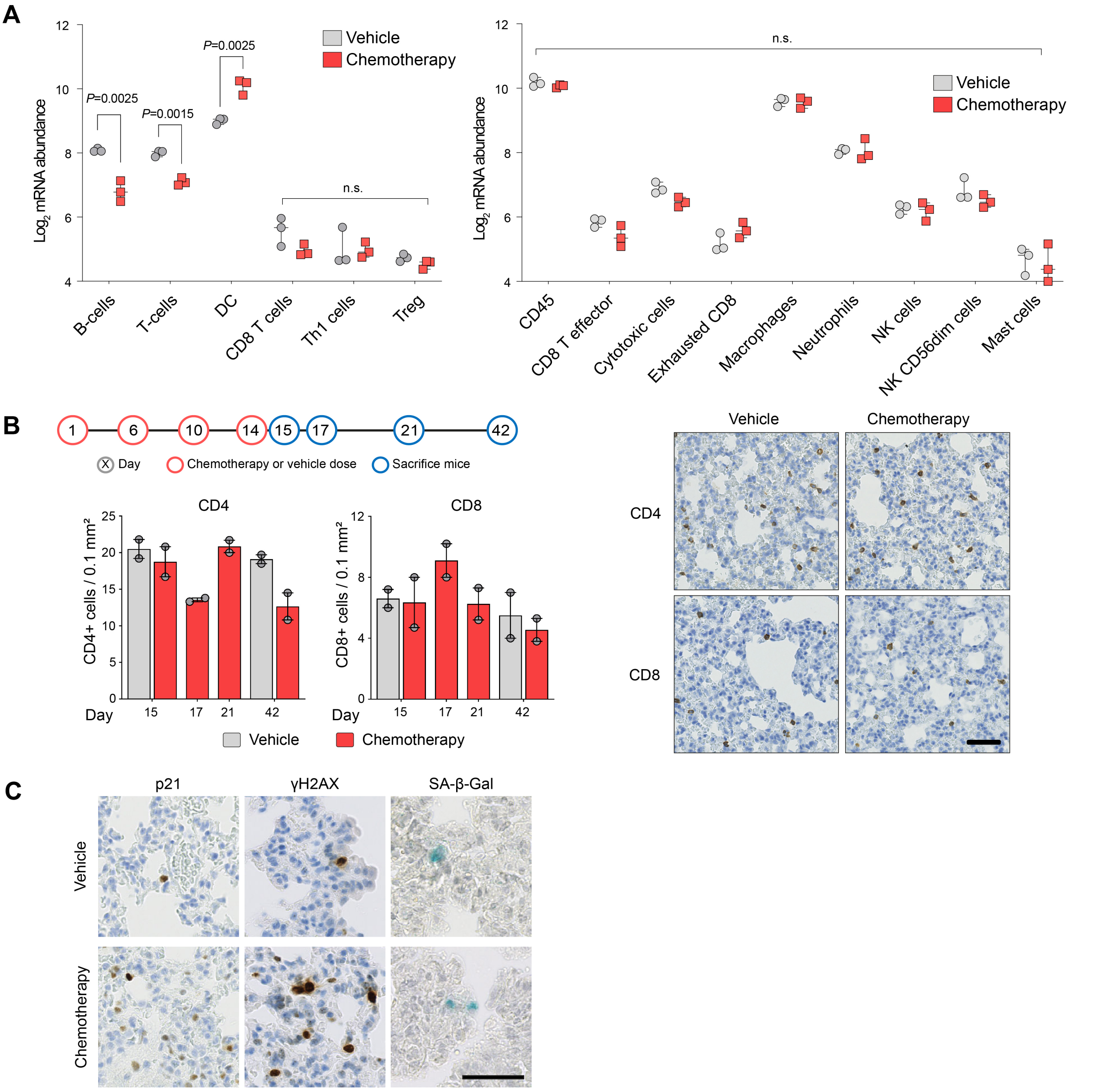


**Supplementary Figure 2.** Effects of chemotherapy treatment on immune cell populations in the lung. Related to Figure 2. **A** NanoString immune cell population abundance signatures for data shown in Figure 2A,B (median values, ±min to max; multiple unpaired *t*-tests, n.s. = non-significant). **B** BALB/c mice treated with a course of chemotherapy or vehicle as indicated and sacrificed on Day 15, 17, 21 and 42 (1, 3, 7 and 28 days after the last treatment dose). Lung sections were stained for CD4 or CD8, and the number of positively stained cells was quantified in a blinded fashion, using ImageJ software. Positive cells were counted in 6, randomly selected, 0.1 mm$^2$ fields of view per lung section. Shown are the mean number of CD4$^+$ and CD8$^+$ cells per 0.1 mm$^2$ ±SEM. Representative images are shown from vehicle-treated mice sacrificed on Day 15 and from chemotherapy-treated mice sacrificed on Day 21. Scale bar, 50 μm. **C** Higher power images from Figure 2D (scale bar, 50 μm).

Supp. Figure 3

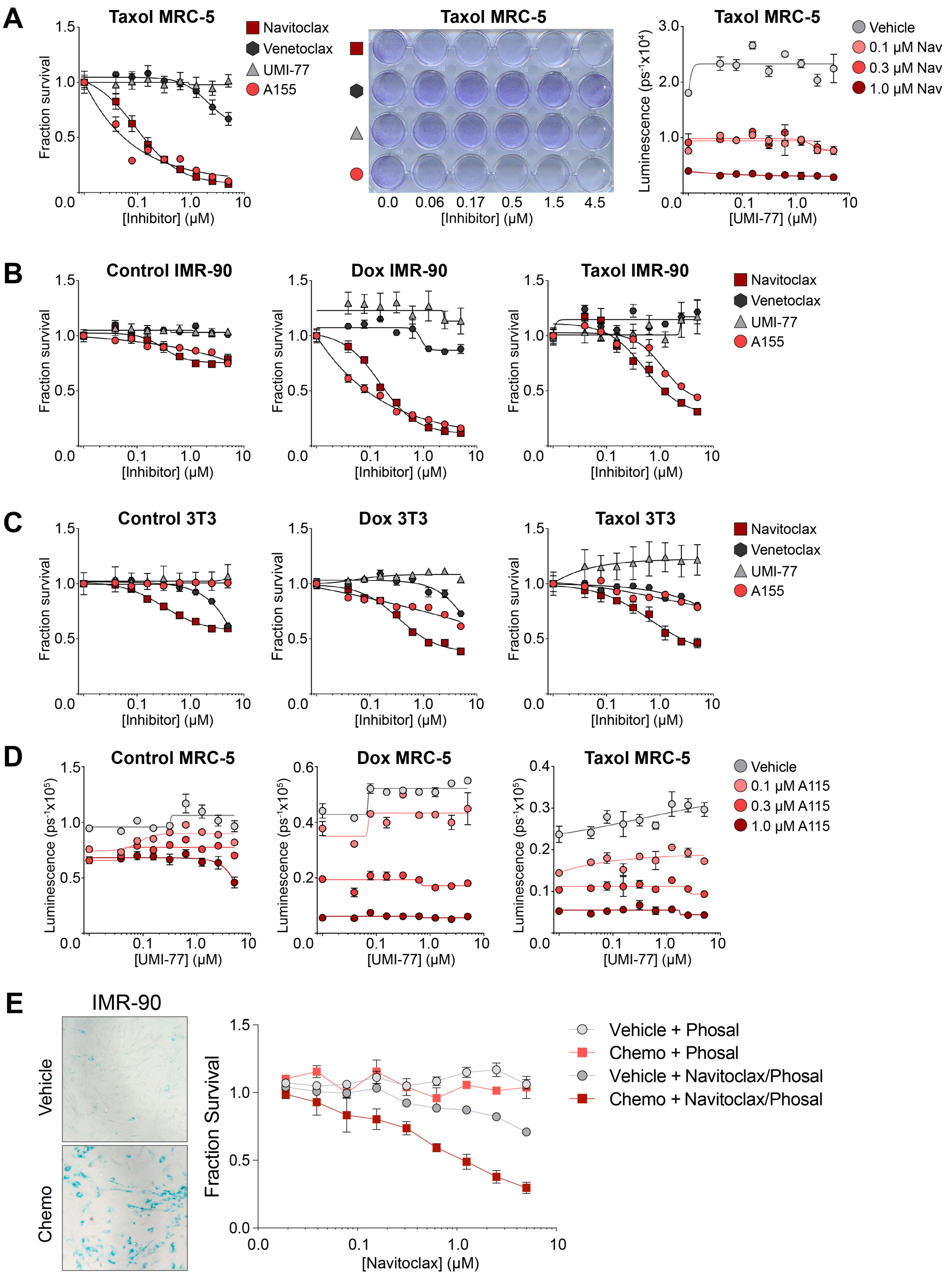


**Supplementary Figure 3.** Response of chemotherapy-treated fibroblasts to BCL-2 family inhibitors *in vitro*. Related to Figure 5. **A** MRC-5-146a fibroblasts were induced to senescence by 24 hour treatment with 1.2 μm docetaxel. 9 days after treatment withdrawal, chemotherapy-treated or control fibroblasts were plated into 96-well plates (n=3 wells per condition) or 24 well plates (n=1 well per condition). After 24 hours fibroblasts were treated with individual BCL-2 family inhibitors, or with a range of concentrations of UMI-77 in combination with 3 concentrations of navitoclax or DMSO control. 72 hours later, cell viability was measured by CellTiter-Glo (left and right panels, mean values ±SEM) or plates were stained with crystal violet (middle panel). **B** IMR-90-146a fibroblasts were treated for 24 hours with 1.7 μm doxorubicin or 1.2 μm docetaxel. 20 days after treatment withdrawal, chemotherapy-treated or control fibroblasts were plated into 96-well plates (n=3 wells per condition). After 24 hours fibroblasts were treated with BCL-2 family inhibitors as indicated and incubated for 72 hours. Cell viability was measured by CellTiter-Glo (mean values ±SEM). **C** 3T3-146a fibroblasts were treated for 24 hours with 0.17 μm doxorubicin or 0.12 μm docetaxel. 7 days after treatment withdrawal, chemotherapy-treated or control fibroblasts were plated into 96-well plates (n=3 wells per condition). After 24 hours fibroblasts were treated with BCL-2 family inhibitors as indicated and incubated for 72 hours. Cell viability was measured by CellTiter-Glo (mean values ±SEM). **D** MRC-5-146a fibroblasts were treated for 24 hours treatment with 1.7 μm doxorubicin or 1.2 μm docetaxel. 10 days after treatment withdrawal, chemotherapy-treated or control fibroblasts were plated in 96-well plates (n=3 wells per condition). After 24 hours fibroblasts were treated with range of concentrations of UMI-77 alone or in combination with 3 concentrations of A115 and incubated for 72 hours. Cell viability was measured by CellTiter-Glo (mean values ±SEM). **A-D** Equivalent results were obtained in 2 or 3 independent experiments. **E** IMR-90 fibroblasts were treated for 24 hours with 0.17 μm doxorubicin. 17 days after treatment withdrawal senescent cells were stained and visualised using SA-β-Gal staining kit. 7 days after treatment withdrawal, chemotherapy-treated or control fibroblasts were plated into 96-well plates (n=2 wells per condition). After 24 hours fibroblasts were treated with Navitoclax reconstituted in Phosal and incubated for 72 hours. Cell viability was measured by CellTiter-Glo (mean values ±SEM).

Supp. Figure 4

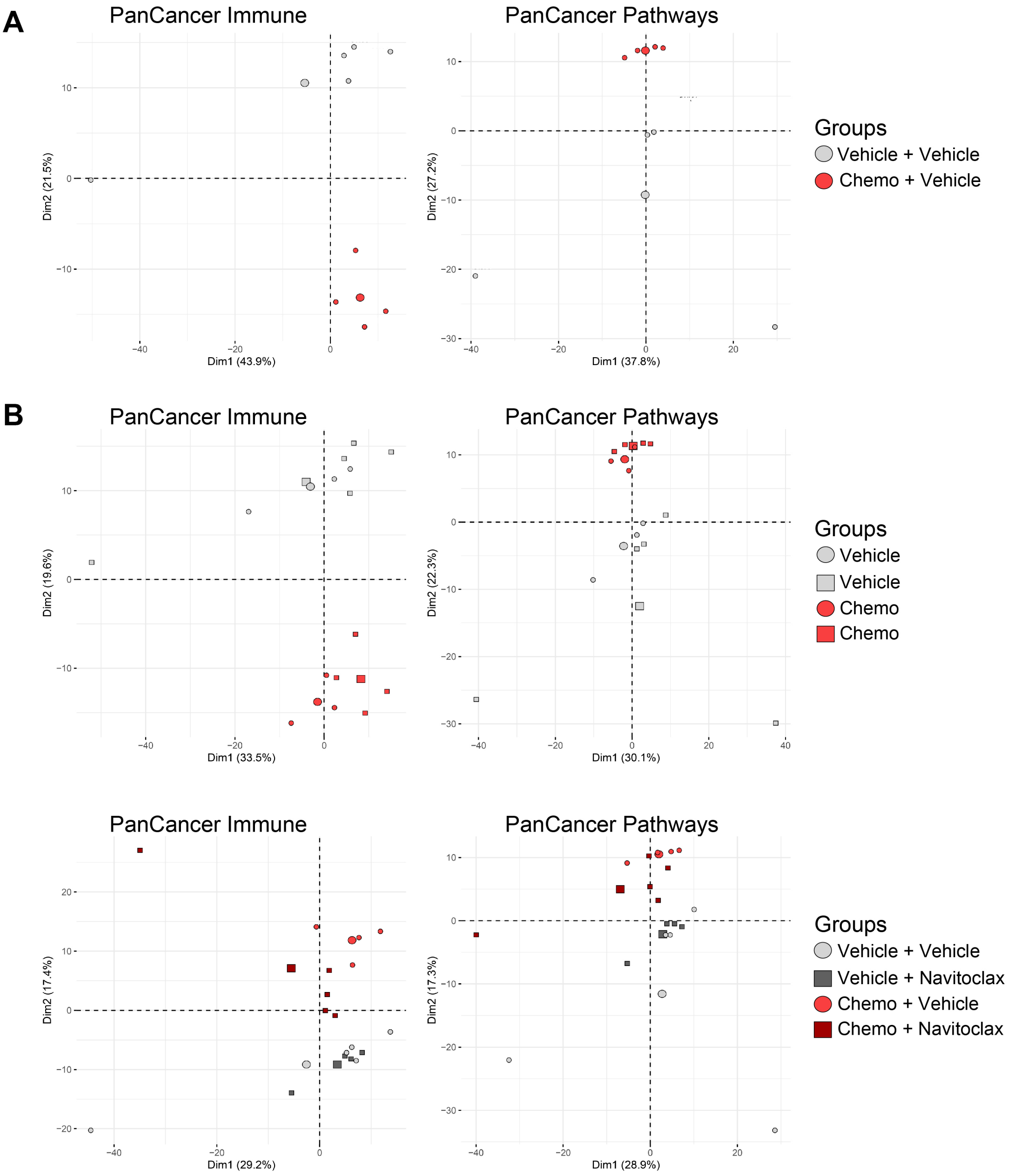


**Supplementary Figure 4.** Principal component analysis (PCA) of NanoString PanCancer Immune and PanCancer Pathways panel data. Related to Figures 2A,B and 7A-C. Shown are the PCA plots based on expression of the 750 target genes in each panel from: **A** Vehicle + Vehicle and Chemo + Vehicle mice from Figure 7B and C; **B** Panel A samples combined with Vehicle and Chemo samples from Figure 2B; **C** All samples from Figure 7B-C.

Supp. Figure 5

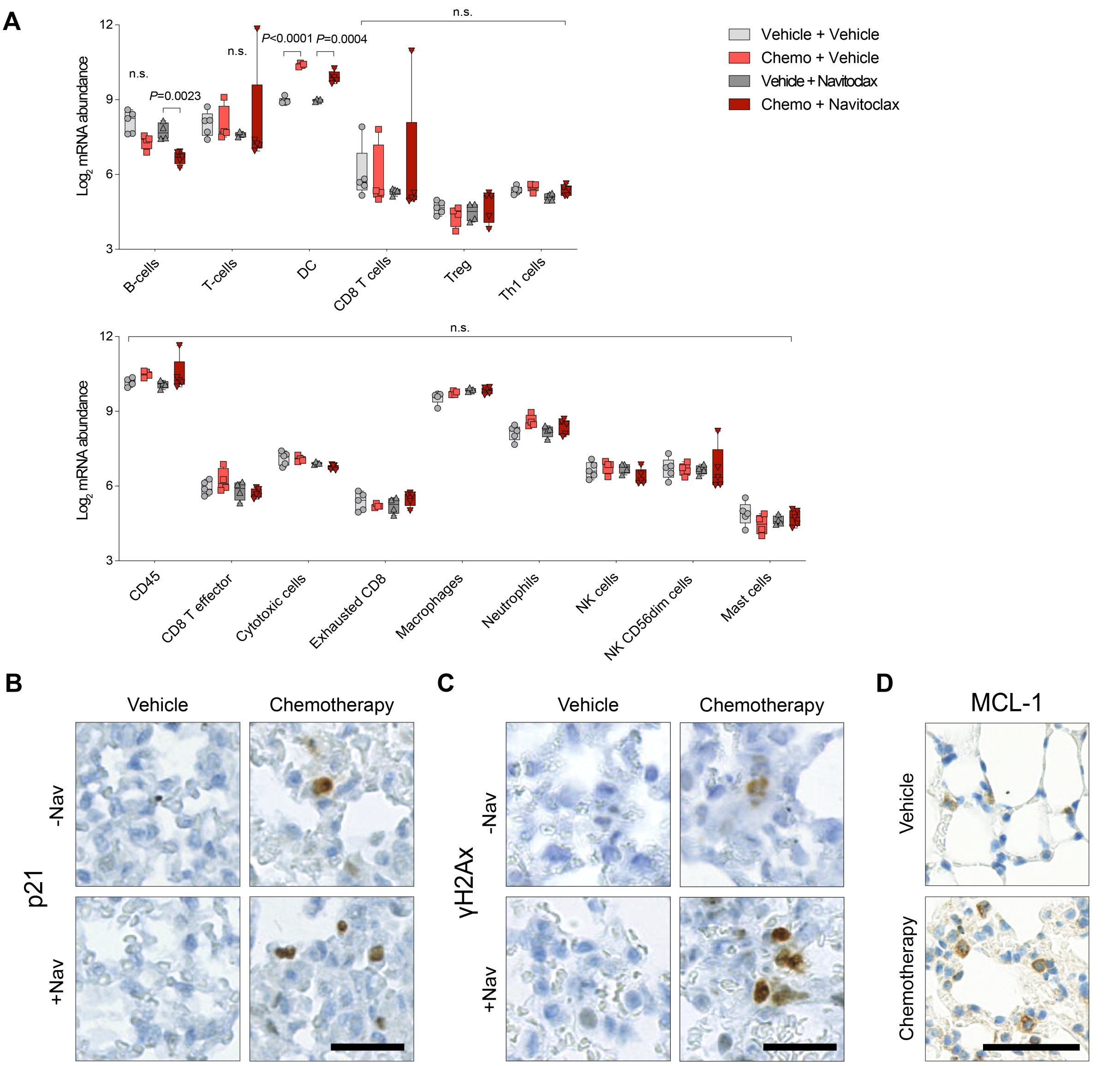


**Supplementary Figure 5.** Effects of chemotherapy and navitoclax treatment on immune cell populations in the lung and higher power images from Figure 7. **A** NanoString immune cell population abundance signatures related to Figure 7B-C. (median values, ±min to max; multiple unpaired *t*-tests, n.s. = non-significant). **B - D** Higher power images from Figure 7D, Figure 7E and Figure 7G, respectively (scale bar, 50 μm).

Supplementary Table 1: Top 150 & bottom 50 NanoString PanCancer Immune panel hits ranked by fold change (FC)

| Fig. 2e Vehicle vs Chemo | | | Fig. 8b Vehicle vs Chemo | | | Fig. 2e+8b Veh. vs Chemo | | | Fig. 8c Vehicle ± Nav | | | Fig. 8c Chemo ± Nav | | |
|---|---|---|---|---|---|---|---|---|---|---|---|---|---|---|
| Factor | FC | adj.P.Val | Factor | FC | adj.P.Val | Factor | FC | adj.P.Val | Factor | FC | adj.P.Val | Factor | FC | adj.P.Val |
| Cdkn1a | 4.87 | 1.38E-04 | Ccl2 | 6.09 | 3.62E-07 | Ccl2 | 5.31 | 3.27E-08 | Mx1 | 3.48 | 6.28E-01 | Mpped1 | 6.63 | 6.55E-02 |
| Ccl8 | 4.70 | 6.72E-03 | Cxcl10 | 4.79 | 5.02E-05 | Cxcl10 | 4.46 | 2.69E-06 | Itga2b | 2.75 | 1.12E-01 | Cfd | 3.92 | 7.82E-01 |
| Ccl2 | 4.63 | 1.38E-04 | Cdkn1a | 3.59 | 6.69E-05 | Cdkn1a | 4.18 | 4.14E-07 | Mpped1 | 1.88 | 9.10E-01 | Itga2b | 2.52 | 8.69E-02 |
| Cxcl10 | 4.16 | 3.06E-03 | C3ar1 | 3.52 | 5.21E-07 | Ccl8 | 3.66 | 3.15E-04 | Ppbp | 1.84 | 8.78E-01 | Rag1 | 2.27 | 8.52E-01 |
| Ccl7 | 3.13 | 1.17E-03 | Ccr3 | 3.36 | 4.35E-01 | Ccl7 | 3.24 | 1.25E-06 | Defb1 | 1.77 | 9.90E-01 | Slamf1 | 2.17 | 1.12E-01 |
| C8a | 3.02 | 3.99E-01 | Ccl7 | 3.36 | 4.88E-05 | Ccr3 | 2.77 | 4.21E-01 | Crp | 1.68 | 8.78E-01 | A2m | 2.03 | 3.26E-01 |
| Ccl12 | 3.00 | 3.12E-03 | Lif | 2.97 | 1.08E-08 | Ccl12 | 2.75 | 1.62E-05 | Il13ra2 | 1.66 | 8.78E-01 | Bcl2l1 | 1.81 | 6.94E-04 |
| Il19 | 2.69 | 3.35E-02 | Ccl8 | 2.85 | 2.52E-02 | C3ar1 | 2.74 | 2.64E-07 | Ctsg | 1.63 | 8.78E-01 | Marco | 1.78 | 2.04E-01 |
| Birc5 | 2.69 | 2.89E-01 | Usp9y | 2.75 | 4.35E-01 | Lif | 2.52 | 3.36E-09 | Tnfsf4 | 1.62 | 8.78E-01 | Dmbt1 | 1.77 | 7.25E-01 |
| Mbl2 | 2.59 | 3.90E-01 | Cx3cr1 | 2.74 | 5.21E-07 | Trem2 | 2.43 | 9.29E-08 | Serpinb2 | 1.60 | 9.73E-01 | Il2ra | 1.75 | 6.03E-01 |
| S100b | 2.54 | 3.33E-01 | Il6 | 2.65 | 1.36E-02 | Usp9y | 2.43 | 3.03E-01 | Chit1 | 1.55 | 9.90E-01 | Ctsg | 1.72 | 3.42E-01 |
| Trem2 | 2.36 | 1.38E-04 | Ccl17 | 2.59 | 5.10E-04 | Birc5 | 2.31 | 7.74E-02 | Slamf1 | 1.55 | 8.78E-01 | H2-DMb1 | 1.65 | 4.94E-01 |
| Ccr3 | 2.29 | 8.25E-01 | Ccl12 | 2.53 | 6.23E-04 | Ccl17 | 2.30 | 1.02E-04 | Il1rapl2 | 1.54 | 8.96E-01 | Rorc | 1.60 | 7.70E-01 |
| Prg2 | 2.21 | 1.44E-01 | Trem2 | 2.49 | 2.15E-06 | Cx3cr1 | 2.25 | 3.39E-07 | Il4 | 1.52 | 8.78E-01 | Tnfrsf8 | 1.58 | 6.30E-01 |
| Egr1 | 2.17 | 4.31E-03 | C1qa | 2.22 | 1.91E-04 | Il6 | 2.10 | 1.25E-02 | Il11 | 1.48 | 8.78E-01 | Masp2 | 1.57 | 8.05E-02 |
| Rag1 | 2.15 | 8.92E-01 | S100a8 | 2.21 | 4.50E-03 | C1qb | 2.09 | 2.52E-05 | Lyve1 | 1.47 | 8.78E-01 | Defb1 | 1.55 | 9.07E-01 |
| Thbs1 | 2.15 | 1.67E-02 | Ccl9 | 2.19 | 4.88E-05 | C1qa | 2.09 | 1.54E-05 | Klrb1c | 1.45 | 8.78E-01 | Cd8a | 1.54 | 8.87E-01 |
| Lif | 2.14 | 1.28E-04 | C1qb | 2.17 | 4.03E-04 | C4b | 1.96 | 1.10E-06 | Cd70 | 1.44 | 9.10E-01 | Klra3 | 1.54 | 4.26E-01 |
| Usp9y | 2.14 | 7.11E-01 | Ccr5 | 2.11 | 4.73E-04 | Cd80 | 1.86 | 1.28E-03 | Raet1 | 1.42 | 8.78E-01 | Ppbp | 1.51 | 7.06E-01 |
| C3ar1 | 2.13 | 3.69E-03 | Clec5a | 2.07 | 5.35E-04 | Ccl9 | 1.85 | 2.29E-05 | Trem2 | 1.41 | 5.74E-01 | Crp | 1.51 | 5.17E-01 |
| Cd80 | 2.10 | 1.95E-02 | Ccr2 | 2.04 | 6.99E-07 | Cfb | 1.83 | 3.53E-03 | C3ar1 | 1.40 | 8.78E-01 | Lyve1 | 1.51 | 3.35E-01 |
| Ccl17 | 2.05 | 3.51E-02 | Birc5 | 1.98 | 3.25E-01 | Clec5a | 1.83 | 1.89E-04 | C8a | 1.39 | 9.90E-01 | H60a | 1.50 | 6.03E-01 |
| Cfb | 2.03 | 5.18E-02 | Cxcr1 | 1.92 | 8.79E-02 | Col4a1 | 1.79 | 2.69E-06 | Klra21 | 1.39 | 8.96E-01 | Mertk | 1.50 | 6.96E-02 |
| Cxcl9 | 2.02 | 1.00E-01 | C4b | 1.91 | 6.90E-05 | C8a | 1.79 | 4.02E-01 | Il22ra2 | 1.38 | 8.78E-01 | Cdk1 | 1.48 | 3.38E-01 |
| C4b | 2.01 | 5.22E-04 | Il1rapl2 | 1.86 | 2.31E-01 | Il19 | 1.77 | 2.49E-02 | Epsti1 | 1.38 | 9.10E-01 | C9 | 1.48 | 7.82E-01 |
| C1qb | 2.00 | 8.64E-03 | Mx1 | 1.85 | 4.26E-01 | Ccr5 | 1.76 | 5.38E-04 | Tpsab1 | 1.37 | 9.10E-01 | Tcf7 | 1.47 | 8.42E-01 |
| C1qa | 1.96 | 7.78E-03 | Ccl25 | 1.78 | 6.77E-01 | Mbl2 | 1.74 | 2.89E-01 | Ambp | 1.36 | 8.78E-01 | Twist1 | 1.45 | 8.87E-01 |
| Mefv | 1.93 | 1.63E-01 | Col4a1 | 1.72 | 2.31E-04 | Thbs1 | 1.72 | 3.31E-03 | Ada | 1.33 | 9.90E-01 | Birc5 | 1.45 | 7.90E-01 |
| Rsad2 | 1.92 | 5.03E-02 | Ptgdr2 | 1.69 | 1.12E-02 | C3 | 1.71 | 3.16E-03 | Il22 | 1.32 | 8.78E-01 | Tigit | 1.44 | 4.97E-01 |
| C3 | 1.90 | 4.01E-02 | Cxcl1 | 1.69 | 3.68E-02 | Mefv | 1.71 | 3.13E-02 | Bst1 | 1.31 | 8.78E-01 | Klrb1 | 1.44 | 7.13E-01 |
| Dmbt1 | 1.86 | 6.10E-01 | Zbp1 | 1.69 | 1.36E-02 | Ccr2 | 1.70 | 1.10E-06 | Cdk1 | 1.31 | 8.78E-01 | Gfi1 | 1.44 | 7.82E-01 |
| Il24 | 1.86 | 3.00E-01 | Cxcl5 | 1.69 | 5.43E-02 | S100a8 | 1.69 | 9.79E-03 | C2 | 1.31 | 8.78E-01 | Tpsab1 | 1.42 | 7.19E-01 |
| Igll1 | 1.86 | 1.43E-01 | Aicda | 1.67 | 3.40E-01 | Cxcl5 | 1.69 | 1.33E-02 | Havcr2 | 1.30 | 8.96E-01 | Il10 | 1.41 | 4.94E-01 |
| Col4a1 | 1.85 | 9.25E-04 | Il2rb | 1.66 | 3.32E-03 | Cxcr1 | 1.67 | 8.48E-02 | Itgb3 | 1.29 | 8.78E-01 | Il17rb | 1.41 | 8.24E-01 |
| Cx3cr1 | 1.85 | 4.34E-03 | Cfb | 1.66 | 6.15E-02 | Prg2 | 1.67 | 8.03E-02 | Pmch | 1.29 | 9.90E-01 | Cd5 | 1.40 | 8.65E-01 |
| Cxcl11 | 1.83 | 4.51E-01 | Cd80 | 1.65 | 4.69E-02 | S100b | 1.66 | 3.12E-01 | Cxcl14 | 1.29 | 8.78E-01 | Il23r | 1.37 | 4.97E-01 |
| Il11 | 1.81 | 3.17E-01 | Ifit3 | 1.65 | 1.56E-03 | Ulbp1 | 1.64 | 3.13E-02 | Chil3 | 1.29 | 9.18E-01 | Klra5 | 1.37 | 7.16E-01 |
| Glycam1 | 1.78 | 7.17E-01 | Ulbp1 | 1.65 | 1.12E-01 | Cxcl1 | 1.63 | 9.79E-03 | Rsad2 | 1.29 | 8.78E-01 | C7 | 1.36 | 4.97E-01 |
| Klra3 | 1.77 | 1.90E-01 | Itgam | 1.64 | 7.24E-03 | Cxcl11 | 1.63 | 2.13E-01 | Ccl22 | 1.28 | 9.71E-01 | Il12rb1 | 1.36 | 7.82E-01 |
| Camp | 1.77 | 2.71E-01 | Ccl19 | 1.64 | 4.07E-02 | Zbp1 | 1.63 | 3.53E-03 | Tdo2 | 1.28 | 8.78E-01 | Igll1 | 1.36 | 6.11E-01 |
| Tnfrsf10b | 1.77 | 4.99E-03 | Rrad | 1.60 | 1.61E-02 | Pdcd1 | 1.61 | 1.62E-01 | Smpd3 | 1.28 | 9.90E-01 | Ada | 1.36 | 9.95E-01 |
| Mpo | 1.77 | 5.59E-01 | Fcer1g | 1.59 | 3.66E-04 | Ifit3 | 1.60 | 1.41E-04 | Vwf | 1.27 | 6.83E-01 | Notch1 | 1.35 | 1.43E-02 |
| Mpped1 | 1.74 | 6.41E-01 | Klra1 | 1.59 | 2.62E-01 | Ccl19 | 1.59 | 1.02E-02 | Fcgr4 | 1.26 | 8.78E-01 | Ccl25 | 1.35 | 9.49E-01 |
| Ptgs2 | 1.73 | 6.47E-02 | Tnfrsf1b | 1.59 | 4.57E-04 | Cxcl9 | 1.59 | 4.34E-02 | Clec4n | 1.26 | 8.78E-01 | Lck | 1.35 | 8.87E-01 |
| Il12rb1 | 1.71 | 5.36E-01 | Siglec1 | 1.58 | 8.57E-03 | Camp | 1.57 | 6.84E-02 | Gzmb | 1.26 | 8.78E-01 | Cd70 | 1.35 | 7.82E-01 |
| Pdcd1 | 1.70 | 4.59E-01 | Chil3 | 1.57 | 1.84E-01 | Rsad2 | 1.56 | 1.88E-02 | Ifit3 | 1.26 | 8.78E-01 | Atg16l1 | 1.34 | 5.83E-02 |
| Cxcl5 | 1.69 | 1.96E-01 | Ly86 | 1.57 | 6.82E-03 | Socs3 | 1.55 | 4.55E-03 | Lyz2 | 1.26 | 8.78E-01 | Fas | 1.34 | 8.05E-02 |
| Bax | 1.69 | 1.38E-04 | Socs3 | 1.56 | 2.38E-02 | Il11 | 1.52 | 1.69E-01 | Cxcr4 | 1.26 | 8.78E-01 | Ctsl | 1.34 | 5.03E-01 |
| Klra4 | 1.68 | 3.66E-01 | Plau | 1.56 | 5.32E-04 | Raet1 | 1.52 | 8.35E-03 | Cd84 | 1.26 | 8.78E-01 | Flt3l | 1.33 | 1.75E-01 |
| Il6 | 1.67 | 4.10E-01 | Sh2d1a | 1.56 | 6.75E-01 | Bax | 1.52 | 3.06E-06 | Msln | 1.26 | 8.78E-01 | Cd247 | 1.33 | 8.73E-01 |
| Ccr8 | 1.64 | 3.34E-01 | Tdo2 | 1.55 | 8.24E-02 | Siglec1 | 1.51 | 2.59E-03 | Prdm1 | 1.25 | 8.78E-01 | Foxp3 | 1.33 | 6.25E-01 |
| Ifnl2 | 1.64 | 2.47E-01 | Ccr1 | 1.55 | 2.10E-02 | Cd14 | 1.49 | 1.22E-04 | Lrrn3 | 1.25 | 8.78E-01 | Il17b | 1.33 | 6.11E-01 |
| Ulbp1 | 1.63 | 2.89E-01 | C3 | 1.54 | 7.08E-02 | Tnfrsf17 | 1.49 | 2.71E-01 | Fcer1g | 1.25 | 8.75E-01 | Fcer1a | 1.33 | 6.57E-01 |
| Lag3 | 1.62 | 1.92E-01 | Csf3r | 1.53 | 4.88E-02 | Fcer1g | 1.48 | 1.08E-04 | Abcg1 | 1.25 | 8.78E-01 | Ccl26 | 1.33 | 6.20E-01 |
| Clec5a | 1.61 | 7.90E-02 | Slc11a1 | 1.53 | 1.22E-02 | Mx1 | 1.47 | 4.63E-01 | Tnfsf14 | 1.24 | 8.78E-01 | Cd3e | 1.32 | 8.95E-01 |
| Raet1 | 1.61 | 1.01E-01 | Ifi44l | 1.52 | 5.89E-02 | Pdgfrb | 1.47 | 2.28E-05 | Cxcl10 | 1.24 | 9.90E-01 | Il11 | 1.32 | 7.57E-01 |
| Cxcl1 | 1.58 | 2.04E-01 | Cd7 | 1.51 | 3.83E-02 | Isg15 | 1.47 | 4.64E-03 | Gpr44 | 1.24 | 8.96E-01 | Bcl6 | 1.32 | 3.42E-01 |
| Tmem173 | 1.56 | 8.64E-03 | Pdcd1 | 1.51 | 3.80E-01 | Jak3 | 1.47 | 9.96E-05 | Csf2 | 1.24 | 8.78E-01 | Cd163 | 1.32 | 6.11E-01 |
| Nt5e | 1.56 | 3.96E-02 | Mefv | 1.51 | 2.40E-01 | Plau | 1.46 | 1.37E-04 | Ifna2 | 1.23 | 9.90E-01 | Ctla4 | 1.30 | 6.12E-01 |
| Ccl9 | 1.56 | 4.38E-02 | Il4 | 1.50 | 2.87E-01 | Igll1 | 1.46 | 1.04E-01 | Pdcd1 | 1.23 | 9.90E-01 | Lrrn3 | 1.30 | 3.26E-01 |
| Zbp1 | 1.56 | 1.48E-01 | Fn1 | 1.50 | 1.31E-02 | Ptgdr2 | 1.45 | 1.58E-02 | Fcgr1 | 1.23 | 8.78E-01 | Il1rapl2 | 1.30 | 8.26E-01 |

| | | | | | | | | | | | | | | |
|---|---|---|---|---|---|---|---|---|---|---|---|---|---|---|
| Fos | 1.56 | 7.06E-01 | Isg15 | 1.49 | 2.10E-02 | Ccl25 | 1.45 | 7.29E-01 | Hsd11b1 | 1.22 | 8.78E-01 | Nfkbia | 1.29 | 3.42E-01 |
| Ifit3 | 1.55 | 2.80E-02 | C2 | 1.49 | 1.24E-01 | Itgam | 1.45 | 8.59E-03 | Klra2 | 1.22 | 8.78E-01 | Cd244 | 1.28 | 6.20E-01 |
| Ccl19 | 1.55 | 1.96E-01 | Il13ra2 | 1.49 | 4.99E-01 | Tnfrsf10b | 1.45 | 1.82E-03 | Xcl1 | 1.22 | 9.18E-01 | Nfatc3 | 1.28 | 5.96E-01 |
| Socs3 | 1.54 | 1.37E-01 | Klra2 | 1.48 | 4.92E-02 | Il24 | 1.44 | 2.45E-01 | Il12b | 1.22 | 9.46E-01 | Ccl28 | 1.28 | 7.69E-01 |
| Jak3 | 1.53 | 6.03E-03 | Tnfrsf17 | 1.48 | 4.52E-01 | Ifi44l | 1.43 | 4.04E-02 | Card9 | 1.22 | 8.96E-01 | Klra15 | 1.28 | 8.13E-01 |
| Col3a1 | 1.52 | 2.82E-01 | Gzmb | 1.48 | 9.85E-02 | Tnfrsf1b | 1.43 | 3.19E-04 | Cd68 | 1.22 | 8.78E-01 | Zap70 | 1.28 | 8.42E-01 |
| Tnfsf15 | 1.51 | 1.37E-01 | Gzma | 1.48 | 8.24E-02 | Rag1 | 1.43 | 8.60E-01 | Il10 | 1.21 | 9.46E-01 | Atf1 | 1.27 | 1.35E-01 |
| H2-Q2 | 1.50 | 2.39E-01 | Cd14 | 1.47 | 2.39E-03 | Ccr1 | 1.42 | 1.44E-02 | F13a1 | 1.21 | 9.68E-01 | Pla2g6 | 1.27 | 1.75E-01 |
| Cd14 | 1.50 | 1.42E-02 | Gbp5 | 1.47 | 4.07E-02 | Ptgs2 | 1.42 | 3.72E-02 | Ccl6 | 1.21 | 9.10E-01 | Inpp5d | 1.27 | 8.66E-02 |
| C9 | 1.50 | 6.81E-01 | Tnfsf14 | 1.47 | 2.37E-02 | Cd99 | 1.42 | 2.22E-03 | C4b | 1.21 | 8.78E-01 | Mx1 | 1.27 | 8.95E-01 |
| Cxcl2 | 1.50 | 3.43E-01 | Pdgfrb | 1.46 | 5.81E-04 | Il2rb | 1.41 | 9.65E-03 | Il1rn | 1.21 | 8.78E-01 | Smpd3 | 1.27 | 9.05E-01 |
| Col1a1 | 1.50 | 1.47E-01 | Msr1 | 1.46 | 1.36E-02 | Bst2 | 1.40 | 1.37E-04 | Fut7 | 1.20 | 9.10E-01 | Cd96 | 1.26 | 8.85E-01 |
| Tnfrsf17 | 1.49 | 6.47E-01 | Cxcl11 | 1.46 | 5.34E-01 | Tnf | 1.40 | 1.71E-02 | Aicda | 1.20 | 9.90E-01 | Cd1d1 | 1.26 | 8.52E-01 |
| Foxj1 | 1.49 | 3.25E-01 | Oasl1 | 1.46 | 8.27E-02 | Mx2 | 1.40 | 6.85E-02 | Ccl28 | 1.20 | 9.90E-01 | Ikzf2 | 1.26 | 7.82E-01 |
| Ifna4 | 1.49 | 6.32E-01 | Hck | 1.45 | 6.83E-03 | H2-Q2 | 1.39 | 5.94E-02 | Nt5e | 1.20 | 8.78E-01 | Nup107 | 1.25 | 3.42E-01 |
| Nos2 | 1.48 | 9.82E-02 | Klrb1c | 1.45 | 1.08E-01 | Il12rb1 | 1.39 | 4.05E-01 | Il23r | 1.20 | 9.62E-01 | Itgb4 | 1.25 | 8.40E-01 |
| Pdgfrb | 1.48 | 4.99E-03 | Cd99 | 1.44 | 1.29E-02 | Csf1r | 1.39 | 2.32E-05 | Itgb2 | 1.19 | 8.78E-01 | Cfh | 1.25 | 4.75E-01 |
| Ccr4 | 1.47 | 6.61E-01 | Fcgr1 | 1.43 | 1.27E-02 | Ccr4 | 1.39 | 3.93E-01 | Lcn2 | 1.19 | 9.63E-01 | Cd4 | 1.25 | 9.36E-01 |
| Tnfrsf8 | 1.46 | 7.06E-01 | Raet1 | 1.43 | 8.13E-02 | Nt5e | 1.38 | 9.63E-03 | Ceacam1 | 1.19 | 8.84E-01 | Pdcd1lg2 | 1.25 | 7.50E-01 |
| Ccr5 | 1.46 | 2.13E-01 | Ccl4 | 1.42 | 2.87E-01 | Ifnl2 | 1.38 | 1.49E-01 | Cd200r1 | 1.18 | 8.78E-01 | Klra27 | 1.25 | 8.40E-01 |
| Cxcr1 | 1.46 | 6.14E-01 | Ncf4 | 1.42 | 6.83E-03 | Il3ra | 1.37 | 5.32E-04 | Ctsh | 1.18 | 8.78E-01 | Klra6 | 1.24 | 7.42E-01 |
| Oas2 | 1.45 | 3.42E-01 | Il4ra | 1.42 | 1.31E-03 | Msr1 | 1.37 | 8.59E-03 | Cdh1 | 1.18 | 8.78E-01 | Mme | 1.24 | 7.06E-01 |
| Il3ra | 1.45 | 1.46E-02 | Bst2 | 1.41 | 1.92E-03 | Hcst | 1.37 | 2.23E-01 | Ccl8 | 1.17 | 9.90E-01 | Myc | 1.24 | 8.42E-01 |
| Isg15 | 1.45 | 1.44E-01 | Il18rap | 1.41 | 6.15E-02 | C2 | 1.37 | 1.03E-01 | Lilra5 | 1.17 | 9.18E-01 | Elk1 | 1.24 | 6.65E-01 |
| Siglec1 | 1.44 | 1.46E-01 | Il1rl1 | 1.41 | 3.93E-02 | Il4ra | 1.37 | 1.83E-04 | Itgae | 1.17 | 8.96E-01 | Ewsr1 | 1.24 | 5.83E-02 |
| Clec4a2 | 1.44 | 1.27E-01 | Jak3 | 1.40 | 4.64E-03 | Tmem173 | 1.36 | 2.22E-03 | Hck | 1.17 | 8.78E-01 | Ltk | 1.24 | 7.82E-01 |
| Tnf | 1.43 | 1.76E-01 | Csf1r | 1.40 | 5.10E-04 | Vwf | 1.36 | 1.65E-03 | Klra15 | 1.17 | 9.90E-01 | Thy1 | 1.24 | 9.59E-01 |
| Klra6 | 1.42 | 4.37E-01 | Usp18 | 1.40 | 2.52E-02 | Il1rl1 | 1.36 | 1.37E-02 | C7 | 1.17 | 9.88E-01 | Ifng | 1.23 | 7.90E-01 |
| Ccr2 | 1.41 | 2.12E-02 | Mx2 | 1.39 | 1.77E-01 | F13a1 | 1.36 | 1.78E-01 | Cxcr6 | 1.16 | 9.35E-01 | Chil3 | 1.22 | 8.26E-01 |
| Vegfc | 1.41 | 6.99E-02 | Camp | 1.39 | 3.14E-01 | Runx3 | 1.36 | 5.94E-02 | Tnfsf13b | 1.16 | 8.84E-01 | Ilf3 | 1.22 | 3.26E-01 |
| Cd44 | 1.41 | 4.37E-01 | F13a1 | 1.39 | 2.95E-01 | Ly86 | 1.36 | 1.37E-02 | Bid | 1.16 | 9.90E-01 | Il9 | 1.22 | 8.34E-01 |
| Il2 | 1.40 | 4.43E-01 | Vwf | 1.39 | 1.05E-02 | Fn1 | 1.35 | 1.33E-02 | Il13 | 1.16 | 9.90E-01 | Arg2 | 1.21 | 6.03E-01 |
| Mx2 | 1.40 | 3.90E-01 | Klrg1 | 1.38 | 1.84E-01 | Fcgr2b | 1.35 | 2.39E-03 | Fcgr3 | 1.16 | 8.78E-01 | Sele | 1.21 | 8.74E-01 |
| Cd99 | 1.40 | 1.03E-01 | Thbs1 | 1.38 | 2.10E-01 | Egr1 | 1.35 | 4.99E-02 | Cd7 | 1.16 | 9.46E-01 | Egr2 | 1.21 | 7.90E-01 |
| Bst2 | 1.39 | 1.96E-02 | Tnf | 1.37 | 1.00E-01 | Col1a1 | 1.35 | 4.62E-02 | Mrc1 | 1.15 | 8.78E-01 | Il1r2 | 1.21 | 9.49E-01 |
| Csf1r | 1.38 | 6.39E-03 | Fcgr2b | 1.37 | 1.36E-02 | Slc11a1 | 1.34 | 2.23E-02 | Ulbp1 | 1.15 | 9.90E-01 | Cd44 | 1.21 | 7.90E-01 |
| Fcer1g | 1.38 | 5.08E-02 | Batf | 1.37 | 1.11E-01 | Oas2 | 1.34 | 1.21E-01 | Ctss | 1.15 | 9.46E-01 | Il17a | 1.21 | 8.44E-01 |
| Runx3 | 1.38 | 3.38E-01 | Hcst | 1.37 | 3.83E-01 | Tnfrsf8 | 1.33 | 5.04E-01 | Serping1 | 1.15 | 8.78E-01 | Spink5 | 1.20 | 9.85E-01 |
| Hcst | 1.38 | 6.12E-01 | Bax | 1.36 | 2.37E-03 | Fas | 1.33 | 4.33E-03 | Il18 | 1.15 | 9.10E-01 | Slamf6 | 1.20 | 8.63E-01 |
| Pmch | 1.37 | 7.69E-01 | Nlrp3 | 1.36 | 5.31E-02 | Ifi44 | 1.33 | 7.95E-02 | Syt17 | 1.15 | 9.88E-01 | F12 | 1.20 | 8.42E-01 |
| Cspg4 | 1.37 | 1.72E-01 | Csf1 | 1.36 | 4.70E-03 | Hck | 1.32 | 6.38E-03 | Lamp3 | 1.15 | 9.81E-01 | Il16 | 1.20 | 8.03E-01 |
| Plau | 1.37 | 5.18E-02 | Ido1 | 1.35 | 4.13E-01 | Tdo2 | 1.32 | 1.43E-01 | Tlr8 | 1.15 | 8.78E-01 | Cxcl13 | 1.20 | 9.77E-01 |
| Il17b | 1.36 | 5.68E-01 | Trem1 | 1.35 | 3.07E-01 | Cspg4 | 1.32 | 2.29E-02 | C1ra | 1.15 | 8.78E-01 | Ms4a2 | 1.20 | 8.49E-01 |
| Lbp | 1.36 | 1.66E-01 | Hamp | 1.35 | 8.24E-01 | Klra17 | 1.32 | 1.44E-01 | Lgals3 | 1.15 | 9.10E-01 | Csf2 | 1.20 | 6.77E-01 |
| Il23r | 1.36 | 5.18E-01 | Cd70 | 1.34 | 5.96E-01 | Tnfrsf11a | 1.32 | 1.59E-03 | C6 | 1.15 | 9.81E-01 | Klrb1c | 1.19 | 7.25E-01 |
| Ifi44l | 1.35 | 4.43E-01 | Fas | 1.34 | 2.16E-02 | Fcgr1 | 1.31 | 1.51E-02 | Abcb1a | 1.15 | 8.78E-01 | Ccnd3 | 1.19 | 8.34E-01 |
| Mnx1 | 1.35 | 6.91E-01 | Runx3 | 1.34 | 2.00E-01 | Col3a1 | 1.31 | 1.80E-01 | Itga2 | 1.14 | 9.10E-01 | H2-Q10 | 1.19 | 8.34E-01 |
| H2-DMb1 | 1.34 | 6.96E-01 | Klra17 | 1.33 | 2.70E-01 | Vegfc | 1.31 | 1.12E-02 | Cd9 | 1.14 | 9.46E-01 | Il15ra | 1.19 | 7.42E-01 |
| Fcgr2b | 1.34 | 1.03E-01 | Entpd1 | 1.33 | 2.18E-03 | Aicda | 1.30 | 5.08E-01 | Itga6 | 1.14 | 8.78E-01 | Gzmm | 1.18 | 7.82E-01 |
| Vwf | 1.34 | 9.39E-02 | Tnfrsf11b | 1.32 | 1.47E-01 | Lcn2 | 1.30 | 2.22E-01 | Cd38 | 1.14 | 9.10E-01 | Itk | 1.18 | 9.49E-01 |
| Tnfrsf1a | 1.34 | 6.03E-03 | Il1b | 1.32 | 4.13E-01 | Il13ra2 | 1.29 | 5.69E-01 | Usp18 | 1.14 | 8.96E-01 | Epsti1 | 1.18 | 8.87E-01 |
| Ifi44 | 1.33 | 3.99E-01 | Itgb3 | 1.32 | 1.15E-01 | Entpd1 | 1.29 | 3.12E-04 | Ccrl2 | 1.14 | 8.78E-01 | Plaur | 1.18 | 4.88E-01 |
| F13a1 | 1.33 | 6.05E-01 | Ifi44 | 1.32 | 2.05E-01 | Rrad | 1.29 | 9.23E-02 | Tnfrsf8 | 1.14 | 9.90E-01 | Cd59b | 1.18 | 8.10E-01 |
| Nfkb2 | 1.33 | 3.48E-02 | Tnfrsf11a | 1.31 | 1.38E-02 | Gbp5 | 1.29 | 8.03E-02 | Cfi | 1.14 | 9.90E-01 | Ccl6 | 1.18 | 7.82E-01 |
| F12 | 1.33 | 6.49E-01 | Lcn2 | 1.31 | 3.71E-01 | Il12b | 1.28 | 2.66E-01 | Ifit1 | 1.14 | 9.46E-01 | Ccl11 | 1.18 | 9.66E-01 |
| Ltf | 1.33 | 4.92E-01 | Csf2rb | 1.31 | 1.19E-02 | Sbno2 | 1.28 | 3.64E-04 | H2-Q2 | 1.14 | 9.90E-01 | Colec12 | 1.17 | 6.61E-01 |
| Ambp | 1.33 | 5.64E-01 | Sbno2 | 1.31 | 2.53E-03 | Csf3r | 1.28 | 1.56E-01 | Muc1 | 1.13 | 9.35E-01 | Mr1 | 1.17 | 3.42E-01 |
| Gata3 | 1.33 | 4.43E-01 | Ccr4 | 1.31 | 6.44E-01 | Il23r | 1.28 | 2.54E-01 | Alcam | 1.13 | 9.10E-01 | Cd27 | 1.17 | 9.78E-01 |
| Il4ra | 1.32 | 4.00E-02 | Klra21 | 1.30 | 5.69E-01 | Itgb3 | 1.28 | 6.51E-02 | Lta | 1.13 | 9.90E-01 | Dock9 | 1.17 | 4.89E-01 |
| Tnfrsf11a | 1.32 | 6.18E-02 | Snai1 | 1.30 | 3.47E-01 | Ncf4 | 1.27 | 1.34E-02 | Icam1 | 1.13 | 9.73E-01 | Cx3cl1 | 1.17 | 5.68E-01 |
| Il1rl1 | 1.32 | 2.76E-01 | Il3ra | 1.30 | 2.10E-02 | Csf1 | 1.27 | 3.15E-03 | Il22ra1 | 1.13 | 9.90E-01 | Yy1 | 1.16 | 1.43E-01 |
| Ccr1 | 1.31 | 3.85E-01 | Emr1 | 1.30 | 7.05E-02 | Gata3 | 1.27 | 1.85E-01 | Txnip | 1.13 | 8.96E-01 | Creb5 | 1.16 | 7.82E-01 |
| Fas | 1.31 | 1.40E-01 | H2-Q2 | 1.29 | 3.14E-01 | Cxcl2 | 1.27 | 2.73E-01 | Anxa1 | 1.13 | 8.78E-01 | Tnfsf18 | 1.16 | 8.85E-01 |
| Ccl20 | 1.30 | 7.11E-01 | Cklf | 1.28 | 3.21E-01 | Emr1 | 1.27 | 3.27E-02 | Cx3cl1 | 1.12 | 8.96E-01 | Cd8b1 | 1.16 | 9.92E-01 |
| Klra17 | 1.30 | 5.65E-01 | Gpr44 | 1.28 | 3.44E-01 | Tnfrsf1a | 1.27 | 2.25E-04 | Il12rb1 | 1.12 | 9.90E-01 | Lag3 | 1.16 | 8.42E-01 |
| S100a8 | 1.30 | 6.05E-01 | Il11 | 1.28 | 6.03E-01 | Foxj1 | 1.26 | 2.64E-01 | Cybb | 1.12 | 9.81E-01 | Tlr7 | 1.16 | 7.82E-01 |

| | | | | | | | | | | | | | | |
|---|---|---|---|---|---|---|---|---|---|---|---|---|---|---|
| Masp1 | 1.29 | 4.86E-01 | Msln | 1.28 | 3.71E-01 | Clec4a2 | 1.26 | 6.67E-02 | Tgfb1 | 1.12 | 8.78E-01 | Slc7a11 | 1.16 | 9.34E-01 |
| Msr1 | 1.29 | 3.06E-01 | Cspg4 | 1.28 | 1.47E-01 | Nfkb2 | 1.26 | 2.87E-03 | Pla2g1b | 1.12 | 9.90E-01 | Il7 | 1.16 | 7.82E-01 |
| Il12b | 1.29 | 6.43E-01 | Il12b | 1.28 | 4.34E-01 | Ccl20 | 1.26 | 4.48E-01 | Il21 | 1.11 | 9.90E-01 | Cd160 | 1.15 | 8.63E-01 |
| Lcn2 | 1.29 | 6.21E-01 | Tyk2 | 1.27 | 1.23E-01 | Oasl1 | 1.26 | 2.04E-01 | Angpt1 | 1.11 | 8.96E-01 | Il22ra2 | 1.15 | 8.85E-01 |
| Ifng | 1.28 | 6.73E-01 | Psmb10 | 1.27 | 3.83E-02 | Batf | 1.25 | 1.43E-01 | Ifna4 | 1.11 | 9.90E-01 | C8g | 1.15 | 9.20E-01 |
| Tnfrsf1b | 1.28 | 1.48E-01 | Cfp | 1.27 | 2.70E-01 | Usp18 | 1.25 | 4.91E-02 | Vegfc | 1.11 | 9.21E-01 | Cd1d2 | 1.15 | 8.42E-01 |
| Tap2 | 1.28 | 2.77E-01 | Il10ra | 1.27 | 5.43E-02 | Card9 | 1.25 | 2.14E-01 | Thbd | 1.11 | 9.90E-01 | Dll4 | 1.15 | 6.25E-01 |
| Ifit1 | 1.28 | 4.22E-01 | Tnfaip3 | 1.27 | 1.74E-01 | Klra4 | 1.24 | 4.75E-01 | Mr1 | 1.11 | 8.78E-01 | Il1rl2 | 1.15 | 7.90E-01 |
| Itgam | 1.28 | 4.15E-01 | Prg2 | 1.27 | 6.18E-01 | Nos2 | 1.24 | 1.02E-01 | Itgb4 | 1.11 | 9.90E-01 | Rora | 1.15 | 6.61E-01 |
| Mcam | 1.27 | 1.44E-01 | Ccl1 | 1.26 | 5.28E-01 | Psmb10 | 1.24 | 1.37E-02 | Ifitm2 | 1.11 | 8.78E-01 | Il34 | 1.15 | 5.42E-01 |
| Vcam1 | 1.27 | 3.14E-01 | Rsad2 | 1.26 | 4.13E-01 | Tnfaip3 | 1.24 | 8.27E-02 | Trem1 | 1.11 | 9.90E-01 | Ctsh | 1.15 | 7.25E-01 |
| Card9 | 1.26 | 5.77E-01 | Tnfsf4 | 1.26 | 5.43E-01 | Dmbt1 | 1.24 | 7.29E-01 | Isg20 | 1.11 | 8.78E-01 | Il6ra | 1.15 | 6.11E-01 |
| Entpd1 | 1.26 | 5.03E-02 | Cxcl3 | 1.25 | 6.44E-01 | Il10ra | 1.24 | 2.33E-02 | H2-Q1 | 1.11 | 9.90E-01 | Mif | 1.15 | 8.40E-01 |
| C2 | 1.26 | 6.32E-01 | Abca1 | 1.25 | 9.85E-02 | Il4 | 1.24 | 4.63E-01 | Il19 | 1.11 | 9.90E-01 | Bst1 | 1.15 | 7.82E-01 |
| Sbno2 | 1.25 | 5.42E-02 | Cd276 | 1.25 | 7.89E-02 | Ifna1 | 1.24 | 4.63E-01 | Snai1 | 1.10 | 9.90E-01 | Cebpb | 1.15 | 8.42E-01 |
| Ptgdr2 | 1.25 | 5.66E-01 | Ccl24 | 1.25 | 6.43E-01 | Trem1 | 1.23 | 3.44E-01 | Zfp13 | 1.10 | 9.72E-01 | Ikzf1 | 1.15 | 9.34E-01 |
| Zfp13 | 1.24 | 3.38E-01 | Cxcl9 | 1.25 | 5.34E-01 | Csf2rb | 1.23 | 1.06E-02 | Csf2rb | 1.10 | 8.96E-01 | Creb1 | 1.14 | 3.42E-01 |
| Ifna1 | 1.24 | 7.69E-01 | Irf7 | 1.25 | 3.52E-01 | Klra2 | 1.22 | 2.23E-01 | Tnfsf12 | 1.10 | 9.48E-01 | Clec4a2 | 1.14 | 7.25E-01 |
| Itgb3 | 1.24 | 4.59E-01 | H2-Q1 | 1.25 | 5.83E-01 | Cklf | 1.22 | 2.64E-01 | Sh2d1b1 | 1.10 | 9.90E-01 | Jun | 1.14 | 7.90E-01 |
| Pvr | 1.23 | 1.30E-01 | Tirap | 1.24 | 8.42E-02 | Ido1 | 1.22 | 4.75E-01 | Rora | 1.10 | 9.73E-01 | Foxj1 | 1.14 | 8.63E-01 |
| Emr1 | 1.23 | 3.85E-01 | Fcgr4 | 1.24 | 3.97E-01 | Zfp13 | 1.22 | 7.45E-02 | Cxcl16 | 1.10 | 9.90E-01 | F13a1 | 1.14 | 8.65E-01 |
| Vim | 1.23 | 1.90E-01 | Ptprc | 1.24 | 3.85E-01 | Il1b | 1.22 | 4.44E-01 | Abca1 | 1.10 | 9.46E-01 | Il17ra | 1.13 | 7.54E-01 |
| Vhl | 1.23 | 2.49E-01 | Gpr183 | 1.24 | 3.00E-01 | Irf7 | 1.22 | 2.64E-01 | Csf3 | 1.10 | 9.90E-01 | Pvr | 1.13 | 4.88E-01 |
| Cd2 | 0.63 | 3.94E-01 | Cd5 | 0.67 | 6.53E-01 | Cd40 | 0.67 | 6.05E-04 | Cd3g | 0.77 | 9.90E-01 | Klrg1 | 0.76 | 4.97E-01 |
| Defb1 | 0.62 | 8.62E-01 | Fos | 0.67 | 5.50E-01 | Cd53 | 0.66 | 2.92E-02 | Ecsit | 0.76 | 8.78E-01 | Klrd1 | 0.76 | 3.42E-01 |
| Slamf6 | 0.62 | 4.37E-01 | Cr2 | 0.66 | 1.36E-02 | Ccl27a | 0.66 | 2.51E-03 | Osm | 0.76 | 8.78E-01 | H2-Ab1 | 0.76 | 3.42E-01 |
| Klrc2 | 0.62 | 9.39E-02 | Atg16l1 | 0.66 | 5.98E-04 | Slamf1 | 0.66 | 1.02E-01 | Rel | 0.76 | 2.11E-01 | Il18rap | 0.75 | 3.26E-01 |
| Cd53 | 0.60 | 1.48E-01 | H2-DMb2 | 0.66 | 3.83E-02 | Ly9 | 0.66 | 1.04E-01 | Ly9 | 0.76 | 8.96E-01 | Tlr2 | 0.75 | 1.19E-01 |
| Cd8b1 | 0.60 | 8.34E-01 | Chit1 | 0.66 | 7.82E-01 | Serpinb2 | 0.65 | 5.03E-01 | Gzmm | 0.75 | 8.78E-01 | Chit1 | 0.75 | 9.78E-01 |
| Il9 | 0.60 | 3.33E-01 | Cd36 | 0.65 | 3.27E-02 | Cd37 | 0.65 | 2.59E-03 | Ccl25 | 0.75 | 9.90E-01 | H2-Q2 | 0.75 | 4.94E-01 |
| Mme | 0.59 | 1.25E-01 | Defb1 | 0.65 | 7.70E-01 | Cxcl14 | 0.65 | 2.44E-02 | Il15ra | 0.74 | 8.78E-01 | Gpr183 | 0.75 | 3.26E-01 |
| Cxcr5 | 0.59 | 1.42E-01 | H2-Ob | 0.65 | 2.78E-02 | Cmah | 0.64 | 2.70E-03 | Il2 | 0.74 | 8.78E-01 | Tnfsf10 | 0.74 | 3.38E-01 |
| Tnfrsf4 | 0.58 | 2.71E-01 | Cd9 | 0.63 | 1.55E-02 | Flt3l | 0.64 | 1.50E-04 | Ifi27 | 0.74 | 9.03E-01 | C3ar1 | 0.74 | 2.17E-01 |
| Ly9 | 0.58 | 3.11E-01 | Cxcl14 | 0.63 | 7.12E-02 | Defb1 | 0.64 | 6.61E-01 | C8b | 0.74 | 8.78E-01 | Ccr7 | 0.74 | 2.46E-01 |
| Mef2c | 0.58 | 1.48E-01 | Cmah | 0.62 | 1.21E-02 | Ltb | 0.63 | 2.18E-02 | Cd27 | 0.74 | 9.90E-01 | Fcer2a | 0.74 | 5.04E-01 |
| Cr2 | 0.58 | 1.27E-02 | Slamf6 | 0.61 | 2.00E-01 | Il16 | 0.63 | 2.82E-02 | Cd2 | 0.73 | 8.78E-01 | Il1b | 0.73 | 6.11E-01 |
| Cd37 | 0.58 | 2.81E-02 | C8g | 0.61 | 2.28E-01 | Cr2 | 0.62 | 3.33E-04 | Runx3 | 0.72 | 8.78E-01 | Il3 | 0.72 | 7.54E-01 |
| Cfh | 0.57 | 1.46E-02 | Arg1 | 0.60 | 6.04E-01 | Slamf6 | 0.62 | 8.45E-02 | Il1b | 0.72 | 8.78E-01 | Fcgr1 | 0.72 | 8.33E-02 |
| Sell | 0.57 | 6.12E-02 | Pparg | 0.60 | 3.69E-01 | Cd3e | 0.62 | 4.40E-01 | Ccl20 | 0.72 | 8.96E-01 | Nos2 | 0.72 | 2.17E-01 |
| Twist1 | 0.56 | 7.10E-01 | Cfh | 0.59 | 2.78E-03 | Cd1d1 | 0.62 | 1.44E-01 | Tnfrsf4 | 0.71 | 8.78E-01 | Tnfrsf12a | 0.72 | 1.35E-01 |
| Ltb | 0.56 | 1.30E-01 | Cd22 | 0.59 | 6.83E-03 | Ccr7 | 0.62 | 3.34E-04 | Egr3 | 0.71 | 8.78E-01 | Xcr1 | 0.72 | 6.27E-01 |
| Cd3g | 0.56 | 7.17E-01 | Xcr1 | 0.58 | 7.81E-02 | Itga2b | 0.61 | 9.23E-02 | Ccr7 | 0.71 | 5.74E-01 | Ido1 | 0.71 | 6.11E-01 |
| Spink5 | 0.55 | 7.06E-01 | Sh2b2 | 0.58 | 3.79E-01 | Cd79a | 0.61 | 3.72E-02 | Masp2 | 0.71 | 6.83E-01 | Ccl12 | 0.71 | 3.42E-01 |
| Il16 | 0.55 | 1.40E-01 | Cd19 | 0.58 | 1.21E-02 | H2-DMb2 | 0.60 | 1.48E-03 | Il17f | 0.71 | 8.78E-01 | Cd7 | 0.70 | 2.17E-01 |
| Ccr7 | 0.55 | 8.55E-03 | Mef2c | 0.57 | 4.03E-02 | Masp2 | 0.60 | 7.69E-04 | Thy1 | 0.70 | 9.90E-01 | Tnfrsf13c | 0.70 | 4.88E-01 |
| Arg1 | 0.55 | 6.97E-01 | Jun | 0.57 | 4.64E-03 | Cfh | 0.58 | 1.37E-04 | Saa1 | 0.70 | 9.46E-01 | Ciita | 0.69 | 2.17E-01 |
| Cd27 | 0.55 | 7.06E-01 | Mme | 0.57 | 1.72E-02 | Mme | 0.58 | 3.15E-03 | Slamf6 | 0.70 | 8.78E-01 | Hamp | 0.69 | 9.04E-01 |
| H2-DMb2 | 0.55 | 2.80E-02 | Flt3l | 0.57 | 3.72E-04 | Mef2c | 0.58 | 7.93E-03 | Il23a | 0.70 | 8.78E-01 | Ccr2 | 0.69 | 8.15E-03 |
| Cd79a | 0.55 | 1.77E-01 | Il1r2 | 0.56 | 3.90E-01 | Arg1 | 0.58 | 3.97E-01 | Tnfrsf10b | 0.70 | 5.74E-01 | Gzma | 0.69 | 2.73E-01 |
| Cd22 | 0.54 | 1.67E-02 | Btla | 0.54 | 2.69E-03 | Cd22 | 0.56 | 2.69E-04 | Mnx1 | 0.68 | 8.78E-01 | Cr2 | 0.68 | 1.22E-01 |
| Cd3e | 0.54 | 6.94E-01 | Egr3 | 0.54 | 8.87E-02 | H2-Ob | 0.56 | 2.89E-04 | H60a | 0.68 | 8.78E-01 | Cxcl3 | 0.67 | 6.25E-01 |
| Cd70 | 0.53 | 3.99E-01 | Masp2 | 0.54 | 1.90E-03 | Spink5 | 0.55 | 3.60E-01 | Lck | 0.67 | 9.90E-01 | Pou2f2 | 0.65 | 1.03E-01 |
| Tnfrsf13c | 0.53 | 1.01E-01 | Spink5 | 0.54 | 5.52E-01 | Chit1 | 0.54 | 5.22E-01 | Timd4 | 0.67 | 8.78E-01 | Ms4a1 | 0.65 | 1.75E-01 |
| Il5ra | 0.53 | 1.33E-01 | C7 | 0.53 | 1.27E-02 | Blk | 0.54 | 8.32E-04 | Cd4 | 0.67 | 9.90E-01 | Ccl7 | 0.65 | 1.72E-01 |
| Serpinb2 | 0.50 | 6.05E-01 | Il12a | 0.51 | 2.54E-03 | Cxcr5 | 0.53 | 1.84E-03 | Cd69 | 0.66 | 8.78E-01 | Ccr5 | 0.64 | 8.05E-02 |
| Il12a | 0.50 | 1.17E-02 | Slamf1 | 0.50 | 5.25E-02 | Il1r2 | 0.52 | 2.02E-01 | Cd3e | 0.66 | 9.90E-01 | Cd180 | 0.64 | 2.75E-02 |
| Il1r2 | 0.49 | 5.64E-01 | Pax5 | 0.50 | 3.46E-03 | Cd5 | 0.52 | 2.67E-01 | Mpo | 0.66 | 9.10E-01 | Klra1 | 0.64 | 5.11E-01 |
| H2-Ob | 0.48 | 6.03E-03 | Itgae | 0.49 | 1.06E-03 | Ppbp | 0.52 | 3.87E-02 | Cd8b1 | 0.64 | 9.90E-01 | Cd19 | 0.63 | 1.75E-01 |
| Itgae | 0.48 | 6.97E-03 | Cxcr5 | 0.48 | 7.19E-03 | Twist1 | 0.51 | 2.97E-01 | Rag1 | 0.64 | 9.90E-01 | H2-Ob | 0.63 | 1.22E-01 |
| C7 | 0.48 | 2.97E-02 | Ada | 0.48 | 8.06E-01 | Il12a | 0.50 | 1.08E-04 | Pparg | 0.64 | 8.96E-01 | Ncr1 | 0.63 | 6.55E-02 |
| Chit1 | 0.45 | 7.11E-01 | Mpped1 | 0.47 | 4.27E-01 | C7 | 0.50 | 6.38E-04 | Cd6 | 0.63 | 9.46E-01 | Ccl4 | 0.62 | 3.36E-01 |
| Blk | 0.43 | 7.78E-03 | Twist1 | 0.46 | 4.34E-01 | Xcr1 | 0.49 | 3.20E-03 | Sele | 0.62 | 8.78E-01 | Il6 | 0.62 | 4.53E-01 |
| Pax5 | 0.41 | 6.03E-03 | Cd1d1 | 0.45 | 6.15E-02 | Itgae | 0.49 | 4.28E-05 | Ccr3 | 0.61 | 9.90E-01 | Ccl5 | 0.62 | 6.55E-02 |
| Xcr1 | 0.41 | 3.91E-02 | Itga2b | 0.45 | 4.98E-02 | Cd19 | 0.49 | 1.06E-04 | Cd8a | 0.60 | 9.90E-01 | C8b | 0.61 | 3.26E-01 |
| Cd5 | 0.41 | 4.80E-01 | A2m | 0.44 | 6.42E-02 | Pax5 | 0.45 | 8.41E-05 | Cd1d1 | 0.60 | 8.78E-01 | S100a8 | 0.60 | 1.91E-01 |
| Cd19 | 0.41 | 3.69E-03 | Cd207 | 0.44 | 1.79E-03 | Btla | 0.43 | 5.57E-06 | Cfd | 0.60 | 9.90E-01 | C8a | 0.58 | 8.15E-01 |

| | | | | | | | | | | | | | | |
|---|---|---|---|---|---|---|---|---|---|---|---|---|---|---|
| Timd4 | 0.39 | 1.44E-01 | Cd79b | 0.44 | 1.81E-05 | Timd4 | 0.41 | 1.12E-02 | S100b | 0.57 | 9.18E-01 | Cxcl5 | 0.58 | 1.38E-01 |
| Btla | 0.34 | 2.96E-04 | Ms4a1 | 0.44 | 1.91E-04 | Cd207 | 0.39 | 2.32E-05 | C8g | 0.55 | 8.78E-01 | Lif | 0.57 | 2.61E-04 |
| Cd207 | 0.34 | 3.06E-03 | Timd4 | 0.43 | 7.71E-02 | Cd79b | 0.39 | 9.29E-08 | Ccr9 | 0.55 | 9.90E-01 | Rrad | 0.57 | 2.75E-02 |
| Cd79b | 0.34 | 9.03E-05 | Fcer2a | 0.41 | 2.72E-04 | Fcer2a | 0.37 | 3.81E-06 | Ccl26 | 0.51 | 5.74E-01 | Prf1 | 0.56 | 2.75E-02 |
| Fcer2a | 0.34 | 1.17E-03 | Ppbp | 0.38 | 3.27E-02 | Ms4a1 | 0.37 | 5.81E-07 | Fos | 0.43 | 8.78E-01 | Cx3cr1 | 0.53 | 6.94E-04 |
| Ms4a1 | 0.31 | 1.38E-04 | Glycam1 | 0.37 | 2.41E-01 | Cfd | 0.31 | 3.67E-01 | Glycam1 | 0.41 | 8.78E-01 | Cxcl10 | 0.48 | 6.55E-02 |
| Ada | 0.17 | 6.75E-01 | Cfd | 0.15 | 3.39E-01 | Ada | 0.28 | 4.93E-01 | Sh2b2 | 0.33 | 8.78E-01 | Ccl2 | 0.47 | 8.15E-03 |

Supplementary Table 2: Top 150 & bottom 50 NanoString PanCancer Pathways panel hits ranked by fold change (FC)

| Fig. 2e Vehicle vs Chemo | | | Fig. 8b Vehicle vs Chemo | | | Fig. 2e+8b Veh. vs Chemo | | | Fig. 8c Vehicle ± Nav | | | Fig. 8c Chemo ± Nav | | |
|---|---|---|---|---|---|---|---|---|---|---|---|---|---|---|
| Factor | FC | adj.P.Val | Factor | FC | adj.P.Val | Factor | FC | adj.P.Val | Factor | FC | adj.P.Val | Factor | FC | adj.P.Val |
| Cdkn1a | 4.96 | 7.97E-05 | Cdkn1a | 3.98 | 4.10E-05 | Cdkn1a | 4.44 | 2.67E-07 | Zbtb16 | 5.38 | 7.21E-02 | Zbtb16 | 9.45 | 2.95E-03 |
| Mmp3 | 4.02 | 2.32E-02 | Tnc | 3.53 | 1.96E-03 | Tnc | 3.68 | 5.60E-05 | Ifnb1 | 3.33 | 8.02E-01 | Wee1 | 1.83 | 2.95E-03 |
| Tnc | 3.85 | 1.04E-02 | Lif | 2.81 | 4.10E-05 | Mmp3 | 3.13 | 9.88E-04 | Nodal | 2.29 | 8.02E-01 | Bcl2l1 | 1.73 | 2.95E-03 |
| Ifnb1 | 3.33 | 3.61E-01 | Pla1a | 2.52 | 7.45E-02 | Lif | 2.58 | 2.62E-06 | Gata1 | 2.27 | 1.14E-01 | Fgf3 | 1.66 | 7.92E-01 |
| Ccna2 | 2.81 | 4.87E-01 | Mmp3 | 2.44 | 3.47E-02 | Ifnb1 | 2.41 | 1.32E-01 | Rasgrf1 | 1.95 | 8.02E-01 | Sox9 | 1.63 | 4.91E-01 |
| Pla2g5 | 2.66 | 2.11E-01 | Etv4 | 2.36 | 1.40E-02 | Il6 | 2.35 | 3.67E-03 | Mpl | 1.79 | 8.02E-01 | Comp | 1.62 | 8.24E-01 |
| Nr4a1 | 2.63 | 1.16E-01 | Il6 | 2.26 | 3.05E-02 | Ccna2 | 2.27 | 1.36E-01 | Wnt10a | 1.78 | 8.02E-01 | Ddb2 | 1.60 | 2.40E-01 |
| Il6 | 2.45 | 1.19E-01 | Pax3 | 2.21 | 2.40E-02 | Brip1 | 2.05 | 8.17E-03 | Dll3 | 1.72 | 8.04E-01 | Alk | 1.58 | 8.68E-01 |
| Lif | 2.37 | 4.65E-03 | Hist1h3b | 2.15 | 2.75E-01 | Hist1h3b | 2.04 | 1.73E-01 | Pitx2 | 1.63 | 8.02E-01 | Figf | 1.57 | 1.05E-01 |
| Ttk | 2.33 | 5.38E-01 | Cxcl5 | 2.09 | 7.84E-03 | Col5a1 | 2.03 | 5.51E-05 | Creb3l3 | 1.62 | 8.04E-01 | Ccnb1 | 1.54 | 7.24E-01 |
| Inhba | 2.32 | 8.22E-01 | Wnt3a | 2.06 | 1.96E-03 | Etv4 | 2.00 | 6.29E-03 | Il19 | 1.61 | 8.02E-01 | Hspa1a | 1.54 | 9.24E-01 |
| Nr4a3 | 2.31 | 1.71E-01 | Col5a1 | 2.01 | 1.96E-03 | Thbs1 | 1.97 | 2.15E-03 | Amh | 1.60 | 8.02E-01 | Fst | 1.53 | 6.31E-01 |
| Thbs1 | 2.25 | 4.96E-02 | Brip1 | 2.01 | 5.47E-02 | Ttk | 1.93 | 1.90E-01 | Fgf20 | 1.59 | 8.02E-01 | Map3k12 | 1.53 | 4.91E-01 |
| Top2a | 2.18 | 7.04E-01 | Fgf23 | 1.99 | 2.53E-01 | Inhba | 1.90 | 4.64E-01 | Hnf1a | 1.57 | 8.02E-01 | Fgf20 | 1.52 | 7.14E-01 |
| Brip1 | 2.10 | 2.01E-01 | Gadd45g | 1.98 | 2.00E-02 | Top2a | 1.89 | 2.96E-01 | Ccnb1 | 1.56 | 8.02E-01 | Ccne2 | 1.52 | 8.68E-01 |
| Cxxc4 | 2.05 | 1.38E-01 | Nodal | 1.93 | 3.05E-01 | Pla1a | 1.84 | 1.26E-01 | Rps27a | 1.54 | 8.04E-01 | Cacna1h | 1.51 | 7.92E-01 |
| Col5a1 | 2.05 | 1.04E-02 | Pgf | 1.88 | 1.16E-03 | Dll3 | 1.83 | 1.90E-01 | Pla2g4c | 1.54 | 8.02E-01 | Fgf14 | 1.48 | 9.09E-01 |
| Col24a1 | 2.03 | 2.85E-01 | Csf3r | 1.87 | 2.99E-02 | Nodal | 1.77 | 2.51E-01 | Grin2a | 1.53 | 8.02E-01 | Cntfr | 1.46 | 6.94E-01 |
| Lama1 | 1.97 | 2.40E-01 | Cxcl1 | 1.86 | 5.72E-02 | Igf1 | 1.75 | 1.09E-02 | Nog | 1.52 | 8.79E-01 | Epo | 1.46 | 8.68E-01 |
| Igf1 | 1.95 | 1.19E-01 | Ccna2 | 1.84 | 4.15E-01 | Gdf6 | 1.71 | 3.76E-02 | Prmt8 | 1.50 | 8.02E-01 | Pla2g5 | 1.46 | 8.88E-01 |
| Hist1h3b | 1.95 | 7.29E-01 | Nog | 1.83 | 3.86E-01 | Lama1 | 1.71 | 4.32E-02 | Birc7 | 1.49 | 8.02E-01 | Il2ra | 1.45 | 9.09E-01 |
| Dll3 | 1.85 | 7.04E-01 | Dll3 | 1.81 | 3.25E-01 | Cxcl1 | 1.70 | 3.33E-02 | Gng4 | 1.49 | 8.02E-01 | Ptcra | 1.45 | 9.41E-01 |
| Chad | 1.84 | 5.55E-01 | Gdf6 | 1.77 | 9.06E-02 | Pla2g4c | 1.67 | 5.16E-02 | Alk | 1.48 | 8.18E-01 | Mpl | 1.44 | 8.38E-01 |
| Cdkn2a | 1.82 | 2.06E-01 | Ppp2r2c | 1.74 | 1.01E-01 | Col27a1 | 1.65 | 3.64E-03 | Sfrp2 | 1.47 | 8.02E-01 | Il4 | 1.43 | 8.42E-01 |
| Col2a1 | 1.79 | 6.28E-01 | Sfrp2 | 1.74 | 1.04E-01 | Pax3 | 1.65 | 6.03E-02 | Ppp3r2 | 1.44 | 8.02E-01 | Il1r2 | 1.43 | 9.31E-01 |
| Il11 | 1.78 | 4.17E-01 | Pla2g4c | 1.74 | 1.15E-01 | Flna | 1.64 | 4.90E-04 | Hoxa10 | 1.42 | 8.04E-01 | Il22ra2 | 1.42 | 8.38E-01 |
| Wnt5a | 1.77 | 1.09E-03 | Ifnb1 | 1.74 | 5.12E-01 | Smc1b | 1.62 | 8.29E-02 | Pla2g10 | 1.42 | 9.32E-01 | Hist2h3b | 1.42 | 9.31E-01 |
| Wnt11 | 1.75 | 5.24E-02 | Thbs1 | 1.72 | 5.01E-02 | Pgf | 1.62 | 6.45E-04 | Wnt3a | 1.42 | 8.02E-01 | Fzd10 | 1.40 | 9.31E-01 |
| Ifna1 | 1.72 | 5.55E-01 | Pdgfrb | 1.71 | 4.95E-03 | Rad51 | 1.61 | 1.93E-01 | Tnr | 1.40 | 8.04E-01 | Ttk | 1.39 | 9.31E-01 |
| Etv4 | 1.70 | 3.61E-01 | Pla2g2a | 1.71 | 2.92E-02 | Cdc25c | 1.61 | 3.15E-01 | Lefty2 | 1.40 | 8.08E-01 | Ezh2 | 1.39 | 8.68E-01 |
| Ptcra | 1.69 | 8.71E-01 | Rad51 | 1.71 | 2.81E-01 | Cxcl5 | 1.59 | 2.94E-02 | Pax3 | 1.40 | 8.04E-01 | Il11ra1 | 1.39 | 7.92E-01 |
| Fgf8 | 1.67 | 4.98E-01 | Cdc25c | 1.71 | 4.05E-01 | Il23a | 1.59 | 3.56E-02 | Il2 | 1.39 | 8.08E-01 | Igfbp3 | 1.38 | 8.05E-01 |
| Col27a1 | 1.66 | 1.19E-01 | Flna | 1.70 | 4.58E-03 | Epha2 | 1.58 | 3.76E-03 | Wt1 | 1.37 | 8.02E-01 | Gria3 | 1.38 | 4.91E-01 |
| Il23a | 1.65 | 3.28E-01 | Il2rb | 1.70 | 8.31E-03 | Wnt3a | 1.58 | 4.19E-03 | Tnn | 1.37 | 8.02E-01 | Rad51 | 1.37 | 8.88E-01 |
| Smc1b | 1.65 | 5.10E-01 | Fzd2 | 1.68 | 5.10E-03 | Fgf8 | 1.56 | 1.04E-01 | Wnt3 | 1.36 | 8.04E-01 | Gngt1 | 1.36 | 8.68E-01 |
| Gdf6 | 1.65 | 4.65E-01 | Epha2 | 1.66 | 1.73E-02 | Wnt5a | 1.56 | 3.94E-05 | Casp9 | 1.36 | 5.75E-01 | Brca2 | 1.36 | 7.39E-01 |
| Spry4 | 1.64 | 4.96E-02 | Il13 | 1.66 | 2.69E-01 | Sfrp2 | 1.56 | 8.37E-02 | Fgf15 | 1.36 | 8.04E-01 | Cxxc4 | 1.36 | 7.94E-01 |
| Nodal | 1.63 | 8.19E-01 | Plau | 1.65 | 2.82E-04 | Col5a2 | 1.56 | 1.08E-02 | Ntrk1 | 1.35 | 8.02E-01 | Cdk2 | 1.36 | 4.91E-01 |
| Col5a2 | 1.61 | 1.98E-01 | Creb3l3 | 1.64 | 4.13E-01 | Wnt11 | 1.55 | 4.19E-03 | Zic2 | 1.35 | 8.17E-01 | Mmp7 | 1.36 | 8.68E-01 |
| E2f1 | 1.61 | 2.02E-01 | Col27a1 | 1.64 | 2.59E-02 | Fzd2 | 1.54 | 1.91E-03 | Cacng6 | 1.34 | 9.45E-01 | Rps6ka6 | 1.35 | 8.42E-01 |
| Pla2g4c | 1.60 | 5.38E-01 | Top2a | 1.63 | 5.56E-01 | Col24a1 | 1.53 | 1.55E-01 | Cacng4 | 1.33 | 8.12E-01 | Lef1 | 1.35 | 9.09E-01 |
| Socs3 | 1.60 | 9.79E-02 | Wnt7a | 1.63 | 1.10E-02 | Socs3 | 1.52 | 3.67E-03 | Smc1b | 1.33 | 8.17E-01 | Hist1h3b | 1.35 | 9.43E-01 |
| Jak3 | 1.59 | 1.04E-02 | Lama5 | 1.62 | 2.59E-02 | Prmt8 | 1.51 | 6.54E-02 | Wnt2b | 1.33 | 8.04E-01 | Fas | 1.34 | 3.81E-01 |
| Flna | 1.59 | 7.05E-02 | Grin2a | 1.62 | 9.60E-02 | Pdgfrb | 1.51 | 3.64E-03 | Lama3 | 1.33 | 8.02E-01 | Myb | 1.34 | 9.31E-01 |
| Tnfrsf10b | 1.58 | 3.89E-02 | Notch3 | 1.62 | 4.67E-03 | Epor | 1.51 | 3.64E-03 | Pla2g4e | 1.33 | 9.55E-01 | Fgf13 | 1.34 | 9.41E-01 |
| Col1a1 | 1.58 | 9.93E-02 | Epor | 1.60 | 9.94E-03 | Il11 | 1.51 | 1.58E-01 | Efna3 | 1.32 | 9.45E-01 | Klf4 | 1.34 | 7.35E-01 |
| Cacna1h | 1.58 | 6.74E-01 | Smc1b | 1.59 | 2.20E-01 | Brca1 | 1.51 | 3.30E-01 | Gdf6 | 1.32 | 8.08E-01 | Prmt8 | 1.33 | 7.92E-01 |
| Fos | 1.58 | 8.22E-01 | Inhbb | 1.59 | 2.65E-01 | Bax | 1.48 | 2.10E-04 | Wnt1 | 1.32 | 8.79E-01 | Cacng6 | 1.33 | 9.45E-01 |
| Ccne2 | 1.57 | 7.78E-01 | Ttk | 1.59 | 5.12E-01 | Il3ra | 1.48 | 1.91E-03 | Flt1 | 1.31 | 8.02E-01 | Stat4 | 1.33 | 6.94E-01 |
| Bax | 1.57 | 1.19E-02 | Fn1 | 1.58 | 1.36E-02 | Plau | 1.47 | 1.73E-04 | Fzd9 | 1.31 | 9.26E-01 | Cdc25c | 1.33 | 9.31E-01 |
| Cxcl1 | 1.56 | 5.38E-01 | Igf1 | 1.58 | 1.26E-01 | Ddit4 | 1.46 | 2.32E-02 | Calml3 | 1.31 | 9.78E-01 | Cdc7 | 1.32 | 8.68E-01 |
| Prom1 | 1.56 | 3.31E-01 | Il3ra | 1.58 | 6.51E-03 | Hist2h3b | 1.46 | 5.10E-01 | Cacnb4 | 1.31 | 8.04E-01 | Ccnb3 | 1.32 | 9.21E-01 |
| Chek2 | 1.54 | 3.61E-01 | Fgf15 | 1.57 | 2.08E-01 | Chad | 1.46 | 3.43E-01 | Nfe2l2 | 1.31 | 8.02E-01 | Wnt2b | 1.32 | 8.73E-01 |
| Myb | 1.54 | 8.41E-01 | Inhba | 1.55 | 7.26E-01 | Jak3 | 1.45 | 4.90E-04 | Bcl2a1a | 1.31 | 8.02E-01 | Rasal1 | 1.31 | 8.68E-01 |
| Brca1 | 1.53 | 8.05E-01 | Prmt8 | 1.54 | 1.58E-01 | Cdkn2a | 1.44 | 1.05E-01 | Hoxa11 | 1.29 | 9.45E-01 | Mcm7 | 1.31 | 8.68E-01 |
| Rad51 | 1.52 | 7.60E-01 | Fgf20 | 1.54 | 2.91E-01 | Crlf2 | 1.43 | 1.88E-04 | Cacng1 | 1.29 | 9.66E-01 | Ccno | 1.31 | 4.91E-01 |
| Cdc25c | 1.51 | 8.41E-01 | Il23a | 1.53 | 1.30E-01 | Il2rb | 1.43 | 2.17E-02 | Il20rb | 1.29 | 9.27E-01 | Il20rb | 1.31 | 9.31E-01 |
| Col6a6 | 1.51 | 7.60E-01 | Lama3 | 1.52 | 2.71E-02 | Dtx4 | 1.42 | 3.64E-03 | Gzmb | 1.29 | 8.04E-01 | Col4a6 | 1.30 | 8.06E-01 |
| Fgf17 | 1.51 | 7.37E-01 | Col5a2 | 1.51 | 7.72E-02 | Nupr1 | 1.42 | 1.55E-02 | Wee1 | 1.28 | 8.02E-01 | Arid2 | 1.29 | 8.68E-01 |
| Epha2 | 1.51 | 2.03E-01 | Wt1 | 1.51 | 2.07E-01 | Cd14 | 1.42 | 1.02E-03 | Ret | 1.28 | 8.30E-01 | Rps6ka5 | 1.29 | 5.72E-01 |
| Ptprr | 1.50 | 1.92E-01 | Creb3l1 | 1.51 | 7.45E-02 | Tnfrsf10b | 1.42 | 3.67E-03 | Wnt7b | 1.28 | 9.45E-01 | H2afx | 1.29 | 9.10E-01 |

| | | | | | | | | | | | | | | |
|---|---|---|---|---|---|---|---|---|---|---|---|---|---|---|
| Prmt8 | 1.49 | 5.38E-01 | Cdkn1c | 1.50 | 4.67E-03 | Mdm2 | 1.41 | 2.09E-04 | Rac3 | 1.28 | 8.02E-01 | Lamc3 | 1.28 | 8.85E-01 |
| Chek1 | 1.49 | 7.66E-01 | Csf1r | 1.50 | 7.53E-03 | Fgf15 | 1.41 | 2.17E-01 | Bambi | 1.27 | 8.02E-01 | Whsc1 | 1.28 | 8.45E-01 |
| Shc3 | 1.49 | 7.67E-01 | Hist2h3b | 1.50 | 6.16E-01 | Col2a1 | 1.41 | 3.73E-01 | Csf2rb | 1.27 | 8.02E-01 | Il11ra2 | 1.28 | 9.31E-01 |
| Ddit4 | 1.49 | 2.71E-01 | Smo | 1.49 | 3.49E-02 | Fgf23 | 1.40 | 4.92E-01 | Gadd45g | 1.26 | 8.17E-01 | Fgfr4 | 1.28 | 8.68E-01 |
| Fosl1 | 1.48 | 4.87E-01 | Brca1 | 1.49 | 4.74E-01 | Col1a2 | 1.40 | 1.04E-02 | Ifng | 1.26 | 9.15E-01 | Birc7 | 1.27 | 8.77E-01 |
| Nupr1 | 1.48 | 1.98E-01 | Crlf2 | 1.48 | 1.96E-03 | Lama5 | 1.40 | 3.69E-02 | Wnt4 | 1.25 | 9.32E-01 | Notch1 | 1.27 | 4.91E-01 |
| Casp12 | 1.48 | 5.59E-02 | Lama1 | 1.48 | 2.77E-01 | Gadd45g | 1.40 | 1.35E-01 | Efna1 | 1.25 | 8.08E-01 | Bnip3 | 1.27 | 8.88E-01 |
| Ccna1 | 1.47 | 6.74E-01 | Birc7 | 1.47 | 2.55E-01 | Csf1r | 1.40 | 3.64E-03 | Ccnb3 | 1.25 | 9.27E-01 | Baiap3 | 1.27 | 7.39E-01 |
| Fzd9 | 1.46 | 8.41E-01 | Cacng4 | 1.47 | 3.05E-01 | Csf3 | 1.40 | 4.78E-01 | Dll1 | 1.25 | 8.02E-01 | Ret | 1.26 | 9.09E-01 |
| Shc2 | 1.44 | 5.38E-01 | Col4a4 | 1.46 | 2.26E-02 | Fanca | 1.39 | 4.48E-02 | Fzd2 | 1.25 | 8.02E-01 | Chek1 | 1.26 | 9.31E-01 |
| Ccne1 | 1.43 | 8.41E-01 | Fgf8 | 1.46 | 2.93E-01 | Fen1 | 1.39 | 1.03E-01 | Itgb4 | 1.25 | 9.04E-01 | Col11a2 | 1.26 | 9.31E-01 |
| Rasgrf1 | 1.43 | 8.27E-01 | Ppp3r2 | 1.45 | 3.31E-01 | Ifna1 | 1.38 | 3.31E-01 | Fzd10 | 1.24 | 9.45E-01 | Col11a1 | 1.26 | 9.31E-01 |
| Epor | 1.42 | 2.23E-01 | Csf2rb | 1.45 | 1.73E-02 | Ppp3r2 | 1.38 | 2.58E-01 | Smo | 1.24 | 8.02E-01 | Ets2 | 1.26 | 6.00E-01 |
| Hist2h3b | 1.42 | 9.00E-01 | Hist2h3c2 | 1.45 | 1.60E-02 | Col4a4 | 1.38 | 1.06E-02 | Il3ra | 1.24 | 8.02E-01 | Stmn1 | 1.26 | 8.68E-01 |
| H2afx | 1.42 | 7.86E-01 | Itgb6 | 1.44 | 1.04E-01 | Csf3r | 1.37 | 1.54E-01 | Creb5 | 1.24 | 8.02E-01 | Brca1 | 1.25 | 9.31E-01 |
| Arnt2 | 1.42 | 4.87E-01 | Ddit4 | 1.44 | 1.08E-01 | Rps27a | 1.37 | 4.59E-01 | Il11 | 1.23 | 9.04E-01 | Pkmyt1 | 1.25 | 9.02E-01 |
| Fanca | 1.42 | 3.61E-01 | Socs3 | 1.44 | 4.67E-02 | Wnt7a | 1.37 | 3.70E-02 | Pias1 | 1.23 | 8.02E-01 | Pdgfra | 1.25 | 8.06E-01 |
| Mdm2 | 1.41 | 3.05E-02 | Tpo | 1.43 | 2.82E-01 | Cacng4 | 1.37 | 3.06E-01 | Csf2 | 1.23 | 8.17E-01 | Socs2 | 1.25 | 7.39E-01 |
| Dtx4 | 1.41 | 1.45E-01 | Fen1 | 1.43 | 1.82E-01 | E2f1 | 1.36 | 6.86E-02 | Dkk4 | 1.22 | 8.18E-01 | Top2a | 1.25 | 9.45E-01 |
| Reln | 1.41 | 6.74E-01 | Dtx4 | 1.43 | 1.93E-02 | Lama3 | 1.36 | 3.26E-02 | Itgb3 | 1.22 | 8.02E-01 | Ccne1 | 1.25 | 9.31E-01 |
| Col3a1 | 1.41 | 5.44E-01 | Cd14 | 1.43 | 7.24E-03 | Col1a1 | 1.36 | 2.81E-02 | Fgfr2 | 1.22 | 8.02E-01 | Dkk1 | 1.25 | 9.02E-01 |
| Cd14 | 1.41 | 8.93E-02 | Gzmb | 1.42 | 2.13E-01 | Chek1 | 1.36 | 4.08E-01 | Cebpe | 1.22 | 9.15E-01 | Creb3l3 | 1.25 | 9.45E-01 |
| Mcm5 | 1.41 | 7.60E-01 | Rps27a | 1.42 | 5.12E-01 | Casp12 | 1.36 | 4.81E-03 | Mapk8ip1 | 1.22 | 8.02E-01 | Dnmt1 | 1.24 | 7.92E-01 |
| Fzd2 | 1.40 | 2.11E-01 | Casp9 | 1.42 | 1.93E-02 | Chek2 | 1.35 | 1.42E-01 | Il13ra2 | 1.22 | 9.45E-01 | Srsf2 | 1.24 | 3.35E-01 |
| Col1a2 | 1.40 | 2.22E-01 | Ngfr | 1.42 | 2.75E-01 | Fosl1 | 1.35 | 1.58E-01 | Smad2 | 1.21 | 8.02E-01 | Id1 | 1.24 | 5.50E-01 |
| Sfrp2 | 1.40 | 7.29E-01 | Hoxa10 | 1.42 | 4.15E-01 | H2afx | 1.35 | 3.81E-01 | Pdgfra | 1.21 | 8.12E-01 | Il23r | 1.24 | 8.68E-01 |
| Il3ra | 1.39 | 1.88E-01 | Csf2 | 1.42 | 2.08E-01 | Itgb6 | 1.35 | 7.84E-02 | Prkar1b | 1.21 | 8.02E-01 | Amh | 1.24 | 9.34E-01 |
| Pgf | 1.39 | 2.41E-01 | Mdm2 | 1.41 | 4.21E-03 | Rasgrf1 | 1.35 | 4.66E-01 | Fgf17 | 1.21 | 9.32E-01 | Npm1 | 1.23 | 8.38E-01 |
| Cdc6 | 1.39 | 8.15E-01 | Zic2 | 1.41 | 4.13E-01 | Wt1 | 1.34 | 2.37E-01 | Kit | 1.21 | 8.08E-01 | Ccna2 | 1.23 | 9.45E-01 |
| Csf3 | 1.39 | 8.78E-01 | Bax | 1.41 | 8.66E-03 | Shc4 | 1.34 | 2.68E-01 | Rad51 | 1.21 | 9.45E-01 | Il7 | 1.23 | 8.68E-01 |
| Crlf2 | 1.39 | 4.96E-02 | Hoxa11 | 1.41 | 6.42E-01 | Fn1 | 1.34 | 3.56E-02 | Ccna1 | 1.21 | 9.04E-01 | Six1 | 1.22 | 9.45E-01 |
| Vegfc | 1.38 | 1.52E-01 | Csf3 | 1.41 | 6.03E-01 | Notch3 | 1.33 | 1.73E-02 | Hhip | 1.20 | 8.02E-01 | Il10 | 1.22 | 9.31E-01 |
| Ccnd1 | 1.38 | 4.36E-01 | Fzd8 | 1.40 | 1.57E-01 | Nr4a3 | 1.33 | 3.59E-01 | Il12b | 1.20 | 9.32E-01 | Acvr1c | 1.22 | 9.45E-01 |
| Dll1 | 1.38 | 2.03E-01 | Hmga1 | 1.40 | 2.77E-01 | Myb | 1.33 | 5.72E-01 | Idh2 | 1.20 | 9.04E-01 | Itga8 | 1.22 | 8.68E-01 |
| Prl | 1.37 | 7.07E-01 | Col1a2 | 1.40 | 5.47E-02 | Prom1 | 1.33 | 1.73E-01 | Wnt2 | 1.20 | 8.30E-01 | Fgf22 | 1.22 | 9.02E-01 |
| Hnf1a | 1.36 | 8.41E-01 | Fasl | 1.39 | 3.13E-01 | Pla2g2a | 1.32 | 1.56E-01 | Cacna2d2 | 1.20 | 9.45E-01 | Cacna1c | 1.22 | 8.42E-01 |
| Pla1a | 1.34 | 8.84E-01 | Shc4 | 1.39 | 3.56E-01 | Cdc6 | 1.32 | 4.08E-01 | Lama5 | 1.20 | 8.08E-01 | Cdc25a | 1.22 | 8.38E-01 |
| Fen1 | 1.34 | 6.74E-01 | Ntrk1 | 1.39 | 3.01E-01 | Ptprr | 1.32 | 5.26E-02 | Dusp5 | 1.19 | 8.02E-01 | Hdac11 | 1.21 | 4.91E-01 |
| Creb3l4 | 1.34 | 6.53E-01 | Wnt5a | 1.38 | 1.10E-02 | Inhbb | 1.32 | 4.07E-01 | Bap1 | 1.19 | 8.04E-01 | Sos1 | 1.21 | 6.78E-01 |
| Pdgfrb | 1.33 | 3.61E-01 | Il13ra2 | 1.37 | 6.29E-01 | Ccnd1 | 1.32 | 8.75E-02 | Itga8 | 1.19 | 8.37E-01 | Med12 | 1.20 | 4.91E-01 |
| Il23r | 1.33 | 7.04E-01 | Wnt11 | 1.37 | 1.15E-01 | Hells | 1.31 | 4.36E-01 | Pdgfrb | 1.19 | 8.04E-01 | Blm | 1.20 | 9.09E-01 |
| Grin2b | 1.33 | 8.05E-01 | Fanca | 1.37 | 1.62E-01 | Ccne1 | 1.31 | 5.07E-01 | Polr2j | 1.19 | 8.04E-01 | Wnt2 | 1.20 | 9.02E-01 |
| Fgf16 | 1.33 | 8.57E-01 | Cacna2d4 | 1.36 | 2.04E-01 | Mcm2 | 1.30 | 3.43E-01 | Ccno | 1.19 | 8.02E-01 | Rasgrp2 | 1.20 | 4.91E-01 |
| Il3 | 1.32 | 8.41E-01 | Nupr1 | 1.36 | 1.08E-01 | Creb3l1 | 1.30 | 1.20E-01 | Csf3r | 1.18 | 9.04E-01 | Acvr2a | 1.20 | 8.68E-01 |
| Epo | 1.32 | 8.78E-01 | Mfng | 1.36 | 4.98E-02 | Zic2 | 1.30 | 4.12E-01 | Mfng | 1.18 | 8.02E-01 | Pias1 | 1.20 | 8.67E-01 |
| Ccno | 1.32 | 3.60E-01 | Cebpe | 1.35 | 4.61E-01 | Gadd45a | 1.30 | 1.58E-03 | Lamc3 | 1.18 | 9.24E-01 | Fance | 1.20 | 6.16E-01 |
| Rps27a | 1.32 | 8.90E-01 | Hells | 1.34 | 5.33E-01 | Pkmyt1 | 1.30 | 3.51E-01 | Ddit3 | 1.18 | 8.02E-01 | Fancf | 1.19 | 4.91E-01 |
| Plau | 1.31 | 1.45E-01 | Gata3 | 1.34 | 3.05E-01 | Csf2rb | 1.29 | 2.81E-02 | Csf1r | 1.18 | 8.02E-01 | Ube2t | 1.19 | 9.45E-01 |
| Pkmyt1 | 1.31 | 8.14E-01 | Ccnb1 | 1.34 | 5.12E-01 | Jag2 | 1.29 | 1.01E-01 | Lefty1 | 1.18 | 8.02E-01 | Lig4 | 1.19 | 9.09E-01 |
| Ccnd2 | 1.31 | 1.16E-01 | Prkcg | 1.34 | 1.18E-01 | Hmga1 | 1.29 | 2.91E-01 | Dnmt3a | 1.18 | 8.04E-01 | Dvl2 | 1.19 | 5.18E-01 |
| Ppp3r2 | 1.31 | 8.21E-01 | Wnt10a | 1.34 | 5.66E-01 | Cdkn1c | 1.29 | 1.51E-02 | Brip1 | 1.17 | 9.35E-01 | Fbxw7 | 1.19 | 6.75E-01 |
| Il20ra | 1.31 | 9.14E-01 | Bambi | 1.34 | 5.29E-02 | Ccnd2 | 1.28 | 3.76E-03 | Akt3 | 1.17 | 8.02E-01 | Il12b | 1.19 | 9.45E-01 |
| Mcm2 | 1.30 | 8.15E-01 | Hsp90b1 | 1.34 | 5.17E-03 | Nog | 1.28 | 6.61E-01 | Prdm1 | 1.17 | 8.02E-01 | Msh6 | 1.19 | 9.02E-01 |
| Csf1r | 1.30 | 2.85E-01 | Tnf | 1.34 | 2.99E-01 | Hnf1a | 1.28 | 4.99E-01 | Erbb3 | 1.17 | 8.02E-01 | Atrx | 1.19 | 5.50E-01 |
| Shc4 | 1.30 | 8.09E-01 | Il24 | 1.33 | 6.44E-01 | Birc7 | 1.28 | 3.50E-01 | Csf3 | 1.17 | 9.55E-01 | Il5ra | 1.19 | 9.45E-01 |
| Col4a4 | 1.30 | 3.79E-01 | Jag2 | 1.33 | 1.82E-01 | Ppp2r2c | 1.28 | 4.06E-01 | Hdac4 | 1.17 | 8.02E-01 | Kmt2c | 1.19 | 4.91E-01 |
| Alk | 1.29 | 9.14E-01 | Rfc4 | 1.32 | 4.13E-01 | Il13 | 1.28 | 5.09E-01 | Pold1 | 1.17 | 8.79E-01 | Pcna | 1.19 | 9.24E-01 |
| Casp3 | 1.29 | 4.97E-01 | Jak3 | 1.32 | 3.19E-02 | Fasl | 1.28 | 3.50E-01 | Ptch1 | 1.17 | 8.02E-01 | Cacna1d | 1.19 | 8.88E-01 |
| Gadd45a | 1.29 | 1.08E-01 | Bmp8a | 1.32 | 4.07E-01 | Pla2g5 | 1.28 | 5.56E-01 | Il12rb2 | 1.17 | 9.04E-01 | Plcb1 | 1.18 | 7.48E-01 |
| Il2 | 1.29 | 8.62E-01 | Bcl2a1a | 1.31 | 2.15E-01 | Ptcra | 1.27 | 7.45E-01 | Il23a | 1.17 | 9.15E-01 | Il6ra | 1.18 | 7.39E-01 |
| Ret | 1.29 | 8.41E-01 | Pim1 | 1.31 | 4.15E-01 | Mfng | 1.27 | 3.79E-02 | Ltbp1 | 1.17 | 8.02E-01 | Lat | 1.18 | 9.56E-01 |
| Pdgfa | 1.29 | 3.28E-01 | Gadd45a | 1.31 | 9.80E-03 | Fgf17 | 1.27 | 4.94E-01 | Notch3 | 1.16 | 8.04E-01 | Rasgrf1 | 1.18 | 9.45E-01 |
| Hells | 1.28 | 8.78E-01 | Skp2 | 1.31 | 4.65E-01 | Bambi | 1.27 | 3.26E-02 | Bmp6 | 1.16 | 8.24E-01 | Angpt1 | 1.18 | 7.92E-01 |
| Bdnf | 1.28 | 7.04E-01 | Il19 | 1.31 | 6.44E-01 | Fzd8 | 1.26 | 2.08E-01 | Cdk4 | 1.16 | 8.04E-01 | Gli1 | 1.18 | 8.42E-01 |
| Blm | 1.28 | 7.60E-01 | Mpo | 1.30 | 5.04E-01 | Smo | 1.26 | 9.71E-02 | Figf | 1.16 | 8.12E-01 | Mcm2 | 1.17 | 9.31E-01 |
| Plat | 1.28 | 3.60E-01 | Mcm2 | 1.30 | 4.99E-01 | Plat | 1.26 | 3.96E-02 | Creb3l1 | 1.15 | 8.79E-01 | Eya1 | 1.17 | 9.04E-01 |

| | | | | | | | | | | | | | | |
|---|---|---|---|---|---|---|---|---|---|---|---|---|---|---|
| Pak3 | 1.28 | 8.57E-01 | Trp53 | 1.30 | 6.25E-02 | Epo | 1.26 | 5.66E-01 | Dll4 | 1.15 | 8.17E-01 | Cacng1 | 1.17 | 9.80E-01 |
| Fgf9 | 1.27 | 7.70E-01 | Mapk8ip1 | 1.29 | 2.04E-01 | Casp3 | 1.26 | 9.51E-02 | Igf1 | 1.15 | 9.32E-01 | Smc1a | 1.17 | 5.69E-01 |
| Pik3r3 | 1.27 | 3.31E-01 | Ngf | 1.29 | 6.35E-01 | Nr4a1 | 1.26 | 4.83E-01 | Lamb3 | 1.15 | 8.04E-01 | Pik3r5 | 1.17 | 9.02E-01 |
| Cacng4 | 1.27 | 8.62E-01 | H2afx | 1.28 | 6.09E-01 | Pdgfa | 1.26 | 3.82E-02 | Itga6 | 1.15 | 8.02E-01 | Prkar2b | 1.16 | 9.56E-01 |
| Fgf15 | 1.27 | 8.41E-01 | Pkmyt1 | 1.28 | 5.12E-01 | Dll1 | 1.26 | 5.26E-02 | Igf1r | 1.15 | 8.04E-01 | Cdc6 | 1.16 | 9.45E-01 |
| Hhip | 1.27 | 5.57E-01 | Il11 | 1.28 | 5.56E-01 | Spry4 | 1.26 | 8.13E-02 | Gadd45b | 1.15 | 8.04E-01 | Nf1 | 1.16 | 4.91E-01 |
| Jag2 | 1.26 | 6.74E-01 | Socs1 | 1.27 | 6.11E-01 | Rfc4 | 1.26 | 3.81E-01 | Eya1 | 1.15 | 8.92E-01 | Il12rb2 | 1.16 | 9.29E-01 |
| Itgb6 | 1.26 | 7.04E-01 | Rasgrf1 | 1.27 | 7.09E-01 | Arnt2 | 1.26 | 2.43E-01 | Inhbb | 1.15 | 9.45E-01 | Idh2 | 1.16 | 9.31E-01 |
| Col11a2 | 1.25 | 9.14E-01 | Tnfrsf10b | 1.27 | 1.30E-01 | Gata3 | 1.25 | 3.15E-01 | Acvr1b | 1.15 | 8.02E-01 | Fgfr3 | 1.16 | 8.57E-01 |
| Itgb3 | 1.24 | 5.72E-01 | Bmp6 | 1.27 | 2.82E-01 | Vegfc | 1.24 | 4.59E-02 | Nkd1 | 1.15 | 8.08E-01 | Wnt7b | 1.16 | 9.66E-01 |
| Lamc2 | 1.24 | 6.97E-01 | Spry1 | 1.26 | 2.13E-01 | Tnfaip3 | 1.24 | 1.17E-01 | Pold4 | 1.14 | 8.02E-01 | Plcb4 | 1.15 | 6.20E-01 |
| Pax3 | 1.23 | 8.78E-01 | Sgk2 | 1.26 | 8.38E-01 | Il13ra2 | 1.24 | 6.62E-01 | Bmp7 | 1.14 | 9.27E-01 | Ddit4 | 1.15 | 9.02E-01 |
| Fgf18 | 1.23 | 7.04E-01 | Ccnd1 | 1.26 | 2.95E-01 | Hist2h3c2 | 1.24 | 5.59E-02 | Cacnb3 | 1.14 | 8.08E-01 | Kmt2d | 1.15 | 6.94E-01 |
| Tnfaip3 | 1.22 | 6.74E-01 | Ccnd2 | 1.26 | 3.92E-02 | Il19 | 1.24 | 6.10E-01 | Epo | 1.14 | 9.55E-01 | Dtx3 | 1.15 | 7.39E-01 |
| Gpc4 | 1.22 | 6.73E-01 | Tnfaip3 | 1.26 | 2.13E-01 | Ccne2 | 1.24 | 6.40E-01 | Lamc2 | 1.14 | 8.64E-01 | Endog | 1.15 | 9.21E-01 |
| Numbl | 1.22 | 3.28E-01 | Gata1 | 1.26 | 5.12E-01 | Ngfr | 1.23 | 4.08E-01 | Fzd8 | 1.14 | 9.15E-01 | Gata2 | 1.15 | 9.02E-01 |
| Col4a3 | 1.22 | 7.60E-01 | B2m | 1.25 | 9.68E-02 | Hoxa11 | 1.23 | 6.92E-01 | Etv4 | 1.14 | 9.45E-01 | Sf3b1 | 1.15 | 8.38E-01 |
| Lama3 | 1.22 | 7.04E-01 | Cdc6 | 1.25 | 6.44E-01 | Trp53 | 1.23 | 4.46E-02 | Angpt1 | 1.14 | 8.08E-01 | Hmga2 | 1.15 | 9.31E-01 |
| Wnt3a | 1.22 | 7.37E-01 | Stk4 | 1.25 | 1.35E-01 | Hhip | 1.23 | 1.42E-01 | Lama1 | 1.14 | 9.45E-01 | Mpo | 1.15 | 9.45E-01 |
| Wnt3 | 1.21 | 8.88E-01 | Nkd1 | 1.25 | 1.87E-01 | Il2 | 1.23 | 5.39E-01 | Spry1 | 1.14 | 8.29E-01 | Skp2 | 1.15 | 9.43E-01 |
| Lama5 | 1.21 | 7.60E-01 | Fgf11 | 1.25 | 2.76E-01 | Prkcg | 1.23 | 1.58E-01 | Il1r2 | 1.14 | 9.66E-01 | Whsc1l1 | 1.15 | 7.39E-01 |
| Wee1 | 1.21 | 6.01E-01 | Casp12 | 1.24 | 1.23E-01 | Fgf9 | 1.23 | 3.67E-01 | Plat | 1.14 | 8.04E-01 | Mlh1 | 1.15 | 8.21E-01 |
| Cxcl5 | 1.21 | 8.41E-01 | Plat | 1.24 | 1.42E-01 | Il24 | 1.22 | 6.62E-01 | Cdkn1c | 1.14 | 8.04E-01 | Tslp | 1.14 | 9.31E-01 |
| Arid2 | 0.67 | 6.13E-01 | Fgf1 | 0.69 | 7.29E-02 | Col11a1 | 0.70 | 3.39E-01 | Col11a1 | 0.77 | 9.04E-01 | Flnc | 0.77 | 9.45E-01 |
| Wnt6 | 0.66 | 6.28E-01 | Npm1 | 0.68 | 9.06E-02 | Ets2 | 0.70 | 6.29E-03 | Cxxc4 | 0.77 | 8.04E-01 | Nupr1 | 0.77 | 5.50E-01 |
| Prkcb | 0.66 | 7.70E-01 | Ccr7 | 0.67 | 6.41E-02 | Sost | 0.69 | 6.50E-01 | Dusp8 | 0.77 | 8.02E-01 | Fgf18 | 0.76 | 5.50E-01 |
| Il20rb | 0.66 | 8.18E-01 | Cxxc4 | 0.67 | 2.05E-01 | Stat4 | 0.69 | 4.86E-02 | Il23r | 0.77 | 8.02E-01 | Fgf6 | 0.76 | 8.88E-01 |
| Creb3l3 | 0.65 | 8.22E-01 | Eif4ebp1 | 0.67 | 1.49E-01 | Bmp7 | 0.69 | 7.79E-02 | Tnfrsf10b | 0.76 | 8.02E-01 | Col1a1 | 0.76 | 4.91E-01 |
| Dtx1 | 0.65 | 3.34E-01 | Six1 | 0.67 | 5.62E-01 | Lef1 | 0.69 | 3.39E-01 | Mapk10 | 0.76 | 9.45E-01 | Tpo | 0.76 | 8.68E-01 |
| Cd40 | 0.65 | 5.24E-02 | Mpl | 0.66 | 3.03E-01 | Sos1 | 0.69 | 1.74E-03 | Socs3 | 0.76 | 8.02E-01 | Pla2g10 | 0.76 | 9.45E-01 |
| Egf | 0.64 | 8.62E-01 | Thbs4 | 0.66 | 5.80E-01 | Fgf10 | 0.68 | 4.87E-03 | Jun | 0.76 | 8.02E-01 | Tgfb3 | 0.76 | 2.34E-01 |
| Il12b | 0.64 | 7.23E-01 | Pla2g3 | 0.66 | 6.45E-01 | Itga8 | 0.68 | 4.09E-02 | Acvr1c | 0.75 | 9.27E-01 | Cdkn2b | 0.76 | 6.94E-01 |
| Wnt4 | 0.64 | 7.60E-01 | Fos | 0.65 | 5.04E-01 | Jun | 0.68 | 2.32E-02 | Lep | 0.75 | 9.04E-01 | Ngfr | 0.76 | 8.42E-01 |
| Lefty2 | 0.63 | 7.04E-01 | Fgf10 | 0.64 | 1.11E-02 | Il12b | 0.68 | 2.76E-01 | Vegfb | 0.74 | 8.02E-01 | Ntrk1 | 0.75 | 8.42E-01 |
| Col11a1 | 0.63 | 7.23E-01 | Sost | 0.63 | 6.64E-01 | Egf | 0.68 | 5.10E-01 | Ifna1 | 0.74 | 8.17E-01 | Fgf4 | 0.75 | 9.09E-01 |
| Rasal1 | 0.63 | 5.38E-01 | Ets2 | 0.63 | 8.46E-03 | Il1r2 | 0.68 | 5.26E-01 | Il6 | 0.74 | 8.21E-01 | Tlr2 | 0.75 | 6.00E-01 |
| Cacna1d | 0.63 | 2.22E-01 | Sos1 | 0.62 | 3.95E-03 | Klf4 | 0.67 | 4.09E-02 | Prlr | 0.74 | 8.21E-01 | Fasl | 0.75 | 8.56E-01 |
| Stat4 | 0.63 | 2.85E-01 | Bnip3 | 0.62 | 1.40E-01 | Cacna1d | 0.67 | 2.32E-02 | Rasgrp1 | 0.73 | 8.92E-01 | Pla2g2a | 0.75 | 6.94E-01 |
| Ccnb3 | 0.62 | 7.29E-01 | Osm | 0.62 | 1.14E-01 | Bnip3 | 0.67 | 9.43E-02 | Il5ra | 0.73 | 8.37E-01 | Il2rb | 0.75 | 4.91E-01 |
| Il4 | 0.62 | 6.74E-01 | Pla2g5 | 0.61 | 3.53E-01 | Il22ra2 | 0.66 | 1.63E-01 | Tnf | 0.73 | 8.02E-01 | Creb3l1 | 0.73 | 5.18E-01 |
| Lef1 | 0.61 | 7.04E-01 | Fgfr4 | 0.61 | 1.01E-01 | Pla2g4e | 0.66 | 6.51E-01 | Map2k6 | 0.73 | 5.75E-01 | Tnfsf10 | 0.73 | 4.91E-01 |
| Cacna1e | 0.61 | 3.22E-01 | Il11ra1 | 0.61 | 1.15E-01 | Tshr | 0.66 | 5.10E-01 | Fgf6 | 0.72 | 8.04E-01 | Fzd2 | 0.73 | 3.48E-01 |
| Mmp9 | 0.60 | 3.61E-01 | Acvr2a | 0.61 | 1.74E-02 | Cacna1e | 0.66 | 4.86E-02 | Cxcl2 | 0.71 | 8.17E-01 | Il23a | 0.73 | 6.94E-01 |
| Rasgrp1 | 0.60 | 7.60E-01 | Il22ra2 | 0.60 | 1.75E-01 | Hspb1 | 0.65 | 2.37E-01 | Gngt1 | 0.71 | 8.04E-01 | Gadd45g | 0.72 | 6.94E-01 |
| Pla2g3 | 0.60 | 8.62E-01 | Nr4a1 | 0.60 | 2.13E-01 | Acvr2a | 0.64 | 5.47E-03 | Il1b | 0.71 | 8.02E-01 | Gzmb | 0.72 | 6.94E-01 |
| Calml3 | 0.60 | 9.66E-01 | Comp | 0.60 | 3.13E-01 | Ppargc1a | 0.64 | 2.24E-01 | Pax5 | 0.71 | 8.02E-01 | Il3 | 0.72 | 8.68E-01 |
| Itga8 | 0.60 | 2.03E-01 | Wnt7b | 0.58 | 5.35E-01 | Fgf13 | 0.63 | 3.82E-02 | Fgf12 | 0.69 | 9.32E-01 | Epor | 0.72 | 3.35E-01 |
| Wnt2 | 0.60 | 2.40E-01 | Ptpn5 | 0.57 | 7.72E-02 | Pla2g3 | 0.63 | 4.87E-01 | Cd19 | 0.69 | 5.75E-01 | Itgb6 | 0.72 | 4.91E-01 |
| Bmp7 | 0.59 | 2.41E-01 | Pparg | 0.56 | 2.69E-01 | Ddb2 | 0.62 | 4.94E-03 | Ntrk2 | 0.69 | 8.17E-01 | Col3a1 | 0.71 | 6.16E-01 |
| Wnt10a | 0.59 | 6.32E-01 | Rps6ka6 | 0.55 | 7.23E-02 | Zbtb16 | 0.61 | 3.40E-01 | Suv39h2 | 0.69 | 8.02E-01 | Il1b | 0.70 | 7.48E-01 |
| Flt3 | 0.58 | 3.61E-01 | Cntfr | 0.55 | 5.98E-02 | Vegfb | 0.61 | 2.89E-02 | Ntf3 | 0.68 | 8.02E-01 | Rxrg | 0.70 | 8.68E-01 |
| Gng4 | 0.57 | 4.97E-01 | Jun | 0.54 | 6.94E-03 | Sfrp4 | 0.61 | 3.59E-01 | Cntfr | 0.68 | 8.02E-01 | Sgk2 | 0.69 | 9.41E-01 |
| Hspb1 | 0.57 | 6.01E-01 | Tshr | 0.53 | 4.46E-01 | Efna3 | 0.61 | 4.48E-01 | Ibsp | 0.68 | 8.02E-01 | Inhbb | 0.69 | 8.42E-01 |
| Pla2g4e | 0.53 | 8.78E-01 | Acvr1c | 0.53 | 2.92E-01 | Prkar2b | 0.60 | 2.93E-01 | Tnc | 0.67 | 8.02E-01 | Mycn | 0.69 | 3.57E-01 |
| Pck1 | 0.53 | 9.30E-01 | Calml3 | 0.53 | 8.12E-01 | Pla2g10 | 0.60 | 4.11E-01 | Nr4a3 | 0.67 | 8.02E-01 | Il6 | 0.68 | 7.48E-01 |
| Wnt7b | 0.52 | 7.66E-01 | Ddb2 | 0.53 | 5.82E-03 | Ccr7 | 0.58 | 1.64E-03 | Tshr | 0.66 | 9.04E-01 | Ccr7 | 0.68 | 4.30E-01 |
| Ccr7 | 0.50 | 2.66E-02 | Figf | 0.52 | 1.96E-03 | Lep | 0.57 | 1.90E-01 | Comp | 0.66 | 8.04E-01 | Hhex | 0.66 | 3.86E-01 |
| Ifng | 0.50 | 3.61E-01 | Zbtb16 | 0.52 | 3.57E-01 | Calml3 | 0.56 | 7.10E-01 | Klf4 | 0.64 | 8.02E-01 | Nog | 0.65 | 9.04E-01 |
| Il1r2 | 0.49 | 7.60E-01 | Pax5 | 0.51 | 6.31E-03 | Wnt7b | 0.55 | 3.39E-01 | Uty | 0.63 | 8.02E-01 | Ifna1 | 0.64 | 7.48E-01 |
| Rps6ka6 | 0.49 | 1.35E-01 | Vegfb | 0.51 | 2.24E-02 | Hspa1a | 0.54 | 2.58E-01 | Prkar2b | 0.62 | 8.08E-01 | Zic2 | 0.64 | 7.38E-01 |
| Figf | 0.48 | 6.97E-03 | Sfrp4 | 0.48 | 2.88E-01 | Ntrk2 | 0.54 | 1.42E-01 | Osm | 0.61 | 8.02E-01 | Flt3 | 0.64 | 6.20E-01 |
| Fgf13 | 0.47 | 6.72E-01 | Pdgfra | 0.47 | 2.20E-03 | Fgfr4 | 0.53 | 5.39E-03 | Pparg | 0.60 | 8.02E-01 | Cd19 | 0.62 | 2.34E-01 |
| Efna3 | 0.47 | 7.36E-01 | Il12a | 0.46 | 2.54E-04 | Rps6ka6 | 0.52 | 6.54E-03 | Pla2g5 | 0.60 | 8.02E-01 | Prl | 0.62 | 5.18E-01 |
| Fgfr4 | 0.46 | 8.93E-02 | Ppargc1a | 0.46 | 9.06E-02 | Cacng6 | 0.50 | 3.28E-01 | Fgf21 | 0.56 | 8.02E-01 | Pax5 | 0.61 | 3.57E-01 |
| Ptpn5 | 0.44 | 6.04E-02 | Ntrk2 | 0.42 | 1.14E-01 | Figf | 0.50 | 3.94E-05 | Ppargc1a | 0.51 | 8.02E-01 | Pla2g4e | 0.61 | 9.43E-01 |
| Il12a | 0.42 | 1.09E-03 | Klf4 | 0.42 | 1.96E-03 | Ptpn5 | 0.50 | 3.67E-03 | Sost | 0.50 | 8.17E-01 | Mmp3 | 0.58 | 5.82E-01 |

| | | | | | | | | | | | | | | |
|---|---|---|---|---|---|---|---|---|---|---|---|---|---|---|
| Pdgfra | 0.42 | 7.79E-03 | Cacng1 | 0.41 | 6.10E-01 | Il12a | 0.44 | 2.62E-06 | Fos | 0.45 | 8.02E-01 | Wnt7a | 0.57 | 2.13E-02 |
| Pla2g10 | 0.39 | 6.39E-01 | Cd19 | 0.40 | 6.90E-05 | Pdgfra | 0.44 | 5.15E-05 | Il20ra | 0.44 | 8.02E-01 | Lif | 0.57 | 2.13E-02 |
| Igfbp3 | 0.38 | 4.96E-02 | Prkar2b | 0.39 | 1.12E-01 | Pax5 | 0.41 | 3.94E-05 | Inhba | 0.41 | 8.17E-01 | Tnc | 0.54 | 4.28E-01 |
| Pax5 | 0.33 | 1.66E-03 | Igfbp3 | 0.37 | 4.67E-03 | Igfbp3 | 0.38 | 3.83E-04 | Nr4a1 | 0.38 | 4.14E-01 | Ppp2r2c | 0.52 | 3.19E-01 |
| Cacng1 | 0.32 | 8.22E-01 | Lep | 0.37 | 8.02E-02 | Cacng1 | 0.36 | 4.08E-01 | Hspa1a | 0.38 | 8.02E-01 | Cxcl5 | 0.49 | 5.22E-02 |
| Cd19 | 0.30 | 7.97E-05 | Hspa1a | 0.28 | 7.72E-02 | Cd19 | 0.35 | 2.67E-07 | Sgk2 | 0.35 | 8.02E-01 | Ifnb1 | 0.45 | 7.92E-01 |
| Cacng6 | 0.28 | 4.31E-01 | Pck1 | 0.14 | 2.42E-01 | Pck1 | 0.27 | 3.30E-01 | Pck1 | 0.17 | 8.02E-01 | Inhba | 0.34 | 7.94E-01 |

| **Supplementary Table 3 \|** Taqman® gene expression assays, Life Technologies | | | | |
|---|---|---|---|---|
| **Probe ID** | **Gene** | **Species** | **Full name** | **Reporter** |
| Mm00446190_m1 | *Il6* | Mouse | Interleukin 6 | FAM |
| Mm04207460_m1 | *Cxcl-1* | Mouse | Chemokine (C-X-C motif) ligand 1, KC | FAM |
| Mm00440295_m1 | *Mmp3* | Mouse | Matrix metalloproteinase 3 | FAM |
| Mm00441818_m1 | *Timp1* | Mouse | TIMP metallopeptidase inhibitor 1 | FAM |
| Mm00494449_m1 | *Cdkn2a* | Mouse | CDK inhibitor 2a, p16 | FAM |
| Mm01303209_m1 | *Cdkn1a* | Mouse | CDK inhibitor 1a, p21 | FAM |
| Mm00438168_m1 | *Cdkn1b* | Mouse | CDK inhibitor 1b, p27 | FAM |
| Mm00439620_m1 | *Il1a* | Mouse | Interleukin 1a | FAM |
| Mm00439560_m1 | *Igf1* | Mouse | Insulin-like growth factor 1 | FAM |
| Mm00437762_m1 | *B2m* | Mouse | Beta-2-Microglobulin | FAM |
| 4352339e-1207040 | *Gapdh* | Mouse | Glyceraldehyde 3 phosphate dehydrogenase | VIC |
| Mm01309913_m1 | *Tmem199* | Mouse | Transmembrane protein 199 | FAM |
| Hs00985639_m1 | *IL6* | Human | Interleukin 6 | FAM |
| Hs00174103_m1 | *IL8* | Human | Interleukin 8 | FAM |
| Hs00968305_m1 | *MMP3* | Human | Matrix metalloproteinase 3 | FAM |
| Hs00171558_m1 | *TIMP1* | Human | TIMP metallopeptidase inhibitor 1 | FAM |
| Hs00355782_m1 | *CDKN1A* | Human | CDK inhibitor 1a, p21 | FAM |
| Hs00923894_m1 | *CDKN2A* | Human | CDK inhibitor 2a, p16 | FAM |
| Hs99999907_m1 | *B2M* | Human | Beta-2-Microglobulin | FAM |
| 4310884E | *GAPDH* | Human | Glyceraldehyde 3 phosphate dehydrogenase | VIC |

| **Supplementary Table 4 \|** Antibodies | | | | |
|---|---|---|---|---|
| **Antibody Target** | **Source** | **ID** | **Species** | **Dilution** |
| p21 | Dako | M7202 | Mouse | 1/50 |
| p21 | Abcam | ab188224 | Rabbit | 1/4000 |
| γH2Ax (S139) | Cell Signaling | 9718S | Rabbit | 1/100 |
| α Smooth muscle actin | Sigma | A2547 | Mouse | 1/5000 |
| EpCAM | Abcam | ab221552 | Rabbit | 1/200 |
| Endomucin (V7C7) | Santa Cruz | sc65495 | Rat | 1/1000 |
| CD4 | Ebioscience | 14-9766 | Rat | 1/250 |
| CD8 | Ebioscience | 14-0808 | Rat | 1/250 |
| Ki67 | Abcam | ab16667 | Rabbit | 1/300 |
| MCL-1 | Cell Signaling | 94296 | Rabbit | 1/80 |
| Firefly luciferase | Abcam | ab181640 | Goat | 1/100 |
| Lamin A/C | Abcam | ab108595 | Rabbit | 1/1000 |
| Mouse IgG-488 | Invitrogen | A11001 | Goat | 1/1000 |
| Mouse IgG-555 | Invitrogen | A21127 | Goat | 1/1000 |
| Rat IgG-488 | Invitrogen | A11006 | Goat | 1/1000 |
| Rabbit IgG-488 | Invitrogen | A11008 | Goat | 1/1000 |
| IgG, immunoglobulin G; γH2Ax, phosphorylated histone H2AX | | | | |
